# Evolutionary Dynamics of the Proanthocyanidin Biosynthesis Gene *LAR*

**DOI:** 10.1101/2025.08.06.668978

**Authors:** Maria F. Marin-Recinos, Boas Pucker

## Abstract

**Background:** Leucoanthocyanidin reductase (LAR) is a key enzyme in proanthocyanidin (PAs) biosynthesis, catalyzing the conversion of leucoanthocyanidins to catechins. While early steps in the flavonoid pathway are broadly conserved across plant lineages, increasing evidence demonstrates lineage-specific evolutionary trajectories and functional diversification in its terminal branches, particularly in the case of LAR. To explore the evolutionary dynamics and functional divergence of LAR genes, we conducted a large-scale comparative and phylogenetic analyses across major plant clades.

**Results:** Phylogenetic analysis revealed multiple independent duplication events and lineage-specific expansions of *LAR* lineages, particularly among dicots and gymnosperms. In dicots, *LAR1* and *LAR2* were differentially retained and diversified, whereas gymnosperm LAR homologs formed early-diverging clades, suggesting an ancient duplication and potential neofunctionalization. Coexpression analyses across species and tissues indicate paralog-specific expression patterns. Sequence analysis identified both conserved and clade-specific protein domains, supporting functional divergence. Promoter analyses showed differences in transcription factor binding site composition between *LAR1* and *LAR2*, pointing to regulatory sub- or neofunctionalization. Lastly, synteny analyses supports the potential absence of *LAR* in multiple *Brassicacea* genomes.

**Conclusions:** LAR shows evidence of evolutionary diversification, shaped by both coding and regulatory changes. These patterns of diversification help explain variation in flavonoid profiles in gymnosperms and angiosperms. Understanding the evolutionary dynamics of LAR not only deepens our knowledge of metabolic pathway evolution but also provides insights relevant to the breeding and metabolic engineering of plant traits related to pigmentation, stress resilience, and nutritional quality.

## INTRODUCTION

Proanthocyanidins (PAs), also referred to as condensed tannins, constitute a group of polyphenolic compounds within the flavonoid biosynthesis pathway. As secondary metabolites, PAs play an important role in plant defense mechanisms against insect herbivores and microbial pathogens. Their contribution to herbivore deterrence is largely attributed to the astringent and bitter taste they confer to various plant tissues, which reduces palatability and impairs digestibility in insect pests [1–3]. In addition, PAs exhibit antimicrobial activity against pathogens, including bacteria [4–6] and fungi [7]. Historically, PAs have been utilized as tanning agents for leather preservation and in the regulation of different flavor qualities and profiles in beverages such as wine, teas, and fruit juices [8, 9]. Furthermore, their antioxidant properties have been reported for their potential health benefits in both human and ruminant animal consumption [10]. In addition to their role in plant defense, PAs have been extensively studied for their impact on seed coat pigmentation [11, 12], influencing seed dispersal [13] and dormancy mechanisms [14], thereby contributing to plant reproductive efficacy [15, 16].

PA synthesis represents one branch of the flavonoid biosynthesis pathway, a process extensively elucidated across a wide range of plant species and notably conserved among diverse taxa [17–19]. The synthesis of PAs and anthocyanins, as competing branches of the pathway, share common precursors which involve a series of enzymatic reactions. The key enzymes involved in PA synthesis start with the enzyme dihydroflavonol 4-reductase (DFR) catalyzing the production of leucoanthocyanidins which are then processed by leucoanthocyanidin reductase (LAR) for the conversion into 2,3-*trans*-flavan-3-ols, commonly known as the monomer 2,3-*trans*-(+)-catechin. In a parallel route, anthocyanidin synthase (ANS) converts leucoanthocyanidins to anthocyanidins, which can be reduced by anthocyanidin reductase (ANR) to form 2,3-*cis*-flavan-3-ols, also known as the monomer 2,3-*cis*-(-)-epicatechin. These monomeric units then polymerize to form PA oligomers (Figure 1).

**Figure 1.**
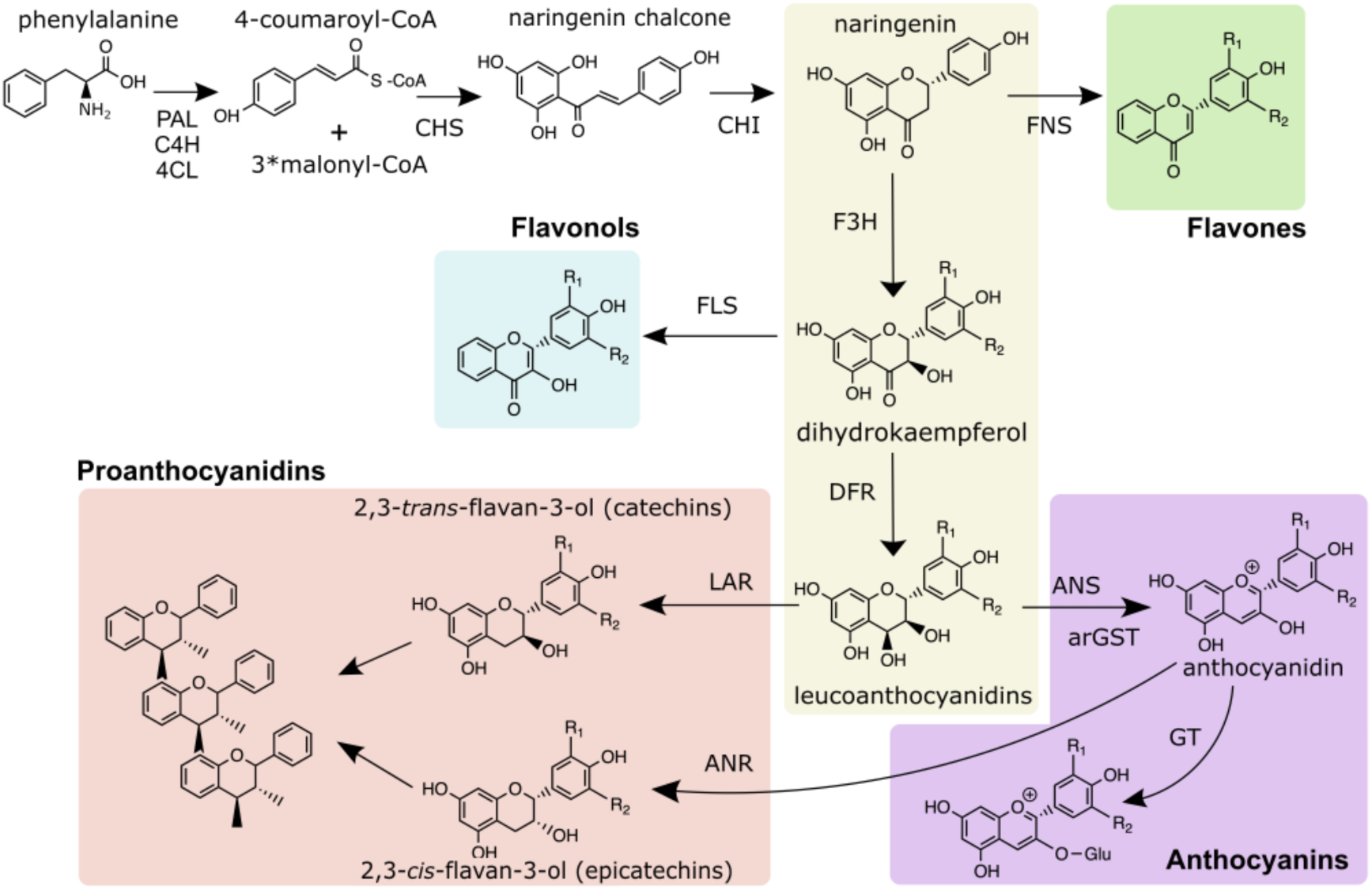
Schematic representation of the general flavonoid biosynthesis pathway. Enzyme names are abbreviated as follows: PAL - phenylalanine ammonia-lyase, C4H - cinnamic acid 4-hydroxylase, 4CL - 4-coumarate-CoA ligase, CHS - chalcone synthase, CHI - chalcone isomerase, FNS - flavone synthase, F3H - flavanone 3-hydroxylase, FLS - flavonol synthase, DFR - dihydroflavonol 4-reductase, ANS - anthocyanidin synthase, arGST – anthocyanin-related glutathione S-transferase, GT - UDP-dependent anthocyanidin- 3-O-glucosyltransferase, ANR - anthocyanidin reductase.

LAR and ANR play a central role in defining PA composition by synthesizing the flavan-3-ol monomers that act as starter and extension units. Canonically, the primary constituents of these polymers are (+)-catechin, produced by LAR, and (-)-epicatechin, synthesized by ANR. These two monomers serve as the basic building blocks in the formation of proanthocyanidin polymers. However, recent studies in *Medicago truncatula* have reported that the function of LAR extends beyond its traditional role in the synthesis of (+)-catechins [20, 21]. Specifically, LAR has been shown to be responsible for converting 4β-(S-cysteinyl) epicatechin back to epicatechin, a reaction that was previously attributed exclusively to ANR [20, 21]. This dual LAR activity may explain the observed increase in catechin and epicatechin levels under overexpression of *LAR*, indicating that LAR is crucial for regulating both monomer types that serve as starter and extension units in the polymerization process of PAs [20, 21]. Furthermore, Jun *et al.* demonstrated that leucoanthocyanidin dioxygenase (LDOX) in *M. truncatula* plays a role in the formation of epicatechin starter units from catechin, contrasting with the function of ANS in the conversion of leucocyanidin to cyanidin [21]. Meanwhile, ANR is involved in the synthesis of the extension epicatechin units [21].

In addition to *M. truncatula*, *LAR* genes have been characterized in multiple economically important plants such as *Vitis vinifera* [22], *Theobroma cacao* [23], *Gossypium hirsutum* [24], *Malus domestica* [25], *Pyrus communis* [26], and *Camellia sinensis* [27], all which belong to a diverse range of plant families and orders. Interestingly, despite the widespread presence of *LAR* genes in many plants, it has been known for years that the model plant *Arabidopsis thaliana* does not harbor a *LAR* gene [19, 28]. However, *A. thaliana* is still capable of synthesizing proanthocyanidins, which provide a dark color to its seed coat [12]. Knowledge obtained from studying the *transparent testa* (*tt*) and *tannin-deficient seed* (*tds*) mutants collection in *A. thaliana* have been essential in understanding the synthesis, regulation, and transport mechanisms involved in the PA production, even in the absence of a functional *LAR* gene [12, 19, 29–32]. The regulation of flavonoid biosynthesis, including the PA branch, is controlled by a well-characterized complex of transcription factors (TFs) known as the MBW complex, consisting of MYB, basic helix-loop-helix (bHLH), and WD40 proteins [33]. In *A. thaliana*, key TFs such as TT2 (R2R3-MYB), TT8 (bHLH), and TTG1 (WD40) regulate the expression of *BANYULS (BAN)* which encodes for ANR involved in the PA production in seeds [12, 34, 35].

Given the wide-spread occurrence of LAR in various plant lineages and its absence in *A. thaliana*, understanding the evolutionary trajectory of LAR genes can provide valuable insights into their functional diversification and potential compensatory mechanisms in species lacking LAR. Here, we investigate the evolutionary dynamics of *LAR* through a systematic analysis of its presence/absence across land plant lineages. Our findings reveal a deep duplication event at the base of the evolutionary split between gymnosperms and angiosperms that has resulted in many plant species harboring two *LAR* copies, suggesting potential subfunctionalization or neofunctionalization within these paralogs. However, we also identified entire plant lineages without *LAR*. Most noticeable, *LAR* appears to be largely missing within the *Brassicaceae* family. Additionally, we explored the functional divergence of the two *LAR* lineages in plants with multiple copies, aiming to understand how gene duplication has influenced the evolution and PA biosynthesis.

## RESULTS

### Phylogenetic Analysis of Leucoanthocyanidin Reductase (LAR)

This study explored the phylogenetic relationships of LAR genes across a wide range of plant taxa. In total, 287 LAR, 268 DFR, and 112 ANR sequences were retained for alignment and phylogenetic analysis. Four primary clades were identified in the LAR portion of the tree, corresponding to the major plant lineages: gymnosperms, monocotyledons, magnoliids, and dicotyledons (Figure 2). Within both the gymnosperm and dicotyledon clades, two distinct subclades were identified. These will be referred to here as LARI and LARII for those in the gymnosperm clade (shown in orange and green in Figure 2, respectively) and LAR1 and LAR2 for those in the dicotyledon clade (shown in red and blue in Figure 2, respectively). These distinctions are necessary to highlight clear phylogenetic divergence within each lineage and to avoid confusion in downstream analysis.

Interestingly, the presence of separate *LAR* gene copies from the same species in both subclades of gymnosperms (LARI and LARII), as well as in both subclades of dicotyledons (LAR1 and LAR2) implies that an ancient gene duplication event occurred prior to the diversification of each of these groups (Figure 2). This deep duplication appears to have given rise to two major evolutionary lineages of *LAR* genes in both gymnosperms and dicots. Following this deep duplication, additional lineage-specific duplication events occurred at the family, genus, or species level independently within each clade. For example, in gymnosperms, species such as *Thuja plicata, Picea sitchensis,* and *Cryptomeria japonica* present multiple LAR copies distributed across the subclades LARI and LARII. In dicots, the LAR1 subclade represents a particularly large and diverse collection of gene copies. Many species exhibit multiple paralogs within this subclade, pointing to repeated gene duplication events within specific lineages. Families such as *Theaceae, Fabaceae, Rosaceae,* and *Malvaceae* are especially well represented in LAR1, often with two or more copies per species. The LAR2 subclade, while smaller, also includes multiple gene copies of species present in LAR1. Notably, species such as *Bretschneidera sinensis* (*Akaniaceae*) and *Carica papaya* (*Caricaceae*) both belonging to the *Brassicales* order, contain LAR gene copies only in LAR1, suggesting a potential absence of the LAR2 copy in these lineages. This pattern further supports the idea that, following the initial ancient duplication, LAR1 and LAR2 have undergone independent expansions in different lineages, leading to a potential clade-specific diversification of the LAR genes.

**Figure 2.**
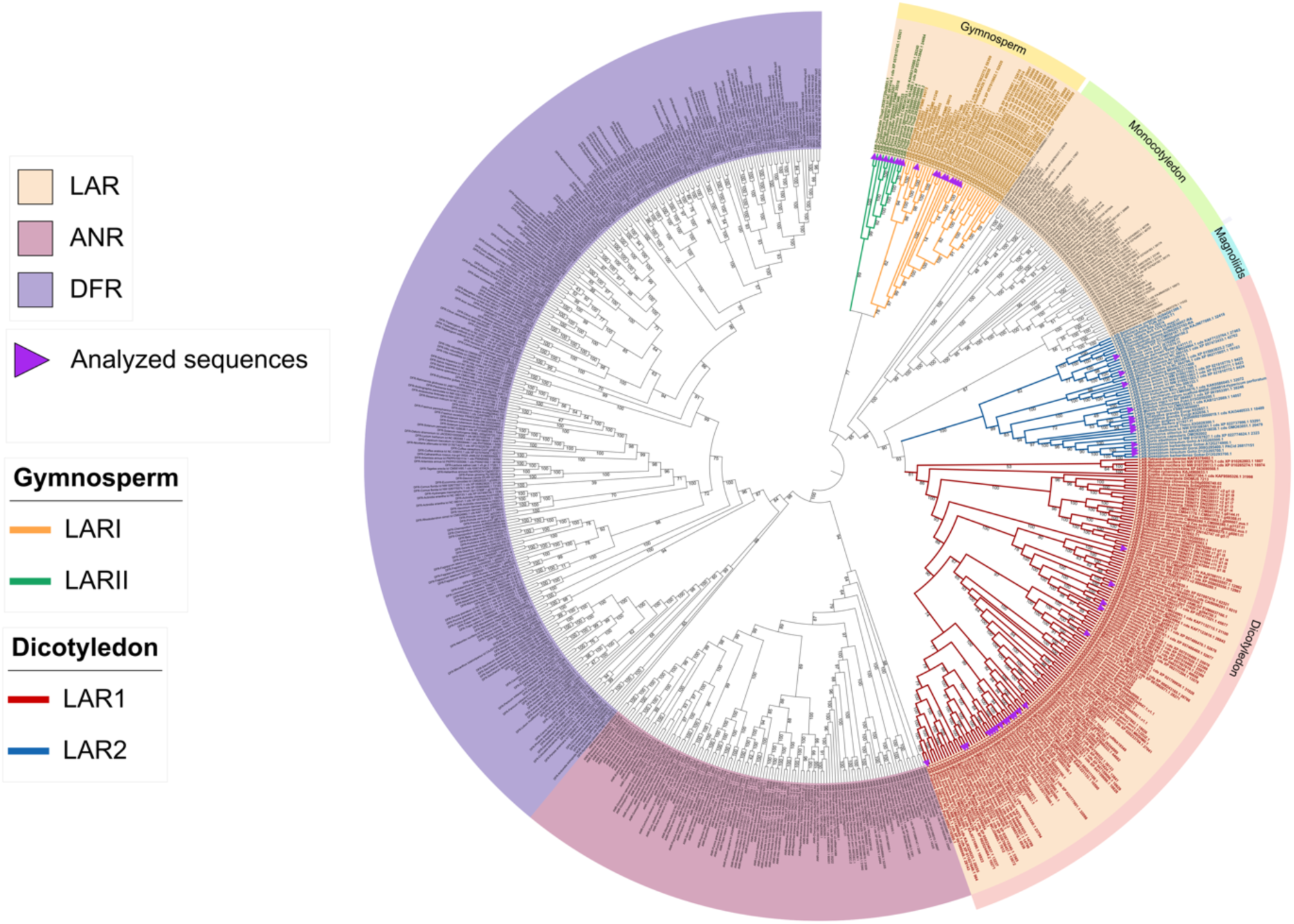
Phylogenetic analysis of LAR, DFR, and ANR sequences in different plant species. Sequences were collected via KIPEs3 results, codon-based alignments were generated with MACSE [36], and the phylogeny was inferred with IQ-TREE2 [37, 38] based on maximum likelihood. Distinct clades of LARI (orange) and LARII (green) in the Gymnosperm group as well as LAR1 (red) and LAR2 (blue) in the Dicotyledon group are highlighted. The purple arrows indicate the sequences used for the synteny and gene expression analysis.

### Leucoanthocyanidin reductase duplication event and functional divergence

The phylogenetic analysis (Figure 2) revealed the presence of multiple *LAR* gene copies in several species within both gymnosperms and dicotyledons. This pattern points to an ancestral gene duplication event, followed by divergence within each lineage. Not only these gene copies are phylogenetically distinct, but also they are not located near each other in the genome. This suggests that they did not arise from recent tandem duplication but rather from older duplication events followed by independent evolutionary paths. Over time, these copies have likely diverged in sequence and regulation, which may reflect differences in function or tissue-specific roles. To explore this potential functional divergence, gene expression patterns of the different LAR paralogs were examined across multiple tissues in representative species presented in the phylogenetic tree. For gymnosperms, *Taxus chinensis*, *Ginkgo biloba*, *Pinus pinaster, Picea sitchensis,* and *Thuja plicata* were analyzed (Figure 3, Additional file 1) and for dicotyledons, *Vitis vinifera*, *Theobroma cacao*, *Moringa oleifera, Gossypium hirsutum, Actinidia chinensis, Eucalyptus grandis, Populus trichocarpa, Carya illinoinensis,* and *Morella rubra* (Figure 4; Additional file 1). *T. chinensis*, showed a significant difference in expression between *TchLARI* (KAH9310580.1) and *TchLARII* (KAH9295439.1). *TchLARII* showed consistently higher expression across all the sampled tissues (Figure 3A). A similar pattern was observed in *G. biloba,* where *GbiLARII* (GBI00018207) had a significantly higher overall expression than *GbiLARI* (GBI00017466) with tissue-specific expression patterns showing stronger *GbiLARII* expression in seedling stem tissues (p-value = 0.0004) and root tissues (p-value = 0.002), and a stronger *GbiLARI* expression in stem tissues (p-value <0.0001) (Figure 3B). In *P. pinaster,* a significant overall expression difference was also detected between *PpiLAR1* (PPI00016665) and *PpiLARII* (PPI00057446), with *PpiLARII* being more highly expressed in needles (p-value <0.0001), shoots (p-value <0.0001), and stem tissues (p-value <0.0001) (Figure 3C).

**Figure 3.**
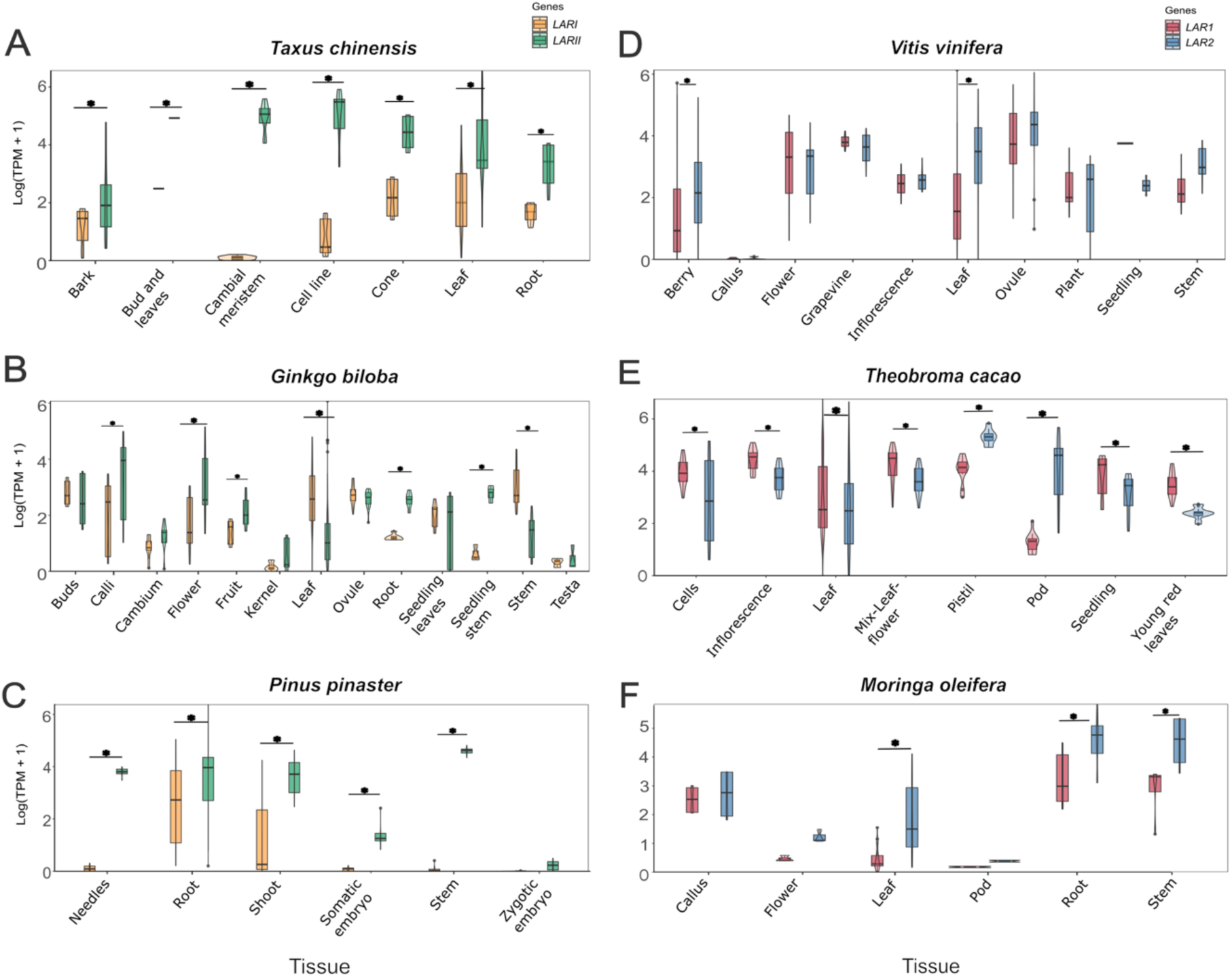
Comparison of the expression of *LARI* (orange) and *LARII* (green) genes in the species *Taxus chinensis* (A), *Ginkgo biloba* (B) and *Pinus pinaster* (C), and *LAR1* (red) and *LAR2* (blue) in the species *Vitis vinifera* (D), *Theobroma cacao* (E), and *Moringa oleifera* (F) genes across different plant tissues. An asterisk above the violin plot indicates a statistically significant difference (p < 0.05) between LARs in that specific tissue, based on a linear mixed-effects model followed by estimated marginal means analysis. The complete results of the LMM and EMMs are provided in Additional file 1.

In dicots, clear expression differences between paralogs were also identified. In *V. vinifera, VviLAR2* (VIT_217s0000g04150.2) showed significantly higher expression than *VviLAR1* (VIT_201s0011g02960.1) in leaf tissues (p-value <0.0001) and berry tissues (p-value <0.0001) (Figure 3D). In *T. cacao*, the gene expression of *TcaLAR1* (Thecc.02G361400.1) was dominant in cells, inflorescence, leaf, seedlings, young red leaves, and mix of leaf and flower tissues while *TcaLAR2* (Thecc.03G028300.1) presented higher expression in pod (p-value <.0001) and pistil tissues (p-value <0.0001) (Figure 3E). Similarly, in *M. oleifera,* a significant overall expression difference was detected between *MolLAR1* (g8395.t1) and *MolLAR2* (g11921.t1), with *MolLAR2* exhibiting higher expression in roots (p-value = 0.002), leaf (p-value <0.0001), and stem tissues (p-value = 0.005) (Figure 3F).

When comparing the 36 sequences of LARI and the 11 sequences of LARII in gymnosperms, both groups showed strong conservation at many positions, numbered according to the *Theobroma cacao* LAR1 reference sequence (Figure 4A, Additional file 2). Particularly interesting are positions with contrasting differences between LARI and LARII (Figure 4A). In dicotyledonous species, 106 sequences of LAR1 and 43 sequences of LAR2 were analyzed, revealing comparable conservation with notable divergences at several key residues (Figure 4A, Additional file 2). Conserved regions suggest key functional or structural roles. Amino acid substitutions at five specific positions in gymnosperms (18, 117, 126, 161, 167) and seven in dicots (15, 117, 157, 161, 167, 168, 255) clearly differentiate the gymnosperm LARs (LARI, LARII) from their dicot counterparts (LAR1, LAR2). At position 15, a consistent presence of alanine (A) was found in LARI, LARII, and LAR1, while a substitution to serine (S) was observed in LAR2. At position 18, tyrosine (Y) was exclusive to LARI, while phenylalanine (F) was dominant in LARII and dicots (Figure 4B). At position 117, valine (V) was retained in both LARII and LAR1, while isoleucine (I) was found in LARI and LAR2. A unique alanine (A) in LARII was observed in position 126, substituting glycine (G) in LARI and dicots. The ICCN motif region centered on position 157 showed a particular divergence in dicots, while serine (S) is highly conserved in LAR1, alanine (A) is more conserved in LAR2. Similar results can be observed at position 167, serine (S) was conserved in LARI and LAR1, while threonine (T) appeared in LARII, and alanine (A) was found in LAR2. Given the proximity of this position to the active site and the polarity differences, this variation was interpreted as functionally relevant (Figure 4C). An additional substitution was identified at position 168, located within a conserved motif, remained glutamate (E) in LARI, LARII, and LAR1, but was substituted with aspartate (D) in LAR2, reducing the side chain length and potentially altering charge distribution or hydrogen bonding patterns. Finally, position 255, while this has not been reported as part of any LAR-specific motif, showed a predominant conservation of asparagine (N) in all groups except LAR2, which presented a substitution to methionine (M). This polar-to-nonpolar shift, occurring near the active site, was interpreted as likely contributing to a reduction in the catalytic efficiency of LAR2 as reported by the gene expression in Figure 3E for *T. cacao*. The proximity of several variable positions to conserved motifs and the active site further supports their relevance in the functional diversification of LAR enzymes.

**Figure 4.**
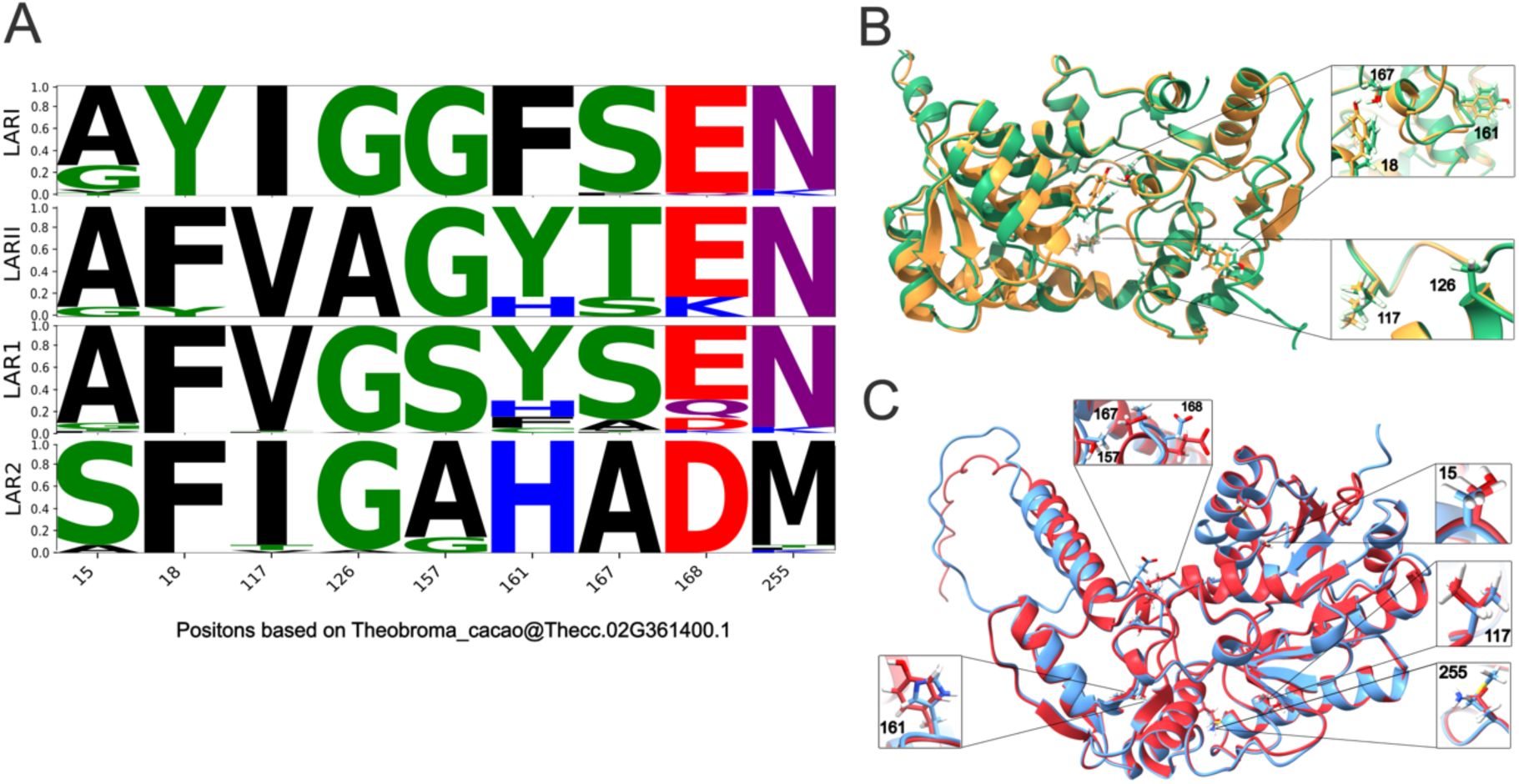
Functional divergence in LAR proteins. (A) Comparative sequence logo showing conservation and variability across LARI and LARII sequences in gymnosperm species, and LAR1 and LAR2 sequences in dicotyledon species. The X-axis represents the position based on *Theobroma cacao* LAR1 reference sequence, while the Y-axis indicates the frequency in percentage, reflecting sequence conservation at each position. Individual amino acids are color-coded based on their chemical properties: nonpolar/hydrophobic residues (black), polar amino acids with hydroxyl groups (green), polar amino acids with amide residues (purple), positively charged residues (blue), and negatively charged residues (red). The height of each letter within a column corresponds to its relative frequency at that position with taller letters indicating more conserved regions and shorter ones reflecting variability. (B) Structural model of LARI (orange) and LARII (green) of the gymnosperm species *Taxus chinensis*, highlighting five positions with amino acid substitution near the active site of the protein. (C) Structural model of LAR1 (red) and LAR2 (blue) of the dicotyledon species *Theobroma cacao*, highlighting seven positions with amino acid substitution near the active site of the protein.

The combined analysis of gene expression patterns and protein sequence conservation indicates a clear functional differentiation between the *LAR* gene groups. *LAR1* and *LAR2* in dicots, as well as *LARI* and *LARII* in gymnosperms, exhibit distinct tissue-specific expression profiles. These expression patterns are supported at the molecular level by systematic amino acid substitutions revealed through the comparative analysis of LAR sequences. These substitutions differentiate dicots and gymnosperms LARs, as well as within each LAR lineage. Mutations such as Y18F in gymnosperms LARII, and S157A, and E168D both occurring in dicots LAR2, present particular interest due to occurring within conserved motifs.

### Expression analysis among *LAR1* and *LAR2* genes

Given the distinct expression patterns and systematic amino acid differences previously identified between LAR1 and LAR2, a coexpression analysis was conducted to further investigate their potential functional divergence and relationships with other metabolic pathways. The results are presented in two tables, in which each *LAR* gene is depicted along with a list of coexpressed genes. For each gene, the Spearman correlation coefficient and gene expression value are provided (See Additional file 3). In *V. vinifera, VviLAR1* (VIT_201s0011g02960.1) was notably coexpressed with the genes; VIT_206s0004g08150.1 (r = 0.710) encoding trans-cinnamate 4-monooxygenase, a key enzyme in the biosynthesis of flavonoids and lignin, and VIT_213s0067g02870.2 (r = 0.657) encoding the enzyme chalcone-flavone isomerase, involved in the flavonoid biosynthesis. In *T. cacao* the analysis identified strong correlations between *TcaLAR1* (Thecc.02G361400.1) and a diverse set of high expressed protein-coding genes involved in stress responses such as proline-rich protein-1 (Thecc.04G253900.1, r = 0.666) and the bax inhibitor-1 (Thecc.04G271000.1, r = 0.683). Additionally, genes involved in signaling pathways like Calmodulin7 (Thecc.09G265600.2, r = 0.763) which plays an important role in seedling development, and vesicle trafficking like ADP-ribosylation factor (Thecc.01G320100.1, r = 0.708) and ESCRT-related protein CHMP1B (Thecc.06G106700.1, r = 0.695). *MolLAR1* (g8395.t1) in *M. oleifera*, presented strong correlation with genes encoding metallothionein-like protein 1 (g23076.t1, r = 0.702), copper transport protein ATX1 (g22635.t1, r = 0.701), and Profilin2 (g19932.t1, r = 0.682).

Regarding the genes coexpressed with *LAR2*, in *V. vinifera* the gene expression of *VviLAR2* (VIT_217s0000g04150.2) showed stronger correlation with an isoform of the same gene (VIT_217s0000g04150.7, r = 0.741), followed by genes encoding sigma intracellular receptor 2 (VIT_212s0059g02290.1, r = 0.664), sucrose transport protein (VIT_218s0076g00250.1, r = 0.658), and patatin-like protein 2 (VIT_218s0001g10830.1, r = 0.651). In *T.cacao*, *TcaLAR2* (Thecc.03G028300.1) coexpressed strongly with the genes encoding the protein SPIRAL1 (Thecc.02G328900.1, r = 0.681), GTP-binding nuclear protein (Thecc.09G120500.1, r = 0.722), and 60S ribosomal protein (Thecc.01G108600.1, r = 0.707). Lastly, *MolLAR2* (g11921.t1) in *M. oleifera* showed coexpression with genes encoding proteins such as WALLS ARE THIN1 (g3827.t1, r = 0.684), Major latex protein-like 43 (g15381.t1, r = 0.672), and triose-phosphate/phosphate translocator (g12330.t1, r = 0.665).

Coexpression analyses further supported divergence among LAR paralogs, with *TcaLAR1* and *MolLAR1* showing strong positive correlations with genes involved in protein regulation and stress response. Likewise, the coexpression profile of *TcaLAR2* was strongly related to tissue-specific expression, specifically to reproductive development, consistent with its higher expression in pods and flower tissues.

### Regulatory divergence between *LAR1* and *LAR2* gene promoters and identification of transposable elements

To explore the regulatory divergence between the two *LAR* gene copies, promoter sequences from *V. vinifera, T. cacao,* and *M. oleifera* were extracted and analyzed for transcription factor binding motifs and transposable elements (TE) insertions. This species-specific comparison provides additional insights into the functional differentiation of these LAR paralogs and highlights the potential influence of TEs on *LAR* gene regulation, particularly in the context of proanthocyanidin biosynthesis.

In *V. vinifera*, both promoters *pVviLAR1* and *pVviLAR2* included motifs associated with flavonoid biosynthesis gene regulation, notably those bound by MYB transcription factors. *pVviLAR1* exclusively contained ABRELATERD1 motifs and shared with *pVviLAR2* the presence of MYCCONSENSUSAT, MYBCORE, MYB1AT, MYBGAHV, and MYB2CONSENSUSAT. In contrast, *pVviLAR2* uniquely harbored a MYBPZM motif. While both genes showed considerable overlap in motif composition, *pVviLAR2* presented elements in closer proximity to the transcription start site (TSS) and a more varied motif composition, which may reflect a broader regulatory role (Figure 5A). Additionally, TEs annotated with EDTA revealed a Helitron TE insertion in *pVviLAR1* and a Mutator-like TIR transposon in *pVviLAR2*.

In *T. cacao*, both promoter regions displayed a diverse motif landscape sharing elements such as MYB1AT, MYBCORE, MYBPZM, and ABRELATERD1. Notably, *pTcaLAR1* presented a higher number of motifs overall and uniquely included multiple MYCCONSENSUSAT elements, while *pTcaLAR2* was characterized by the presence of MYBGAH (Figure 5B). In *M. oleifera,* the promoter region *pMolLAR1* presented a richer and more dense motif profile compared to *pMolLAR2*. Shared motifs included MYB1AT, MYBCORE, MYCCONSENSUSAT, MYBPZM, and MYBPLANT. In addition, ABRELATERD1 and MYB2CONSENSUSAT were exclusively found in *pMolLAR1.* (Figure 5C). No TE insertions were detected in the promoter regions of either *LAR* gene in *T. cacao* and *M. oleifera*.

Collectively, these findings reveal both conservation and divergence in the regulatory landscapes of *LAR1* and *LAR2* across species. MYB and MYC motifs were consistently found and are likely essential for proanthocyanidin biosynthesis regulation. However, the presence of TEs in the *LAR* promoters of *V. vinifera,* in contrast with their absence in *T. cacao* and *M. oleifera*, point toward species-specific regulatory adaptations, supporting the hypothesis that promoter evolution via motif variation and TE insertions may have contributed to species-specific expression patterns of LAR paralogs.

**Figure 5.**
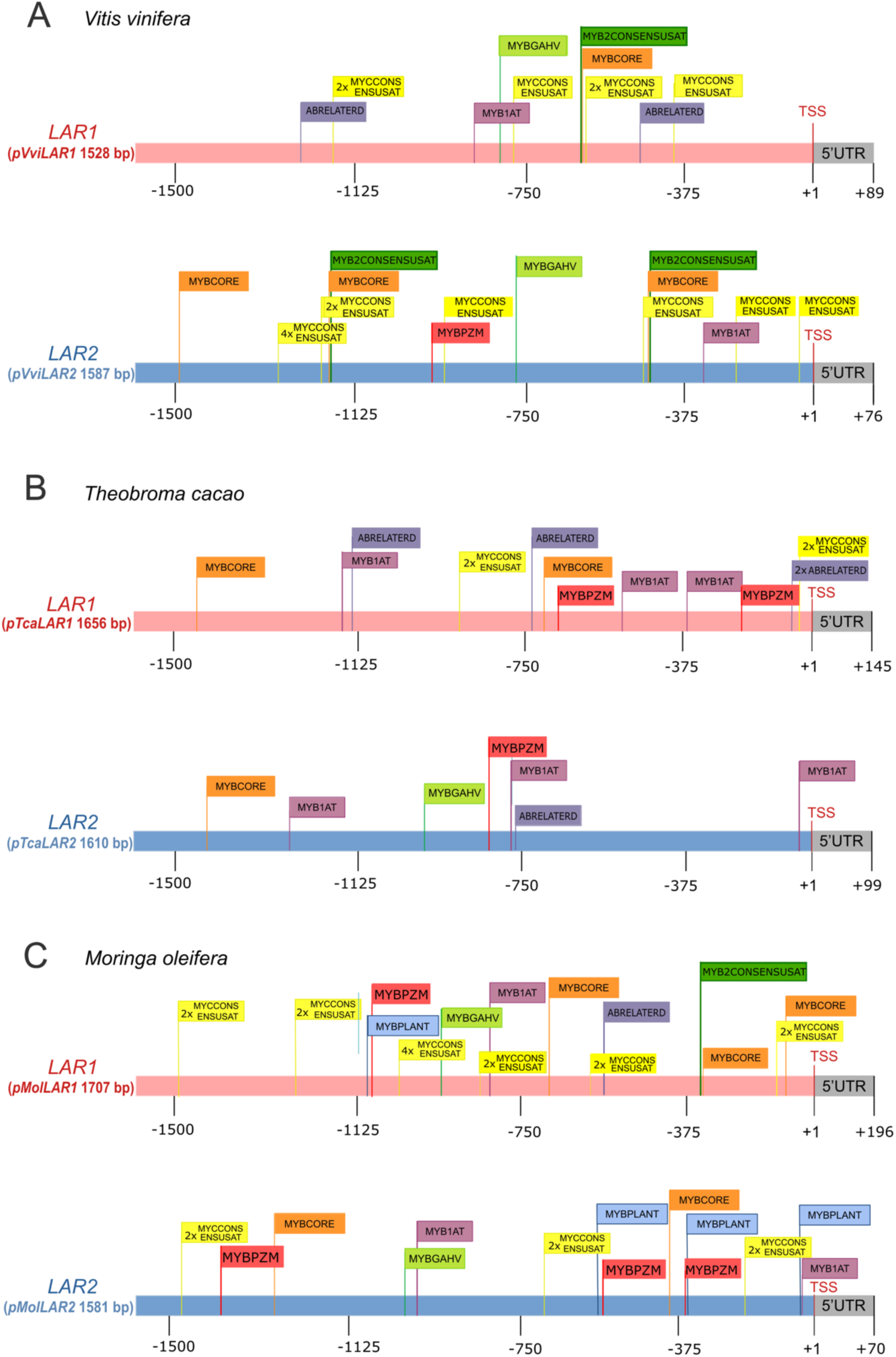
Promoter region constructs with putative cis-acting regulatory elements predicted by the PLACE database [39]. (A) *pVviLAR1* and *pVviLAR2* in *Vitis vinifera.* (*B*) *pTcaLAR1* and *pTcaLAR2* in *Theobroma cacao*. (C) *pMolLAR1* and *pMolLAR2* in *Moringa oleifera.* A comprehensive list of CREs for each promoter is provided in (Additional file 4)

### Loss of LAR in *Brassicaceae*

To confirm the absence of LAR in the *Brassicaceae* family we performed a synteny analysis that first compared the genome sequences of three species belonging to different *Brassicales* families: *Moringa oleifera* (*Moringaceae*), *Carica papaya* (*Caricaceae*), and *Bretschneidera sinensis* (*Bretschneideraceae*) in context with the outgroup species *Theobroma cacao* (*Malvaceae*) and *Vitis vinifera* (*Vitaceae*) (Figure 6A). Microsynteny confirmed the presence of LAR among above-mentioned species of the *Brassicales* order and the outgroups. However, when repeating the analysis only among members of the *Brassicales* order, this time including the species from the *Brassicaceae* family: *Capsella rubella, Boechera stricta, Arabidopsis thaliana, Arabidopsis halleri, Eutrema salsugineum*, and *Brassica rapa* no *LAR1* gene was detected among the *Brassicaceae* (Figure 6B).

After discovering two independent LAR clades through the phylogenetic analysis, it was necessary to determine whether *LAR2* is present among the *Brassicaceae* species. This synteny analysis included as reference the genome sequence of *V. vinifera* with the region where LAR2 is localized. Microsynteny was only observed between *V. vinifera*, *T. cacao*, and *M. oleifera*, corresponding to the LAR2 presence in these species as revealed by the phylogenetic analysis. Like *LAR1*, *LAR2* was also not detected in any *Brassicaceae* species in this synteny analysis (Figure 6C).

**Figure 6.**
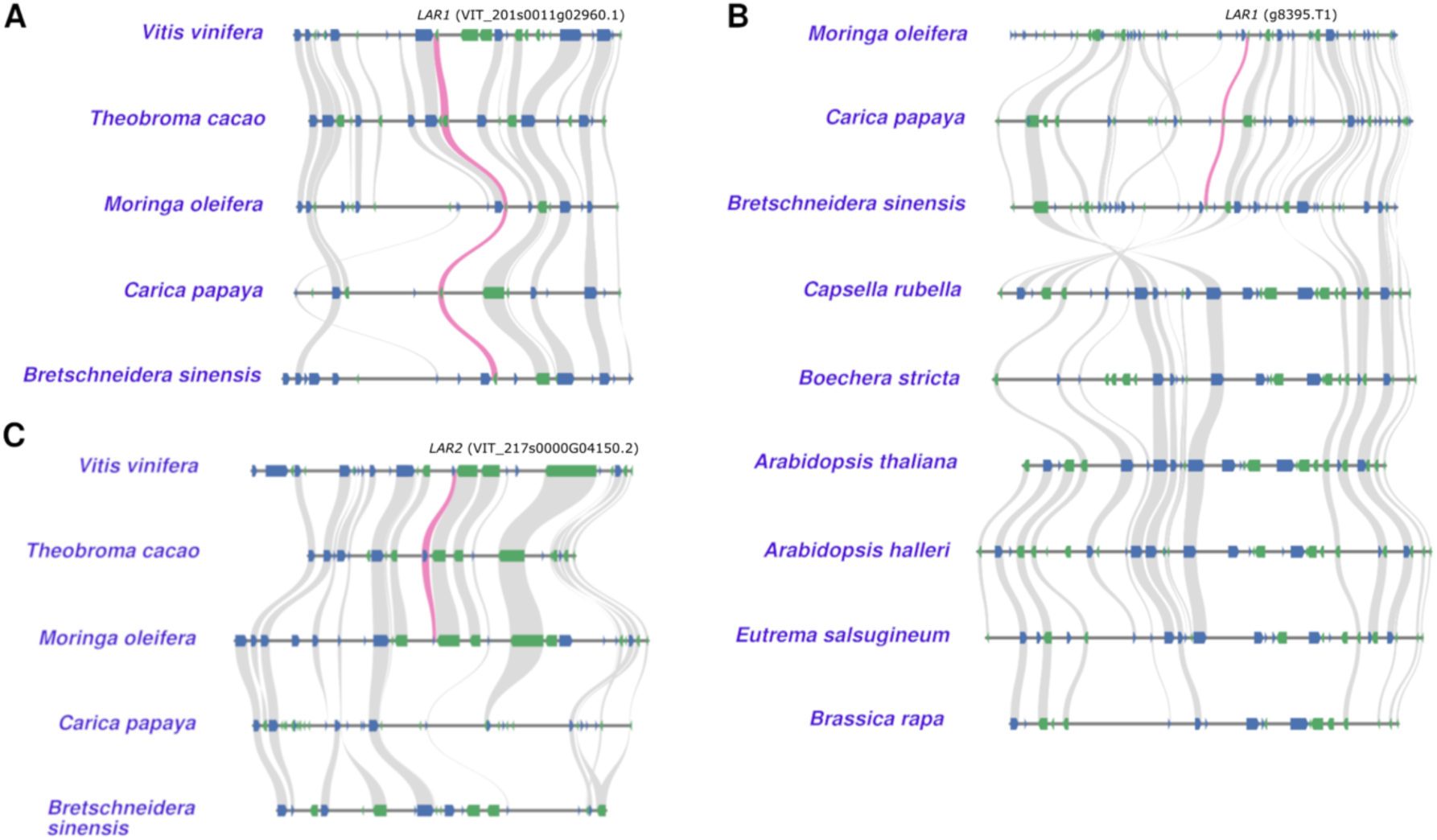
Synteny gene identification of species in the *Brassicales* order. (A) Synteny of *LAR1* (indicated in pink) among the genome sequences of *V. vinifera, T. cacao, M. oleifera, C. papaya,* and *B. sinensis.* (B) Synteny of *LAR1* among the genome sequences of *M. oleifera, C. papaya, B. sinensis, C. rubella, B. stricta, A. thaliana, A. halleri, E. salsugineum,* and *B. rapa.* (C) Synteny of LAR2 among the genome sequences of *V. vinifera, T. cacao, M. oleifera, C. papaya,* and *B. sinensis.* A complete list of syntenic gene positions and coordinates used for this analysis is provided in Additional file 5.

## DISCUSSION

The flavonoid biosynthesis pathway is generally believed to be well conserved across different plant taxa [18, 40]. The evolution of flavonoids has been extensively studied, including the distinct evolutionary mechanisms that led to some of the individual branch products of the pathway [41–43], evolution of this biosynthetic pathway and novel traits are often achieved by changes in transcription factors [44–46]. Such findings have provided significant insights into the genetic and enzymatic variations that contribute to the diversity of flavonoid compounds among plant species [47–49]. Research has revealed that gene duplications and subsequent functional divergences are key processes in the evolution of flavonoid biosynthesis [44, 50]. These genetic variations result in a wide array of flavonoid structures, many of which have well reported biological functions, such as antioxidant activity [51], UV protection [52], and attraction of pollinators and seed dispersers [53]. Additionally, studies have highlighted the role of regulatory genes in controlling the expression of genes encoding enzymes involved in flavonoid production [3, 46, 54]. Our findings on LAR highlight the mechanism on which gene duplication, divergence, and loss of the LAR gene have driven functional diversification in the context of proanthocyanidin biosynthesis across plant species.

### Evolutionary Dynamics of *LAR* Gene Duplications

The phylogenetic analysis revealed that LAR has undergone complex evolutionary trajectories in both gymnosperms and dicotyledons. Within the dicotyledon group, *LAR* genes are segregated into two major clades, referred as LAR1 and LAR2, which likely originated from an ancient duplication event in the common ancestor of the eudicotyledons. This bifurcation of *LAR* genes into distinct evolutionary paths denotes selective pressure favoring the maintenance of both paralogs, possibly due to their acquisition of distinct or complementary functions.

Consistent presence of both LAR1 and LAR2 in multiple dicot families, such as *Malvaceae*, *Rosaceae*, and *Vitaceae*, suggests that these paralogs have been conserved possibly due to functional divergence. Evidence of this phylogeny pattern in LAR has been previously reported by Wang et al. [27], who reported that plant LARs present three distinctive groups; including monocotyledons, gymnosperms and dicotyledons with the latter further splitting into two clades. The authors also noted the presence of sequence-level divergence due to a DNA transversion and transition in a codon of the LAR-specific motif *ICCN* [27]. This finding was corroborated by our results, which also contain a serine residue in *LAR1* substituted by alanine in *LAR2*.

However, we also observed a similar duplication pattern in gymnosperms, with gene copies clustering into two distinct clades, LARI and LARII. Similar to dicots, these clades do not correspond to recent tandem duplication, as the copies are found in separate genomic regions, suggesting an ancient duplication event followed by long-term divergence. Notably, LARII in gymnosperms shows strong amino acid sequence similarity to the LAR paralogs in dicots, supporting the hypothesis that the duplication originated in a common ancestor of seed plants [55]. Under this scenario, one of the ancestral copies was potentially retained and further duplicated within dicots, giving rise to LAR1 and LAR2, while no such duplication has been observed in monocots. This evolutionary model implies that the LAR gene family underwent differential retention and lineage-specific diversification across major plant groups. Additionally, *LARII* in gymnosperms tend to show higher expression across several tissues compared to *LARI,* inferring the possibility that regulatory divergence may also have contributed to their functional differentiation and evolutionary paths.

### Functional Divergence Between LAR1 and LAR2

The functional differentiation of LAR1 and LAR2 is supported not only by their phylogenetic divergence but also by their distinct expression profiles. Gene expression analyses across several dicot species revealed consistent differences between the two copies. For instance, *TcaLAR1* is predominantly expressed in vegetative tissues such as leaves and seedlings, highlighting its role in primary metabolism and general stress responses. This is further supported by coexpression of *TcaLAR1* with stress-related genes, including *proline-rich protein-1*, *bax inhibitor-1* and *Calmodulin7*, which are known to be involved in abiotic stress tolerance [56–58] . In contrast, *TcaLAR2* presented higher expression in reproductive tissues, such as pods and pistils, indicating a more specialized function in these organs.

*TcaLAR2* was found to be coexpressed with genes involved in structural and regulatory processes, including the protein SPIRAL1, known for its expression in tissues undergoing rapid cell expansion [59], these findings support its potential role in reproductive tissue development. This pattern of tissue-specific functional differentiation points to a potential adaptive significance of maintaining multiple LAR copies within a given species. This divergence is also evident in gymnosperms. For example, in *T. chinensis* and *G. biloba*, *LARII* consistently showed higher expression across several tissues compared to *LARI*. In *P. pinaster*, *LARII* was strongly expressed in needles and shoots, reinforcing the notion that regulatory divergence is an independent outcome following LAR duplication in gymnosperms and dicots. Variation in expression patterns among species suggests that these LAR duplicates may have undergone lineage-specific adaptations shaped by ecological or developmental constraints. These findings align with previous studies that have demonstrated how gene duplication events can give rise to paralogs through subfunctionalization or neofunctionalization allowing species to optimize their resources and improve their environmental adaptation [60, 61].

Sequence comparisons among LAR paralogs across gymnosperms and dicots further revealed both conserved patterns and lineage-specific divergences, particularly at functionally relevant motifs. Among gymnosperm copies, LARII exhibited sequence similarity to dicot LARs (LAR1 and LAR2), especially at conserved residues such as positions 157 and 168. This pattern, along with the consistently higher expression levels of LARII compared to LARI, supports the hypothesis that LARII may have retained more ancestral structural features, making it a more functionally conserved paralog than LARI. Additionally, the sequence variations identified between LAR1 and LAR2 reinforce the evidence for their functional divergence. One of the most prominent mutations involves S157A within the conserved *ICCN* motif, a region implicated in substrate binding and catalytic specificity of LAR enzymes [27, 62]. Polar amino acid residues like serine are known to facilitate hydrogen bonding with substrates or cofactors, supporting catalytic activity and proper substrate binding [63, 64]. In contrast, nonpolar residues such as alanine contribute primarily to structural stability and maintaining the integrity of the protein through hydrophobic interactions. The substitutions S157A and S167A in LAR2 may therefore alter the enzyme’s interaction dynamics or catalytic behavior. Although changes in protein structure do not directly control gene expression, structural divergence can result in functional specialization, which may in turn be accompanied by differential expression in different tissues. In this context, the specialized expression of *TcaLAR2* in reproductive tissues could reflect a shift in its biochemical role, favored by changes in substrate preferences, activity, or protein-protein interactions. Supporting this, [23] reported a similar expression profile of a *T. cacao LAR* gene located on chromosome 3, which matches the position of *TcaLAR2* in this study. Liu et al. reported a higher expression in pods and seeds, supporting the idea that *TcaLAR2* may be functionally specialized for reproductive tissue processes such as proanthocyanidin biosynthesis during pod development [23]. Another substitution potentially linked to tissue-specific expression is E168D in LAR2, a variation that also lies within a conserved motif. Although both glutamate and aspartate are negatively charged, the shorter side chain of aspartate can reduce enzyme efficacy and alter substrate specificity. While these effects have not been specifically demonstrated in LAR proteins, similar glutamate-aspartate substitutions in other enzymes have been shown to impair catalytic activity and binding properties [65–67]. In addition, the substitution N255M, which lies adjacent to the active site, was also identified. Asparagine (N), conserved in LARI, LARII, and LAR1 is replaced by methionine (M) in LAR2. As a polar residue, N255 commonly participates in hydrogen bonding; these bonds help stabilize the configuration of the active site [68]. Additionally, the presence of a polar sidechain can contribute to the potential electrostatic interactions, enhancing substrate binding [69].

Contrarily, M255 in LAR2 is larger and has a nonpolar sidechain, which may disrupt these interactions with a possible decrease in catalytic efficiency through altered substrate positioning or transition state stabilization [70, 71]. The proximity of these changes to conserved motifs and active sites suggests that they may impact enzyme stability as well. These findings align with a previous report [27], which highlighted the importance of specific amino acid changes in the evolution of LAR in *C. sinensis*.

Together, these amino acid substitutions in LAR sequences point to biochemical divergence that may contribute to tissue-specific functions. Their location within or in proximity to conserved motifs can be interpreted as indicative of evolutionary shifts likely affecting substrate affinity or motif function. While our findings provide evidence for structural differences, they also propose new directions for understanding how such molecular changes might relate to the distinct functional roles observed in LAR across species.

### Role of CREs and TEs in divergence of LAR1 and LAR2 regulation

The diversification of the flavonoid biosynthesis pathway in angiosperms is closely tied to the evolutionary plasticity of its regulatory hotspots [46, 47, 54, 72], including those involved in the reduction of leucoanthocyanidins to catechins via LAR [73, 74]. In this study, the promoter regions of *LAR1* and *LAR2* from *V. vinifera, T. cacao,* and *M. oleifera* were analyzed to assess regulatory divergence through the identification of cis-regulatory motifs (CREs) and transposable elements (TEs). Our findings propose that promoter divergence, modulated by cis-element variation and TE insertions, may underline functional specialization of the two LAR paralogs in a lineage specific manner. Promoter region analysis revealed a common pattern involving MYB, and bHLH binding elements across all species, consistent with the well documented role of members of these TFs families in regulating flavonoid biosynthesis [75]. Several MYB TFs are established regulators of the phenylpropanoid and flavonoid pathways [76, 77], specific members such as MYBPA1 and MYBPA2 in *V. vinifera,* and MYB5 and MYB14 in *M. truncatula* have been shown to activate the expression of *LAR* and *ANR* genes, thereby controlling proanthocyanidin biosynthesis [78–80]. Similarly, bHLH transcription factors, which function as part of the MBW complex, not only assist MYBs in regulating anthocyanin and proanthocyanidin production but also modulate gene expression dynamically in response to developmental stages and environmental stimuli [81]. For example, in *Vitis amurensis,* the transcription factor *VabHLH137* is induced upon fungal infection and promotes PA biosynthesis by directly activating *VaLAR2,* implying a role in linking stress-responsive signaling to the activation of flavonoid biosynthetic genes [82]. While the presence of MYB and bHLH binding sites in LAR promoters does not provide functional evidence, it suggests the potential regulation by these TF families. This supports the hypothesis that LAR expression, and by extension, PA biosynthesis may be modulated in response to developmental and environmental cues, consistent with the dual roles of these specialized metabolites in pigmentation and protection [83].

While sharing general transcriptional activation elements, such as TATA and CAAT boxes, LAR1 and LAR2 present clear divergence in their promoter architecture across the three analyzed species, indicating a possible subfunctionalization or neofunctionalization post-gene duplication [84]. In *V. vinifera, VviLAR1* promoter uniquely displays an enrichment of abscisic acid (ABA)-responsive elements (ABREs), such as ABRELATERD1, ABRERATCAL, and ACGTATERD1. ABREs are recognized by bZIP transcription factors and mediate transcriptional responses to ABA, especially under drought, salinity, and cold stress [85, 86]. The presence of these motifs aligns with previous findings showing that ABA induces the expression of *LAR* and *ANR* genes during fruit maturation [87, 88]. The abundance of these elements in *VviLAR1* suggests that hormonal regulation via abscisic acid could modulate PA biosynthesis more prominently in this paralog, especially under drought or ripening stages. In contrast, the *VviLAR2* promoter presented a higher abundance of MYC-related cis-elements (E-boxes), which are canonical binding sites for bHLH transcription factors such as MYC1, MYC2 and MYCA1 [89–91]. In *V. vinifera*, MYC1 has been shown to interact with R2R3-MYB proteins to coregulate the biosynthesis of anthocyanins and PAs during berry development [90]. bHLH-binding motifs in the *VviLAR2* promoter may therefore reflect a more prominent integration of bHLH signaling in its transcriptional regulation.

This is consistent with the tissue-specific expression patterns of this paralog, particularly regarding its higher expression levels shown in grape berries. Moreover, evidence from *A. thaliana* shows that bHLH transcription factors cooperatively regulate flavonoid biosynthesis genes by binding to conserved E-box (CANNTG) motifs within light-responsive promoter regions [92]. These motifs often function as part of light regulatory units (LRUs), which also include MYB recognition elements (MRE) and ACGT-containing elements (ACE), and are necessary for the coordinated activation of genes encoding enzymes in the flavonol biosynthesis such as CHS, F3H, and FLS under light stimuli. bHLHs also contribute to the regulation of PA biosynthesis, including the transcriptional control of LAR genes. Previous studies of the promoter regions of both LAR genes in *V. vinifera* have shown that *VviLAR1* expression is sensitive to light conditions, with higher expression under light and repression in the dark, correlating with the activity of bHLH transcription factors such as MYC2 and MYCA1, the latter being upregulated in callus tissues under dark conditions [93]. Our promoter analysis supports this regulatory patterns, *pVviLAR1* presents multiple light-responsive elements (LREs), such as GT1-related motifs, consistent with the light-induced expression profile reported by Chen et al. [93] (Additional file 4). Overall, these findings suggest that *VviLAR1* is predominantly regulated by light-responsive pathways, whereas the enrichment of MYC-binding motifs in *pVviLAR2* implies that this paralog may be more responsive to bHLH-mediated regulation.

In *T. cacao,* both promoters contain core MYB-binding elements, supporting the involvement of multiple MYB transcription factors in regulating LAR gene expression. However, *pTcaLAR1* exhibited a higher overall abundance of these motifs, indicating a potentially more complex regulation. Among these elements is MYBCORE (CNGTTR), a canonical target for R2R3-MYB proteins. This motif is recognized by MYB.Ph3 in *Petunia hybrida*, known to be involved in regulating anthocyanins and PA biosynthetic genes in petal epidermal cells [94]. Similarly, *DkMYB4* in *Diospyros kaki* has been shown to directly bind the MYBCORE motif in the promoter of *DkANR,* downregulation of *DkMYB4* leads to reduced expression of *DkANR* and thus lower PA accumulation [95]. Another MREs found in both promoters is MYBPZM, recognized by the gene *MYBP* identified in *Zea mays* and involved in red kernel pigmentation via activation of A1 gene expression [96]. Interestingly, *pTcaLAR2* contains the MYBGAHV motif (TAACAAA) which is not found in *pTcaLAR1*. This element is the specific binding site for GAMyB, a gibberellin (GA) inducible MYB transcription factor, shown to activate GA-responsive genes such as high-pl α-amylase in barley aleurone cells [97]. The presence of this cis-element in *pTcaLAR2* suggests that this paralog may be selectively responsive to GA signaling, potentially linking its regulation to flower and seed development. GA is known to play critical roles in the transition from vegetative to reproductive phase including the formation of floral organs such as seed and pollen tube growth [98, 99]. This aligns with the enrichment of POLLEN1LELAT52 motif in *pTcaLAR2,* a cis-element known for pollen-specific expression [100]. Additionally, a higher abundance of LREs, including GT-motifs and GATA boxes, was identified in *pTcaLAR2*, hinting at light-responsive expression in this paralog. The presence of these LREs in *LAR2* promoter is consistent with previous transcriptomic studies positively correlating light exposure to proanthocyanidin production [101–103], and aligns with the higher expression levels observed in pods where light-mediated regulation may be more relevant, specifically during pod development and ripening. These findings further support the hypothesis that *TcaLAR2* may be subject to a more restricted regulatory profile, with activity concentrated mostly around reproductive organs. This pattern is consistent with its expression levels and its potential specialization in PA biosynthesis during specific developmental stages.

In *M. oleifera*, the *MolLAR1* promoter shows a particular enrichment of MYB and MYC-binding sites such as MYBCORE and MYCCONSENSUSAT compared to *pMolLAR2*, highlighting that LAR1 may retain a more canonical role in proanthocyanidin biosynthesis. In contrast, *pMolLAR2* is characterized by a higher number of flavonoid-related motifs, including MYBPZM and MYBPLANT with the latter being reported as binding site for transcription factors such as *AmMYB305* and *AmMYB340* in *Antirrhinum majus*, which regulate phenylpropanoid and lignin biosynthesis [104]. This suggests that *MolLAR2* may have acquired a more specialized regulatory function in flavonoid biosynthesis. Additionally, both *MolLAR1* and *MolLAR2* promoter regions in *M. oleifera* contain multiple WRKY-binding sites, including WRKY71OS and WBOXNTERF3, supporting the idea that WRKY-mediated stress signaling might play a role in modulating proanthocyanidin biosynthesis in this species [105].

The differential presence of TE insertions within the LAR promoter regions may also play a significant role in the regulatory divergence between *LAR1* and *LAR2*. In *V. vinifera*, both promoters contain TE insertions, but differ in their composition and position. These elements are inserted in different regions within the promoters and genes likely influencing their expression. TE insertion can influence gene regulation by introducing or removing cis-regulatory elements, modulating chromatin accessibility, or even acting as alternative enhancers or repressors [106]. For instance, LTR retrotransposons have been associated with pigmentation loss in crops such as grape, blood orange and apple by modulating the expression of anthocyanin biosynthesis genes [107–109]. However, the extent to which these regulatory disruptions in the anthocyanin pathway influence proanthocyanidin production remains unclear.

Additionally, the presence of a TIR/Mutator-type element exclusively in the *VviLAR2* promoter implies that this paralog might have originated from a transposon-mediated duplication event, likely contributing to its divergent expression profiles in *V. vinifera* [84, 110, 111]. By contrast, no transposable elements were detected in the *LAR* promoters of *T. cacao* and *M. oleifera*, indicating a more conserved promoter architecture.

Across all three species, promoter region analyses revealed the presence of MYB and bHLH binding motifs associated with LAR regulation, suggesting that particular members of these TF families contribute to the conserved control of PA biosynthesis. However, the distribution and abundance of these conserved motifs, along with their proximity to the TSS and presence of species-specific or paralog-specific elements, supports a model in which LAR gene duplicates have acquired distinct regulatory profiles. The functional implications of these changes may include tissue-specific expression, responsiveness to environmental stimuli, or developmental regulation, all of which could dynamically modulate proanthocyanidin biosynthesis. This regulatory plasticity may reflect the evolutionary necessity of plants to optimize flavonoid production in response to diverse ecological pressures, such as pathogen attack, UV exposure, and nutrient availability [83, 112, 113]. The interplay between conserved regulatory motifs and lineage-specific TE insertions supports broader conclusions regarding the role of gene duplication and promoter evolution in the diversification of plant specialized metabolism [44, 114, 115].

### Insights into the Loss of LAR in *Brassicaceae*

The absence of LAR genes in the *Brassicacea* family, as revealed by the synteny analysis, presents an interesting case of metabolic adaptation. While previous studies have not reported the presence of LAR homologs in *A. thaliana* [20] or in species of the *Brassica* genus [116], our results extend this observation to a broader sampling within the family. Given that *A. thaliana* predominantly accumulates PAs composed of (-)-epicatechin, the presence of (+)-catechins, and consequently a functional LAR gene, has not yet been reported in this species [20]. The retention of other proanthocyanidin biosynthesis genes in *Brassicaceae* [116] points to a functional compensation mechanism or a shift in metabolic compounds that led to LAR not being necessary in the production of proanthocyanidins. It is possible that the pathway in *Brassicaceae* species has evolved to rely solely on ANR activity for the synthesis of PAs, unlike *M. truncatula* which dual activity of LAR has been reported in the production of catechins and its involvement with ANR in the synthesis of epicatechins [21]. This selective loss within the *Brassicaceae* family hints to the likelihood that alternative enzymatic pathways or redundant mechanisms might compensate for the absence of LAR. Similar patterns of LAR loss have been observed in other lineages. For example, both *Zea mays* and *Sorghum bicolor* lack apparent LAR homologs [117]. While maize shows low PA levels and relies solely on ANR to produce epicatechin-based PAs, sorghum is a PA-rich species with a more complete set of regulatory transcription factors, including TT2-type MYBs, synthesizing catechin and procyanidin B3 dimers as major PA components queries [117]. This divergence illustrates that although both species share the absence of LAR, their metabolic compensation strategies and PA profiles are lineage-specific. Such metabolic shifts are not uncommon in plant specialized metabolism. Similar cases include the mutual exclusivity of anthocyanin and betalain pathways in the *Caryophyllales* order [118, 119], or the presence of glucosinolates that are largely confined to *Brassicales* [120]. These examples highlight how evolutionary pressures shape biosynthetic pathways differently across taxa.

Although *A. thaliana* has been instrumental in elucidating the core steps of the flavonoid biosynthesis; its limitations, such as the lack of LAR, highlights the importance of exploring additional model species. Recent reviews have proposed new model systems better suited for studying the full diversity of PA biosynthetic pathway, particularly in species where both LAR and ANR are present [31, 121]. In this context, crops such as *V. vinifera* and *T. cacao,* where the LAR function is well documented and linked to seed development and PA accumulation [22, 23], could provide opportunities to better understand species-specific enzymatic adaptations.

This study confirms and adds to previous findings by providing additional evidence that LAR is likely absent from a broader range of *Brassicaceae* species. In addition, it highlights the dynamic nature of plant specialized metabolism emphasizing the value of integrating model and non-model systems to fully grasp the evolution and diversification of biosynthetic pathways.

## CONCLUSION

We conclude that the four distinct *LAR* clades have functional differences i.e. with respect to gene expression and enzymatic properties. However, a universal pattern of functional difference was not discovered and might not exist due to secondary evolutionary events. Interestingly, LAR1 was represented by a higher number of sequences compared to LAR2. While this may partly reflect differences in species-richness, genome sequence availability, and annotation quality, it could also indicate that LAR1 is more broadly retained and functionally important across dicots. This overrepresentation of LAR1 may reflect a central role in proanthocyanidin biosynthesis and stress responses as indicated by coexpression results. Future research could focus on exploring the biochemical properties of LAR1 and LAR2 enzymes in greater detail to better understand whether these differences result from subfunctionalization or neofunctionalization. Additionally, investigating the interaction dynamics influencing substrate specificity, could provide valuable insights into the evolutionary and ecological significance of proanthocyanidin metabolism in plants. Most importantly, future studies are encouraged to differentiate between members of the LAR1 and LAR2 clade in dicots or LARI and LARII clade in gymnosperms, respectively, when reporting about LAR.

## METHODS

### 1. Data collection

Two primary types of sequence datasets were collected for this study: genome assemblies and corresponding annotations for synteny analysis, and *LAR*, *ANR*, and *DFR* coding and peptide sequences for phylogenetic construction.

#### Genomic data for synteny analysis

The genome assemblies and corresponding gene annotation of 11 species were retrieved from Phytozome and NCBI genome database (Table 1). These data were used for synteny analysis to evaluate genomic neighborhood conservation of the *LAR* gene and support putative loss events (See methods section 9).

#### Candidate sequences for phylogenetic analysis

To reconstruct the evolutionary history of *LAR*, *DFR*, and *ANR* genes, candidate orthologs were identified across a broad range of plant species using KIPEs v3 with default parameters [122]. Protein databases were screened using the bait and reference data FlavonoidBioSynBaits_v3.3 provided by KIPEs3, and only hits showing a 100% match in the conserved amino acid residues were retained to infer presence in a plant lineage. Partial candidates were inspected when inferring absence of LAR. The resulting sequences were then cleaned and mapped to their corresponding coding sequences (CDS) using a custom pipeline (see methods section 3). The species tree used to guide synteny comparisons and phylogenetic interpretation was obtained from [123].

### 2. Gene annotation

Since no annotation of the coding sequences was publicly available for *Moringa oleifera*, BRAKER v3.0.8 [124] with default settings was applied to generate a structural annotation of protein-encoding genes. To support the annotation process with hints, several RNA-seq data sets were retrieved from the Sequence Read Archive [125, 126] (Additional file 4) and aligned to the genome sequence with HISAT v2.2.1 [127] using default parameters. Samtools v1.20 [128] was used to sort the resulting BAM file which was then provided to BRAKER3 [124].

### 3. Alignment and phylogenetic tree construction

A collection of LAR, DFR, and ANR sequences were identified using KIPEs v3 [129]. Sequences were only considered if a strict match of 100% was found in the set of conserved amino acid residues defined in the KIPEs3 bait file, which ensures a high-confidence LAR identification. These sequences were selected for further analysis. To improve tree readability, sequences labeled as “putative”, “predicted”, or repetitive sequences were removed using a custom python script (clean_fasta.py), which is publicly available on GitHub: https://github.com/mariamarinr/LAR_Evolution/. This filtering step was designed to preserve taxonomic representation, retaining at least one high-confidence LAR sequence per major plant lineage. After the sequence cleaning, the corresponding coding sequence (CDS) was retrieved for each polypeptide sequence to enable codon-aware alignments. This was done using a custom pipeline that cross-referenced gene identifiers from the amino acid FASTA headers with available genome annotations and CDS files (extract_kipes_cds.py) on GitHub: https://github.com/mariamarinr/LAR_Evolution/. Only sequences with matching identifiers to their corresponding amino acid sequences were retained. In cases where formatting differences in the headers occurred between CDS and protein files, identifiers were standardized to ensure a perfect match. This ensured consistency between protein-level and nucleotide-level data for downstream phylogenetic analysis (see Additional file 6 for a complete list). Codon-aware multiple sequence alignments were constructed using MACSE v2.07 [36, 130] with default parameters. An initial phylogenetic tree was constructed with FastTree v2.1.10 [131] using the WAG substitution model and without bootstrap support for rapid visualization. To confirm the phylogenetic structure and arranged relationships, a more robust analysis was performed using IQ-TREE v2.0.7 [37, 38], with ultrafast bootstrap and 1000 replicates. Tree visualization and annotation were conducted using the Interactive Tree OF Life (iTOL) v.7 [132].

### 4. Gene expression analysis

Paired-end RNA-seq datasets were retrieved from the Sequence Read Archive using fastq-dump v2.8.1 [125]. Gene expression quantification, including raw counts and Transcripts Per Million (TPMs), were obtained using kallisto v0.44 [133] with default parameters. Customs R scripts (prep_expression_matrix.R and LMM_expression_analysis.R) available through GitHub (https://github.com/mariamarinr/LAR_Evolution/) were developed to prepare the expression data matrix and generate violin plots using the ggplot2 package v3.5.1 [134] illustrating the variation in gene expression (TPMs) across multiple samples (Additional file 7). These plots were specifically used to visualize the expression levels of *LAR* across specific plant tissues in different species, aiming to identify patterns in gene expression and to gain insights into potential functions. To assess the statistical significance of the difference in gene expression levels between the two *LAR* copies, a linear mixed-effects model (LMM) was employed. The model was formulated as:

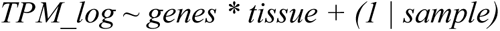

where *TPM_log* represents the log-transformed transcript abundance, *genes* and *tissue* are fixed effects, and *sample* is included as a random effect to account for variability across individual samples. The inclusion of the random effect was essential to control for potential sample-specific variability, thereby ensuring more accurate estimates of fixed effects and reducing the risk of false positives. The model was fitted using restricted maximum likelihood (REML), and statistical significance for the fixed effects was assessed using t-tests with Satterthwaite’s method, as implemented in the lmerTest package v3.1-3 [135]. To further explore differences in gene expression across plant tissues and provide a more detailed evaluation of tissue-specific patterns for each LAR copy, a post-hoc pairwise comparison was conducted.

This analysis was performed using the emmeans function v1.10.1 [136], which computed estimated marginal means (EMMs) [137] and pairwise contrasts for the interaction between genes and tissue. Specifically, pairwise comparisons were made for the genes factor within each level of the tissue factor, using t-tests to assess the significance of differences between the two LAR copies across the various tissues. To account for multiple comparisons, p-values were adjusted using the Bonferroni method. All statistics analysis were performed using R v4.3.3 [138].

### 5. Functionalization analysis

In order to explore the functional divergence of the *LAR* gene copies in the same species, a screen for evidence of subfunctionalization and neofunctionalization was conducted. This involved comparing the sequence profiles of the two *LAR* copies to identify potential nonsynonymous substitutions that would lead to novel functions that had evolved since the duplication event. Specifically, we utilized sequence alignments using MAFFT v7.490 [139] and visualized in JalView v.2.11.4.1 [140] and ChimeraX v.1.8 [141] to detect differences in the amino acid positions within the LAR proteins, which could indicate changes in functional domains or active sites. Additionally, the alignments were subjected to an analysis with CoSeD (https://github.com/k-georgi/CoSeD) to identify specific amino acid positions that show systematic differences between the two groups of LAR sequences.

### 6. Coexpression analysis

Coexpression analysis was performed to identify genes that present similarity in expression with the *LAR* genes of interest. A list with the *LAR* genes and a count table with normalized TPM values were used for this analysis. Pairwise Spearman correlation coefficients for gene expression values across all samples were calculated for each *LAR* gene in the list. The coexpressed genes were assessed based on their Spearman correlation values, adjusted p-values, and total expression level. Genes with low expression levels were excluded, and only gene pairs with a correlation coefficient >0.65 and an adjusted p-value < 0.05 were considered significant. This coexpression analysis was implemented in a Python script - coexp3.py [50]. The functional annotation of the genes belonging to *V. vinifera* (Vvinifera_457_v2.1), *T. cacao* (Tcacao_523_v2.1), and *M. oleifera* (GCA_021397835.1) were derived from *M. truncatula* and *A. thaliana* using the Python script - construct_anno.py [50] available at: https://github.com/bpucker/ApiaceaeFNS1.

### 7. Identification of cis-regulatory elements (CREs) in LAR promoters

To investigate the differences in the regulatory mechanisms of *LAR1* and *LAR2* gene copies, the promoter region of each gene was extracted and analyzed for cis-regulatory elements. For *V. vinifera*, the *LAR1* and *LAR2* promoters previously reported by [93] were used, available in GenBank under accession numbers MT586116 and MT586117, respectively. For *T. cacao* and *M. oleifera*, paired-end RNA-seq reads were retrieved from the Sequence Read Archive (SRA) using fasterq-dump v.3.0.3 [142]. The complete list of SRR accessions is provided in Additional file 4. To ensure accuracy in the identification of the transcription start sites (TSS), an alignment of these reads to their respective genome assemblies was performed using HISAT v2.2.1 [127], a splice-aware aligner optimized for short RNA-seq reads. This mapping workflow was automated using a custom Bash script (hisat2_mapping.sh) available at the GitHub repository: https://github.com/mariamarinr/LAR_Evolution/. This pipeline also handled the conversion, merging, sorting, and indexing of alignment files with Samtools v1.15 [128] for each species. The resulting sorted BAM files (available at: Link to final repository) were visualized using the Integrative Genomics Viewer (IGV) [143] to identify the regions with transcriptional activity upstream of LAR coding sequences. Based on RNA-seq coverage patterns, the transcription start site (TSS) was inferred for each *LAR* gene copy, and promoter regions were defined as approximately 1500 bp upstream of the TSS. These promoter sequences were extracted from the genome assemblies using BEDtools v2.30 [144] (Additional fasta file 4) and analyzed for potential cis-regulatory elements using the PLACE database [39] at https://www.dna.affrc.go.jp/PLACE/.

### 8. Identification of transposable elements (TEs)

To evaluate whether transposable elements (TEs) may contribute to the regulation of *LAR* gene expression, genome-wide annotation of TEs was performed for the species: *V. vinifera*, *T. cacao*, and *M. oleifera*. TE annotation was conducted for each genome sequence using the Extensive *de-novo* TE Annotator (EDTA) v.2.2.2 [145]. EDTA was run in sensitive mode with both annotation and evaluation options enabled to ensure a complete detection of both abundant and low-copy TEs. To identify overlaps between TEs and *LAR* loci, the intersect function of BEDTools v2.30 [144] was employed. This approach was intended to reveal TE insertions located within or adjacent to *LAR* genes or their promoter regions across the three species. Each overlapping TE was classified by type (e.g., LTR retrotransposons, TIR DNA transposons) and family, allowing further exploration of whether particular TE lineages were commonly linked with LAR regulatory regions. The complete list of overlapping TE insertions and their annotations is provided in Additional file 4.

### 9. Synteny analysis

Synteny analysis using JCVI/MCscan [146] was performed to visualize conserved genomic regions across the genome sequences of multiple species (Table 1). Flanking regions surrounding the identified *LAR* loci, typically including 10 to 15 protein coding genes upstream and downstream, were manually selected based on their physical proximity to LAR and the preservation of local gene order, serving as reference points. Gene connections between species were manually validated and revised by comparing the predicted syntenic blocks with the phylogenetic relationships reported by [123], ensuring consistency between gene conservation and species divergence. To identify homologous genes or syntenic regions that might be missing in current annotations, BLAST v.2.13.0 [147] databases were constructed for multiple species using the ‘makeblastdb’ command, specifying nucleotide sequence types. Query sequences were then searched against these databases using ‘tblastn’.

**Table 1.** List of species used for synteny analysis.

| Species | Data set ID | Database | Reference |
| --- | --- | --- | --- |
| *Vitis vinifera* | Vvinifera_457_v2.1 | Phytozome | [148] |
| *Theobroma cacao* | Tcacao_523_v2.1 | Phytozome | [149] |
| *Bretschneidera sinensis* | GCA_018105755.1 | NCBI | [150] |
| *Moringa oleifera* | GCA_021397835.1 | NCBI | [151] |
| *Carica papaya* | Cpapaya_113_ASGPBv0.4 | Phytozome | [152] |
| *Eutrema salsugineum* | Esalsugineum_173_v1.0 | Phytozome | [153] |
| *Capsella rubella* | Crubella_474_v1.1 | Phytozome | [154] |
| *Boechera stricta* | Bstricta_278_v1.2 | Phytozome | [155] |
| *Arabidopsis halleri* | Ahalleri_765_v2.1.0 | Phytozome | [156] |
| *Arabidopsis thaliana* | Athaliana_447_Araport11 | Phytozome | [157, 158] |
| *Brassica rapa* | GCA_003434825.1 | NCBI | [159] |

## Acknowledgements

This work was supported by the BMBF-funded de.NBI Cloud within the German Network for Bioinformatics Infrastructure (de.NBI) (031A532B, 031A533A, 031A533B, 031A534A, 031A535A, 031A537A, 031A537B, 031A537C, 031A537D, 031A538A). We are grateful to de.NBI and the Center for Biotechnology (CeBiTec) at Bielefeld University for providing us with the resources and setting to conduct the necessary computational analyses. We are also thankful to the members of the Plant Biotechnology and Bioinformatics group at University of Bonn for their helpful discussions and valuable comments during the manuscript revision process. Open Access funding enabled and organized by Project DEAL and the University of Bonn.

## Declarations

### Authors’ contributions

MMR and BP conceptualized the research project. MMR carried out the analyses and prepared figures. MMR wrote the manuscript. MMR and BP revised the manuscript. The authors read and approved the final manuscript.

### Availability of data and materials

All the genomic, transcriptomic and annotation data used in this study was obtained from the NCBI Sequence Read Archive (SRA) and Phytozome. The data charts supporting the results and conclusions are included in the article and additional files. All the sequences and scripts used in the different analyses have been deposited in our GitHub repository (https://github.com/mariamarinr/LAR_Evolution/) and additional files.

### Ethics approval and consent to participate

Not applicable.

### Consent for publication

Not applicable.

