## Additional file 2 for "Evolutionary Dynamics of the Proanthocyanidin Biosynthesis Gene *LAR*": Amino acid conservation values.docx

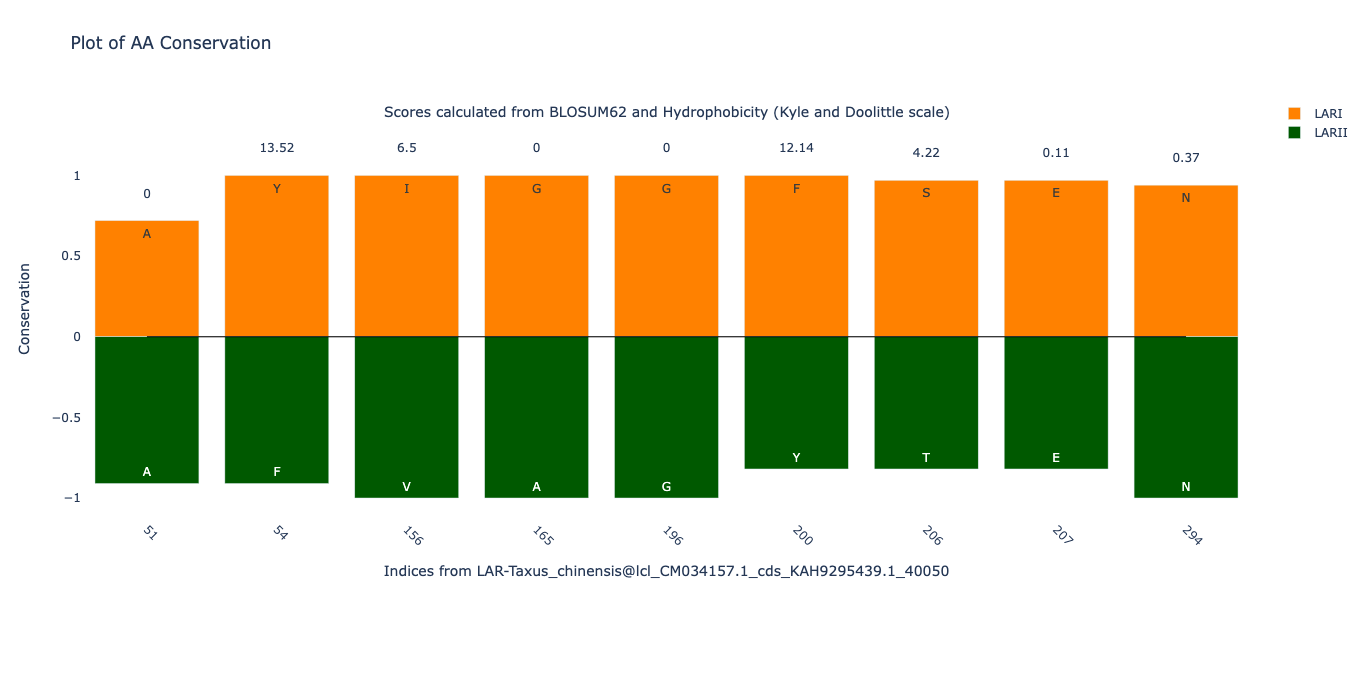

**Figure S9.** Bar plot showing the conservation values of amino acid residues within specific LAR motifs, with positions referenced to *Taxus chinensis*. Amino acid substitutions characteristic of LARI are highlighted in orange, while those of LARII are shown in green. Conservation scores were calculated based on BLOSUM62 substitution matrices and Hydrophobicity indices.

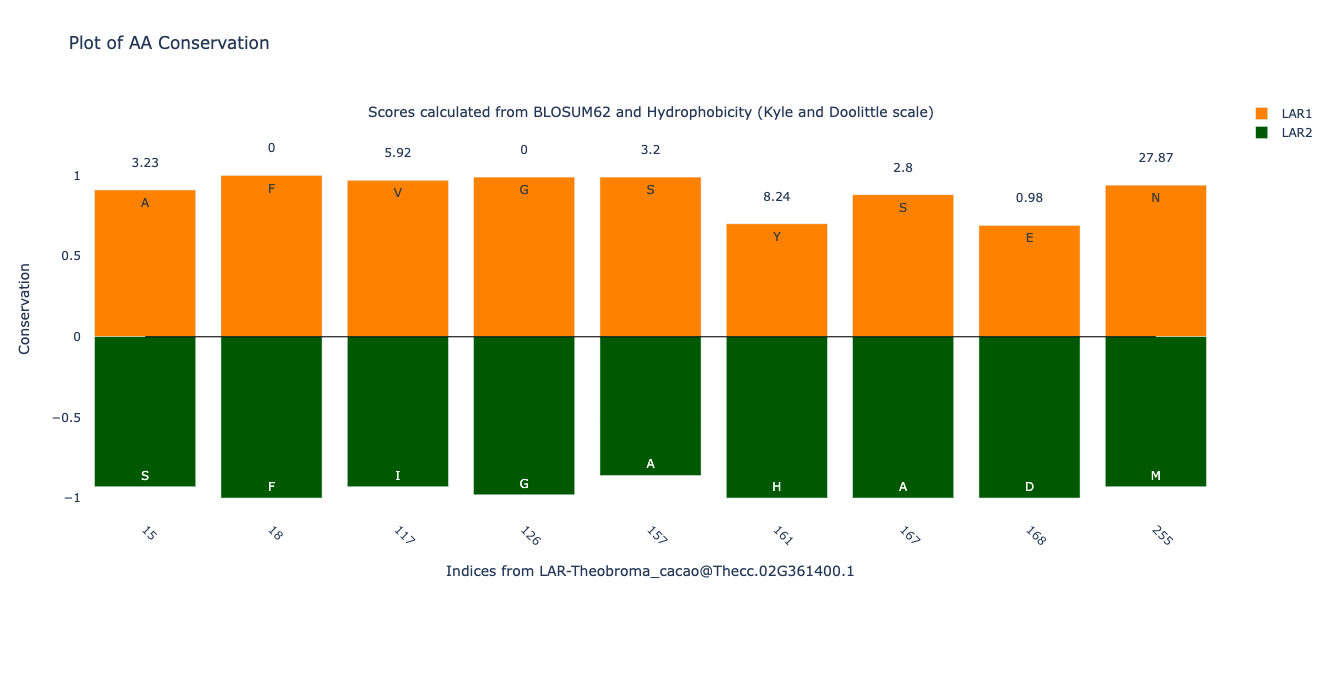

**Figure S10.** Bar plot showing the conservation values of amino acid residues within specific LAR motifs, with positions referenced to *Theobroma cacao*. Amino acid substitutions characteristic of LAR1 are highlighted in orange, while those of LAR2 are shown in green. Conservation scores were calculated based on BLOSUM62 substitution matrices and Hydrophobicity indices.
