## Additional file 1 for "Evolutionary Dynamics of the Proanthocyanidin Biosynthesis Gene *LAR*"

### SUPPLEMENTARY FILE 1

#### Gymnosperm species

#### *Taxus chinensis*

Linear mixed model fit by REML. t-tests use Satterthwaite's method ['lmerModLmerTest']
Formula: TPM_log ~ Genes * Tissue + (1 | Sample)
Data: data_long
REML criterion at convergence: 461.6

Scaled residuals:

| Min | 1Q | Median | 3Q | Max |
|---|---|---|---|---|
| -2.62633 | -0.65638 | -0.01874 | 0.54624 | 2.75311 |

Random effects:

| Groups | Name | Variance | Std.Dev. |
|---|---|---|---|
| Sample | (Intercept) | 0.3826 | 0.6186 |
| Residual | 0.5987 | 0.7737 | |
| Number of obs: | 172 | groups: | Sample, 86 |

Fixed effects:

| | Estimate | Std. Error | df | t value | Pr(>\|t\|) | |
|---|---|---|---|---|---|---|
| (Intercept) | 1.2046 | 0.3133 | 137.1481 | 3.845 | 0.000184 | *** |
| GenesLAR2 | 0.871 | 0.346 | 79 | 2.517 | 0.013851 | * |
| TissueBud and Leave | 1.2824 | 1.039 | 137.1481 | 1.234 | 0.219219 | |
| TissueCambial meristem | -1.1068 | 0.4242 | 137.1481 | -2.61 | 0.010074 | * |
| TissueCell line | -0.4101 | 0.4102 | 137.1481 | -1 | 0.319115 | |
| TissueCone | 0.9612 | 0.5116 | 137.1481 | 1.879 | 0.062373 | . |
| TissueLeaf | 0.8637 | 0.3531 | 137.1481 | 2.446 | 0.015698 | * |
| TissueRoot | 0.4388 | 0.5116 | 137.1481 | 0.858 | 0.392537 | |
| GenesLAR2:TissueBud and leave | 1.5703 | 1.1477 | 79 | 1.368 | 0.175117 | |
| GenesLAR2:TissueCambial meristem | 3.9742 | 0.4685 | 79 | 8.482 | 9.98E-13 | *** |
| GenesLAR2:TissueCell line | 3.3827 | 0.4531 | 79 | 7.466 | 9.48E-11 | *** |
| GenesLAR2:TissueCone | 1.3847 | 0.5651 | 79 | 2.451 | 0.016475 | * |
| GenesLAR2:TissueLeaf | 0.9309 | 0.39 | 79 | 2.387 | 0.019378 | * |
| GenesLAR2:TissueRoot | 0.7534 | 0.5651 | 79 | 1.333 | 0.186281 | |

Signif. codes: 0 ‘***’ 0.001 ‘**’ 0.01 ‘*’ 0.05 ‘.’ 0.1 ‘ ’ 1

**Table 1S.** Estimated marginal means and pairwise comparison for the species *Taxus chinensis*. Degrees-of-freedom method: kenward-roger. Confidence level used: 0.95.

| ***Estimated Marginal Means*** | ***Contrasts*** |
|---|---|
| Tissue = Bark:<br>Genes emmean SE df lower.CL upper.CL<br>LAR1 1.2046 0.313 137 0.585 1.824<br>LAR2 2.0756 0.313 137 1.456 2.695<br><br>Tissue = Bud and leave:<br>Genes emmean SE df lower.CL upper.CL<br>LAR1 2.4869 0.991 137 0.528 4.446<br>LAR2 4.9282 0.991 137 2.969 6.887<br><br>Tissue = Cambial meristem:<br>Genes emmean SE df lower.CL upper.CL<br>LAR1 0.0977 0.286 137 -0.468 0.663<br>LAR2 4.9430 0.286 137 4.378 5.508<br><br>Tissue = Cell line:<br>Genes emmean SE df lower.CL upper.CL<br>LAR1 0.7944 0.265 137 0.271 1.318<br>LAR2 5.0482 0.265 137 4.525 5.572<br><br>Tissue = Cone:<br>Genes emmean SE df lower.CL upper.CL<br>LAR1 2.1658 0.404 137 1.366 2.965<br>LAR2 4.4215 0.404 137 3.622 5.221<br><br>Tissue = Leaf:<br>Genes emmean SE df lower.CL upper.CL<br>LAR1 2.0683 0.163 137 1.746 2.390<br>LAR2 3.8702 0.163 137 3.548 4.192<br>Tissue = Root:<br>Genes emmean SE df lower.CL upper.CL<br>LAR1 1.6433 0.404 137 0.844 2.443<br>LAR2 3.2678 0.404 137 2.468 4.067 | Tissue = Bark:<br>contrast estimate SE df t.ratio p.value<br>LAR1 - LAR2 -0.871 0.346 79 -2.517 0.0139<br><br>Tissue = Bud and leave:<br>contrast estimate SE df t.ratio p.value<br>LAR1 - LAR2 -2.441 1.090 79 -2.231 0.0285<br><br>Tissue = Cambial meristem:<br>contrast estimate SE df t.ratio p.value<br>LAR1 - LAR2 -4.845 0.316 79 -15.339 <.0001<br><br>Tissue = Cell line:<br>contrast estimate SE df t.ratio p.value<br>LAR1 - LAR2 -4.254 0.292 79 -14.545 <.0001<br><br>Tissue = Cone:<br>contrast estimate SE df t.ratio p.value<br>LAR1 - LAR2 -2.256 0.447 79 -5.050 <.0001<br><br>Tissue = Leaf:<br>contrast estimate SE df t.ratio p.value<br>LAR1 - LAR2 -1.802 0.180 79 -10.017 <.0001<br><br>Tissue = Root:<br>contrast estimate SE df t.ratio p.value<br>LAR1 - LAR2 -1.624 0.447 79 -3.636 0.0005 |

### *Ginkgo biloba*

Linear mixed model fit by REML. t-tests use Satterthwaite's method ['lmerModLmerTest']
Formula: TPM_log ~ Genes * tissue + (1 | Sample)
Data: data_long

REML criterion at convergence: 2493.8

Scaled residuals:

| Min | 1Q | Median | 3Q | Max |
|---|---|---|---|---|
| -3.1077 | -0.5304 | -0.0785 | 0.4852 | 4.7825 |

Random effects:

| Groups | Name | Variance | Std.Dev. |
|---|---|---|---|
| Sample | (Intercept) | 0.3123 | 0.5588 |
| Residual | 0.5487 | 0.7407 | |
| Number of obs: | 954 | groups: | Sample, 477 |

Fixed effects:

| | *Estimate* | *Std. Error* | *df* | *T-value* | *Pr(>\|t\|)* | |
|---|---|---|---|---|---|---|
| *(Intercept)* | *2.7967* | *0.3507* | *820.112* | *7.975* | *5.12E-15* | *** |
| *GenesLAR2* | *-0.2512* | *0.3959* | *464* | *-0.634* | *0.526149* | |
| *tissueCalli* | *-0.784* | *0.405* | *820.112* | *-1.936* | *0.053211* | . |
| *tissueCambium* | *-1.9581* | *0.4676* | *820.112* | *-4.188* | *3.13E-05* | *** |
| *tissueFlower* | *-1.0843* | *0.4103* | *820.112* | *-2.643* | *0.008373* | ** |
| *tissueFruit* | *-1.4112* | *0.4676* | *820.112* | *-3.018* | *0.002624* | ** |
| *tissueKernel* | *-2.6392* | *0.4802* | *820.112* | *-5.496* | *5.20E-08* | *** |
| *tissueLeaf* | *-0.182* | *0.3543* | *820.112* | *-0.514* | *0.607551* | |
| *tissueOvule* | *-0.0745* | *0.3986* | *820.112* | *-0.187* | *0.851769* | |
| *tissueRoot* | *-1.576* | *0.5162* | *820.112* | *-3.053* | *0.00234* | ** |
| *tissueSeedling leaves* | *-0.7702* | *0.4676* | *820.112* | *-1.647* | *0.099897* | . |
| *tissueSeedling stem* | *-2.1762* | *0.6403* | *820.112* | *-3.399* | *0.00071* | *** |
| *tissueStem* | *0.2434* | *0.4247* | *820.112* | *0.573* | *0.566711* | |
| *tissueTesta* | *-2.4819* | *0.6403* | *820.112* | *-3.876* | *0.000115* | *** |
| *GenesLAR2:tissueCalli* | *1.6212* | *0.4572* | *464* | *3.546* | *0.000431* | *** |
| *GenesLAR2:tissueCambium* | *0.6589* | *0.5279* | *464* | *1.248* | *0.212642* | |
| *GenesLAR2:tissueFlower* | *1.637* | *0.4632* | *464* | *3.534* | *0.00045* | *** |
| *GenesLAR2:tissueFruit* | *0.9921* | *0.5279* | *464* | *1.879* | *0.060836* | . |
| *GenesLAR2:tissueKernel* | *0.6385* | *0.5422* | *464* | *1.178* | *0.239495* | |
| *GenesLAR2:tissueLeaf* | *-1.2138* | *0.4* | *464* | *-3.035* | *0.002541* | ** |
| *GenesLAR2:tissueOvule* | *0.1129* | *0.45* | *464* | *0.251* | *0.802091* | |
| *GenesLAR2:tissueRoot* | *1.5588* | *0.5828* | *464* | *2.675* | *0.007746* | ** |
| *GenesLAR2:tissueSeedling leaves* | *-0.1148* | *0.5279* | *464* | *-0.217* | *0.82798* | |

| | | | | | | |
|---|---|---|---|---|---|---|
| *GenesLAR2:tissueSeedling stem* | *2.3996* | *0.7229* | *464* | *3.32* | *0.000973* | ***** |
| *GenesLAR2:tissueStem* | *-1.4876* | *0.4795* | *464* | *-3.102* | *0.002036* | **** |
| *GenesLAR2:tissueTesta* | *0.3654* | *0.7229* | *464* | *0.505* | *0.61349* | |

Signif. codes: 0 '***' 0.001 '**' 0.01 '*' 0.05 '.' 0.1 ' ' 1

**Table 2S.** Estimated marginal means and pairwise comparison for the species *Ginkgo biloba*. Degrees-of-freedom method: kenward-roger. Confidence level used: 0.95.

| ***Estimated Marginal Means*** | ***Contrasts*** |
|---|---|
| tissue = Buds:<br>Genes emmean SE df lower.CL upper.CL<br>LAR1 2.797 0.351 820 2.108 3.485<br>LAR2 2.546 0.351 820 1.857 3.234<br><br>tissue = Calli:<br>Genes emmean SE df lower.CL upper.CL<br>LAR1 2.013 0.202 820 1.615 2.410<br>LAR2 3.383 0.202 820 2.985 3.780<br><br>tissue = Cambium:<br>Genes emmean SE df lower.CL upper.CL<br>LAR1 0.839 0.309 820 0.232 1.446<br>LAR2 1.246 0.309 820 0.639 1.853<br><br>tissue = Flower:<br>Genes emmean SE df lower.CL upper.CL<br>LAR1 1.712 0.213 820 1.295 2.130<br>LAR2 3.098 0.213 820 2.680 3.516<br><br>tissue = Fruit:<br>Genes emmean SE df lower.CL upper.CL<br>LAR1 1.386 0.309 820 0.778 1.993<br>LAR2 2.127 0.309 820 1.519 2.734<br>tissue = Kernel:<br>Genes emmean SE df lower.CL upper.CL<br>LAR1 0.158 0.328 820 -0.486 0.802<br>LAR2 0.545 0.328 820 -0.099 1.189<br><br>tissue = Leaf:<br>Genes emmean SE df lower.CL upper.CL<br>LAR1 2.615 0.050 820 2.517 2.713<br>LAR2 1.150 0.050 820 1.052 1.248<br><br>tissue = Ovule:<br>Genes emmean SE df lower.CL upper.CL<br>LAR1 2.722 0.189 820 2.350 3.094<br>LAR2 2.584 0.189 820 2.212 2.956 | tissue = Buds:<br>contrast estimate SE df t.ratio p.value<br>LAR1 - LAR2 0.251 0.3960 464 0.634 0.5261<br><br>tissue = Calli:<br>contrast estimate SE df t.ratio p.value<br>LAR1 - LAR2 -1.370 0.2290 464 -5.993 <.0001<br><br>tissue = Cambium:<br>contrast estimate SE df t.ratio p.value<br>LAR1 - LAR2 -0.408 0.3490 464 -1.168 0.2436<br><br>tissue = Flower:<br>contrast estimate SE df t.ratio p.value<br>LAR1 - LAR2 -1.386 0.2400 464 -5.766 <.0001<br><br>tissue = Fruit:<br>contrast estimate SE df t.ratio p.value<br>LAR1 - LAR2 -0.741 0.3490 464 -2.122 0.0344<br><br>tissue = Kernel:<br>contrast estimate SE df t.ratio p.value<br>LAR1 - LAR2 -0.387 0.3700 464 -1.046 0.2962<br><br>tissue = Leaf:<br>contrast estimate SE df t.ratio p.value<br>LAR1 - LAR2 1.465 0.0565 464 25.938 <.0001<br><br>tissue = Ovule:<br>contrast estimate SE df t.ratio p.value<br>LAR1 - LAR2 0.138 0.2140 464 0.647 0.5180<br><br>tissue = Root:<br>contrast estimate SE df t.ratio p.value<br>LAR1 - LAR2 -1.308 0.4280 464 -3.058 0.0024<br><br>tissue = Seedling leaves:<br>contrast estimate SE df t.ratio p.value |

```
tissue = Root:
 Genes emmean    SE  df lower.CL upper.CL
 LAR1   1.221 0.379 820    0.477    1.964
 LAR2   2.528 0.379 820    1.785    3.272

tissue = Seedling leaves:
 Genes emmean    SE  df lower.CL upper.CL
 LAR1   2.027 0.309 820    1.419    2.634
 LAR2   1.661 0.309 820    1.053    2.268

tissue = Seedling stem:
 Genes emmean    SE  df lower.CL upper.CL
 LAR1   0.621 0.536 820   -0.431    1.672
 LAR2   2.769 0.536 820    1.717    3.821

tissue = Stem:
 Genes emmean    SE  df lower.CL upper.CL
 LAR1   3.040 0.240 820    2.570    3.510
 LAR2   1.301 0.240 820    0.831    1.772
tissue = Testa:
 Genes emmean    SE  df lower.CL upper.CL
 LAR1   0.315 0.536 820   -0.737    1.366
 LAR2   0.429 0.536 820   -0.623    1.481
```

```
 LAR1 - LAR2    0.366 0.3490 464   1.048  0.2952

tissue = Seedling stem:
 contrast    estimate     SE  df t.ratio p.value
 LAR1 - LAR2   -2.148 0.6050 464  -3.552  0.0004

tissue = Stem:
 contrast    estimate     SE  df t.ratio p.value
 LAR1 - LAR2    1.739 0.2700 464   6.429  <.0001

tissue = Testa:
 contrast    estimate     SE  df t.ratio p.value
 LAR1 - LAR2   -0.114 0.6050 464  -0.189  0.8503

Degrees-of-freedom method: kenward-roger
Confidence level used: 0.95
```

#### *Pinus pinaster*

Linear mixed model fit by REML. t-tests use Satterthwaite's method ['lmerModLmerTest']
Formula: TPM_log ~ Genes * tissue + (1 | Sample)
   Data: data_long

REML criterion at convergence: 613.4

Scaled residuals:

| Min | 1Q | Median | 3Q | Max |
|---|---|---|---|---|
| -2.67838 | -0.36856 | -0.00996 | 0.24 | 2.29392 |

Random effects:

| Groups | Name | Variance | Std.Dev. |
|---|---|---|---|
| Sample | (Intercept) | 0.8802 | 0.9382 |
| Residual | 0.3831 | 0.619 | |
| Number of obs: | 224 | groups: | Sample, 112 |

Fixed effects:

| | Estimate | Std. Error | df | T-value | Pr(>\|t\|) | |
|---|---|---|---|---|---|---|
| *(Intercept)* | 0.10854 | 0.32446 | 142.72131 | 0.335 | 0.73847 | |
| *GenesLAR2* | 3.68975 | 0.2527 | 106 | 14.601 | < 2E-016 | *** |
| *TissueRoot* | 2.47978 | 0.36276 | 142.72131 | 6.836 | 2.19E-10 | *** |
| *Tissueshoot* | 1.11747 | 0.40336 | 142.72131 | 2.77 | 0.00634 | ** |
| *TissueSomatic embryo* | -0.03068 | 0.49562 | 142.72131 | -0.062 | 0.95073 | |
| *TissueStem* | -0.03791 | 0.45886 | 142.72131 | -0.083 | 0.93427 | |
| *TissueZygotic embryo* | -0.10689 | 0.49562 | 142.72131 | -0.216 | 0.82956 | |
| *GenesLAR2:TissueRoot* | -2.65545 | 0.28253 | 106 | -9.399 | 1.27E-15 | *** |
| *GenesLAR2:Tissueshoot* | -1.29568 | 0.31415 | 106 | -4.124 | 7.41E-05 | *** |
| *GenesLAR2:TissueSomatic embryo* | -2.42709 | 0.386 | 106 | -6.288 | 7.33E-09 | *** |
| *GenesLAR2:TissueStem* | 0.85597 | 0.35737 | 106 | 2.395 | 0.01837 | * |
| *GenesLAR2:TissueZygotic embryo* | -3.45121 | 0.386 | 106 | -8.941 | 1.35E-14 | *** |

Signif. codes: 0 '***' 0.001 '**' 0.01 '*' 0.05 '.' 0.1 ' ' 1

**Table 3S.** Estimated marginal means and pairwise comparison for the species *Pinus pinaster*. Degrees-of-freedom method: kenward-roger. Confidence level used: 0.95.

| ***Estimated Marginal Means*** | ***Contrasts*** |
|---|---|
| Tissue = Needles:<br>Genes emmean SE df lower.CL upper.CL<br>LAR1 0.10854 0.324 143 -0.533 0.750<br>LAR2 3.79830 0.324 143 3.157 4.440<br><br>Tissue = Root:<br>Genes emmean SE df lower.CL upper.CL<br>LAR1 2.58832 0.162 143 2.268 2.909<br>LAR2 3.62263 0.162 143 3.302 3.943<br><br>Tissue = shoot:<br>Genes emmean SE df lower.CL upper.CL<br>LAR1 1.22601 0.240 143 0.752 1.700<br>LAR2 3.62009 0.240 143 3.146 4.094<br><br>Tissue = Somatic embryo:<br>Genes emmean SE df lower.CL upper.CL<br>LAR1 0.07787 0.375 143 -0.663 0.818<br>LAR2 1.34053 0.375 143 0.600 2.081<br><br>Tissue = Stem:<br>Genes emmean SE df lower.CL upper.CL<br>LAR1 0.07063 0.324 143 -0.571 0.712<br>LAR2 4.61635 0.324 143 3.975 5.258<br><br>Tissue = Zygotic embryo:<br>Genes emmean SE df lower.CL upper.CL<br>LAR1 0.00166 0.375 143 -0.739 0.742<br>LAR2 0.24020 0.375 143 -0.500 0.981 | Tissue = Needles:<br>contrast estimate SE df t.ratio p.value<br>LAR1 - LAR2 -3.690 0.253 106 -14.601 <.0001<br><br>Tissue = Root:<br>contrast estimate SE df t.ratio p.value<br>LAR1 - LAR2 -1.034 0.126 106 -8.186 <.0001<br><br>Tissue = shoot:<br>contrast estimate SE df t.ratio p.value<br>LAR1 - LAR2 -2.394 0.187 106 -12.828 <.0001<br><br>Tissue = Somatic embryo:<br>contrast estimate SE df t.ratio p.value<br>LAR1 - LAR2 -1.263 0.292 106 -4.327 <.0001<br><br>Tissue = Stem:<br>contrast estimate SE df t.ratio p.value<br>LAR1 - LAR2 -4.546 0.253 106 -17.989 <.0001<br><br>Tissue = Zygotic embryo:<br>contrast estimate SE df t.ratio p.value<br>LAR1 - LAR2 -0.239 0.292 106 -0.818 0.4155<br><br>Degrees-of-freedom method: kenward-roger<br>Confidence level used: 0.95 |

### *Picea sitchensis*

Linear mixed model fit by REML. t-tests use Satterthwaite's method ['lmerModLmerTest']
Formula: TPM_log ~ Genes * tissue + (1 | Sample)
Data: data_long

REML criterion at convergence: 308.8

Scaled residuals:

| Min | 1Q | Median | 3Q | Max |
|---|---|---|---|---|
| -2.8217 | -0.41947 | -0.00553 | 0.41571 | 2.8217 |

Random effects:

| Groups | Name | Variance | Std.Dev. |
|---|---|---|---|
| Sample | (Intercept) | 0.1518 | 0.3897 |
| Residual | 0.2081 | 0.4561 | |
| Number of obs: | 178 | groups: | Sample, 89 |

Fixed effects:

| | *Estimate* | *Std. Error* | *df* | *T-value* | *Pr(>\|t\|)* | |
|---|---|---|---|---|---|---|
| *(Intercept)* | *4.87926* | *0.59992* | *139.22303* | *8.133* | *2.07E-13* | *** |
| *GenesLAR2* | *-0.37136* | *0.64509* | *82* | *-0.576* | *0.566413* | |
| *tissueCortex* | *-1.7998* | *0.61636* | *139.22303* | *-2.92* | *0.004083* | ** |
| *tissueEndosperm* | *-3.28643* | *0.73474* | *139.22303* | *-4.473* | *1.59E-05* | *** |
| *tissueNeedle* | *0.35175* | *0.73474* | *139.22303* | *0.479* | *0.632872* | |
| *tissueSclereid* | *-2.19898* | *0.61636* | *139.22303* | *-3.568* | *0.000494* | *** |
| *tissueShoot tip* | *-1.30728* | *0.61229* | *139.22303* | *-2.135* | *0.034505* | * |
| *tissueStem* | *-0.87324* | *0.61229* | *139.22303* | *-1.426* | *0.156051* | |
| *GenesLAR2:tissueCortex* | *-0.46952* | *0.66277* | *82* | *-0.708* | *0.48069* | |
| *GenesLAR2:tissueEndosperm* | *-0.99933* | *0.79007* | *82* | *-1.265* | *0.209507* | |
| *GenesLAR2:tissueNeedle* | *-2.84911* | *0.79007* | *82* | *-3.606* | *0.000533* | *** |
| *GenesLAR2:tissueSclereid* | *0.62355* | *0.66277* | *82* | *0.941* | *0.349556* | |
| *GenesLAR2:tissueShoot tip* | *1.11278* | *0.6584* | *82* | *1.69* | *0.094798* | |
| *GenesLAR2:tissueStem* | *-0.03206* | *0.6584* | *82* | *-0.049* | *0.961284* | |

Signif. codes: 0 '***' 0.001 '**' 0.01 '*' 0.05 '.' 0.1 ' ' 1

**Table 4S.** Estimated marginal means and pairwise comparison for the species *Picea sitchensis*. Degrees-of-freedom method: kenward-roger. Confidence level used: 0.95.

| ***Estimated Marginal Means*** | ***Contrasts*** |
|---|---|
| tissue = Bark:<br>Genes emmean SE df lower.CL upper.CL<br>LAR1 4.879 0.600 139 3.693 6.07 | tissue = Bark:<br>contrast estimate SE df t.ratio p.value<br>LAR1 - LAR2 0.371 0.645 82 0.576 0.5664 |

```
 LAR2   4.508 0.600 139    3.322     5.69

tissue = Cortex:
 Genes emmean    SE  df lower.CL upper.CL
 LAR1   3.079 0.141 139    2.800     3.36
 LAR2   2.239 0.141 139    1.959     2.52

tissue = Endosperm:
 Genes emmean    SE  df lower.CL upper.CL
 LAR1   1.593 0.424 139    0.754     2.43
 LAR2   0.222 0.424 139   -0.617     1.06

tissue = Needle:
 Genes emmean    SE  df lower.CL upper.CL
 LAR1   5.231 0.424 139    4.392     6.07
 LAR2   2.011 0.424 139    1.172     2.85
tissue = Sclereid:
 Genes emmean    SE  df lower.CL upper.CL
 LAR1   2.680 0.141 139    2.401     2.96
 LAR2   2.932 0.141 139    2.653     3.21
tissue = Shoot tip:
 Genes emmean    SE  df lower.CL upper.CL
 LAR1   3.572 0.122 139    3.330     3.81
 LAR2   4.313 0.122 139    4.071     4.56
tissue = Stem:
 Genes emmean    SE  df lower.CL upper.CL
 LAR1   4.006 0.122 139    3.764     4.25
 LAR2   3.603 0.122 139    3.360     3.84
```

```
tissue = Cortex:
 contrast    estimate    SE df t.ratio p.value
 LAR1 - LAR2    0.841 0.152 82   5.530  <.0001

tissue = Endosperm:
 contrast    estimate    SE df t.ratio p.value
 LAR1 - LAR2    1.371 0.456 82   3.005  0.0035

tissue = Needle:
 contrast    estimate    SE df t.ratio p.value
 LAR1 - LAR2    3.220 0.456 82   7.060  <.0001

tissue = Sclereid:
 contrast    estimate    SE df t.ratio p.value
 LAR1 - LAR2   -0.252 0.152 82  -1.659  0.1010

tissue = Shoot tip:
 contrast    estimate    SE df t.ratio p.value
 LAR1 - LAR2   -0.741 0.132 82  -5.631  <.0001

tissue = Stem:
 contrast    estimate    SE df t.ratio p.value
 LAR1 - LAR2    0.403 0.132 82   3.064  0.0030

Degrees-of-freedom method: kenward-roger
Confidence level used: 0.95
```

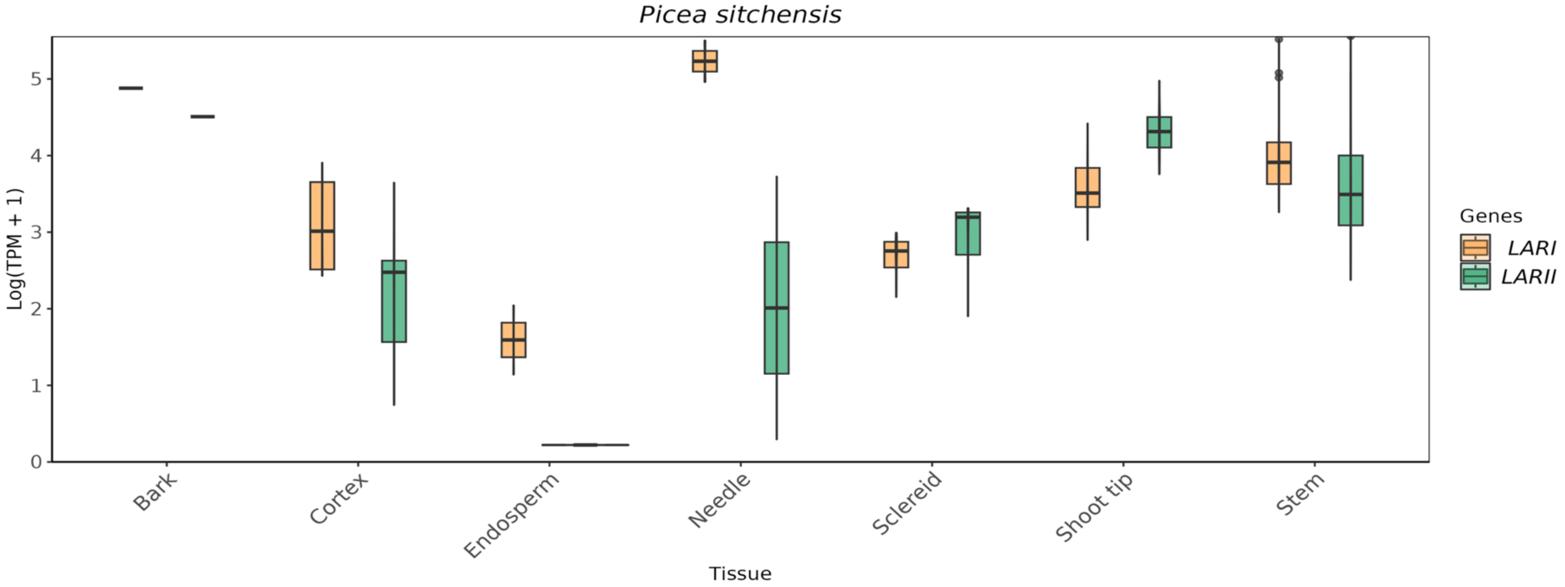

**Figure 1S.** Comparison of the expression of *LARI* (orange) and *LARII* (green) genes in the species *Picea sitchensis*

#### *Thuja plicata*

Linear mixed model fit by REML. t-tests use Satterthwaite's method ['lmerModLmerTest']
Formula: TPM_log ~ Genes * tissue + (1 | Sample)
Data: data_long

REML criterion at convergence: 296.6

Scaled residuals:

| Min | 1Q | Median | 3Q | Max |
|---|---|---|---|---|
| -5.1305 | -0.2767 | -0.0173 | 0.2678 | 5.7278 |

Random effects:

| Groups | Name | Variance | Std.Dev. |
|---|---|---|---|
| Sample | (Intercept) | 0.007017 | 0.08377 |
| Residual | 0.075059 | 0.27397 | |
| Number of obs: | 664 | groups: | Sample, 83 |

Fixed effects:

| | *Estimate* | *Std. Error* | *df* | *T-value* | *Pr(>\|t\|)* | |
|---|---|---|---|---|---|---|
| *(Intercept)* | *4.34E-14* | *7.40E-02* | *6.01E+02* | *0* | *1* | |
| *GenesLAR1b* | *-3.39E-14* | *1.00E-01* | *5.53E+02* | *0* | *1* | |
| *GenesLAR1c* | *4.66E-02* | *1.00E-01* | *5.53E+02* | *0.466* | *0.64142* | |
| *GenesLAR1d* | *7.15E-02* | *1.00E-01* | *5.53E+02* | *0.715* | *0.47484* | |
| *GenesLAR1e* | *-3.49E-14* | *1.00E-01* | *5.53E+02* | *0* | *1* | |
| *GenesLAR1f* | *5.63E-03* | *1.00E-01* | *5.53E+02* | *0.056* | *0.95515* | |
| *GenesLAR2a* | *3.26E+00* | *1.00E-01* | *5.53E+02* | *32.538* | *< 2.00E-16* | *** |
| *GenesLAR2b* | *1.61E+00* | *1.00E-01* | *5.53E+02* | *16.057* | *< 2.00E-16* | *** |
| *tissueFoliage* | *8.01E-04* | *8.97E-02* | *6.01E+02* | *0.009* | *0.99288* | |
| *tissueWhole seedling* | *6.07E-01* | *1.03E-01* | *6.01E+02* | *5.89* | *6.42E-09* | *** |
| *tissueWood core* | *-4.50E-14* | *9.79E-02* | *6.01E+02* | *0* | *1* | |
| *GenesLAR1b:tissueFoliage* | *3.27E-04* | *1.21E-01* | *5.53E+02* | *0.003* | *0.99785* | |
| *GenesLAR1c:tissueFoliage* | *8.89E-01* | *1.21E-01* | *5.53E+02* | *7.334* | *8.02E-13* | *** |
| *GenesLAR1d:tissueFoliage* | *7.64E-01* | *1.21E-01* | *5.53E+02* | *6.304* | *5.93E-10* | *** |
| *GenesLAR1e:tissueFoliage* | *1.87E-02* | *1.21E-01* | *5.53E+02* | *0.154* | *0.87756* | |
| *GenesLAR1f:tissueFoliage* | *5.98E-02* | *1.21E-01* | *5.53E+02* | *0.494* | *0.62182* | |
| *GenesLAR2a:tissueFoliage* | *8.04E-01* | *1.21E-01* | *5.53E+02* | *6.628* | *8.08E-11* | *** |
| *GenesLAR2b:tissueFoliage* | *2.30E+00* | *1.21E-01* | *5.53E+02* | *18.937* | *< 2.00E-16* | *** |
| *GenesLAR1b:tissueWhole seedling* | *-6.04E-01* | *1.39E-01* | *5.53E+02* | *-4.335* | *1.73E-05* | *** |
| *GenesLAR1c:tissueWhole seedling* | *3.10E-01* | *1.39E-01* | *5.53E+02* | *2.223* | *0.02664* | * |
| *GenesLAR1d:tissueWhole seedling* | *-2.18E-01* | *1.39E-01* | *5.53E+02* | *-1.566* | *0.118* | |
| *GenesLAR1e:tissueWhole seedling* | *2.37E-01* | *1.39E-01* | *5.53E+02* | *1.7* | *0.08969* | . |
| *GenesLAR1f:tissueWhole seedling* | *4.30E-01* | *1.39E-01* | *5.53E+02* | *3.087* | *0.00212* | ** |
| *GenesLAR2a:tissueWhole seedling* | *-1.93E+00* | *1.39E-01* | *5.53E+02* | *-13.838* | *< 2.00E-16* | *** |
| *GenesLAR2b:tissueWhole seedling* | *6.23E-01* | *1.39E-01* | *5.53E+02* | *4.47* | *9.49E-06* | *** |
| *GenesLAR1b:tissueWood core* | *3.55E-14* | *1.32E-01* | *5.53E+02* | *0* | *1* | |
| *GenesLAR1c:tissueWood core* | *-4.66E-02* | *1.32E-01* | *5.53E+02* | *-0.352* | *0.72479* | |
| *GenesLAR1d:tissueWood core* | *-1.59E-02* | *1.32E-01* | *5.53E+02* | *-0.12* | *0.90469* | |
| *GenesLAR1e:tissueWood core* | *2.95E-03* | *1.32E-01* | *5.53E+02* | *0.022* | *0.98223* | |
| *GenesLAR1f:tissueWood core* | *-3.51E-03* | *1.32E-01* | *5.53E+02* | *-0.027* | *0.97886* | |
| *GenesLAR2a:tissueWood core* | *-1.69E+00* | *1.32E-01* | *5.53E+02* | *-12.752* | *< 2.00E-16* | *** |
| *GenesLAR2b:tissueWood core* | *3.33E-01* | *1.32E-01* | *5.53E+02* | *2.518* | *0.0121* | * |

Signif. codes:  0 '***' 0.001 '**' 0.01 '*' 0.05 '.' 0.1 ' ' 1

**Table 5S.** Estimated marginal means and pairwise comparison for the species *Thuja plicata*. Degrees-of-freedom method: kenward-roge. P value adjustment: bonferroni method for 28 tests.

| ***Estimated Marginal Means*** | ***Contrasts*** |
|---|---|
| tissue = Cambium: | tissue = Cambium: |
| Genes emmean SE df lower.CL upper.CL | contrast estimate SE df t.ratio p.value |
| LAR1a 0.000000 0.0740 601 -0.1453 0.145 | LAR1a - LAR1b 0.000000 0.1000 553 0.000 1.0000 |
| LAR1b 0.000000 0.0740 601 -0.1453 0.145 | LAR1a - LAR1c -0.046616 0.1000 553 -0.466 1.0000 |
| LAR1c 0.046616 0.0740 601 -0.0987 0.192 | LAR1a - LAR1d -0.071541 0.1000 553 -0.715 1.0000 |
| LAR1d 0.071541 0.0740 601 -0.0737 0.217 | LAR1a - LAR1e 0.000000 0.1000 553 0.000 1.0000 |
| LAR1e 0.000000 0.0740 601 -0.1453 0.145 | LAR1a - LAR1f -0.005629 0.1000 553 -0.056 1.0000 |
| LAR1f 0.005629 0.0740 601 -0.1396 0.151 | LAR1a - LAR2a -3.255100 0.1000 553 -32.538 <.0001 |
| LAR2a 3.255100 0.0740 601 3.1098 3.400 | LAR1a - LAR2b -1.606325 0.1000 553 -16.057 <.0001 |
| LAR2b 1.606325 0.0740 601 1.4611 1.752 | LAR1b - LAR1c -0.046616 0.1000 553 -0.466 1.0000 |
| | LAR1b - LAR1d -0.071541 0.1000 553 -0.715 1.0000 |
| tissue = Foliage: | LAR1b - LAR1e 0.000000 0.1000 553 0.000 1.0000 |
| Genes emmean SE df lower.CL upper.CL | LAR1b - LAR1f -0.005629 0.1000 553 -0.056 1.0000 |
| LAR1a 0.000801 0.0506 601 -0.0987 0.100 | LAR1b - LAR2a -3.255100 0.1000 553 -32.538 <.0001 |
| LAR1b 0.001128 0.0506 601 -0.0983 0.101 | LAR1b - LAR2b -1.606325 0.1000 553 -16.057 <.0001 |
| LAR1c 0.936533 0.0506 601 0.8371 1.036 | LAR1c - LAR1d -0.024924 0.1000 553 -0.249 1.0000 |
| LAR1d 0.836638 0.0506 601 0.7372 0.936 | LAR1c - LAR1e 0.046616 0.1000 553 0.466 1.0000 |
| LAR1e 0.019487 0.0506 601 -0.0800 0.119 | LAR1c - LAR1f 0.040988 0.1000 553 0.410 1.0000 |
| LAR1f 0.066268 0.0506 601 -0.0332 0.166 | LAR1c - LAR2a -3.208484 0.1000 553 -32.072 <.0001 |
| LAR2a 4.059520 0.0506 601 3.9601 4.159 | LAR1c - LAR2b -1.559709 0.1000 553 -15.591 <.0001 |
| LAR2b 3.903093 0.0506 601 3.8036 4.003 | LAR1d - LAR1e 0.071541 0.1000 553 0.715 1.0000 |
| | LAR1d - LAR1f 0.065912 0.1000 553 0.659 1.0000 |
| tissue = Whole seedling: | LAR1d - LAR2a -3.183560 0.1000 553 -31.823 <.0001 |
| Genes emmean SE df lower.CL upper.CL | LAR1d - LAR2b -1.534785 0.1000 553 -15.342 <.0001 |
| LAR1a 0.606477 0.0716 601 0.4658 0.747 | LAR1e - LAR1f -0.005629 0.1000 553 -0.056 1.0000 |
| LAR1b 0.002840 0.0716 601 -0.1378 0.144 | LAR1e - LAR2a -3.255100 0.1000 553 -32.538 <.0001 |
| LAR1c 0.962611 0.0716 601 0.8219 1.103 | LAR1e - LAR2b -1.606325 0.1000 553 -16.057 <.0001 |
| LAR1d 0.460002 0.0716 601 0.3193 0.601 | LAR1f - LAR2a -3.249472 0.1000 553 -32.482 <.0001 |
| LAR1e 0.843202 0.0716 601 0.7025 0.984 | LAR1f - LAR2b -1.600697 0.1000 553 -16.001 <.0001 |
| LAR1f 1.041985 0.0716 601 0.9013 1.183 | LAR2a - LAR2b 1.648775 0.1000 553 16.481 <.0001 |
| LAR2a 1.934677 0.0716 601 1.7940 2.075 | tissue = Foliage: |
| LAR2b 2.835261 0.0716 601 2.6946 2.976 | contrast estimate SE df t.ratio p.value |
| | LAR1a - LAR1b -0.000327 0.0685 553 -0.005 1.0000 |
| tissue = Wood core: | LAR1a - LAR1c -0.935732 0.0685 553 -13.662 <.0001 |
| Genes emmean SE df lower.CL upper.CL | LAR1a - LAR1d -0.835837 0.0685 553 -12.203 <.0001 |
| LAR1a 0.000000 0.0641 601 -0.1258 0.126 | LAR1a - LAR1e -0.018687 0.0685 553 -0.273 1.0000 |
| LAR1b 0.000000 0.0641 601 -0.1258 0.126 | LAR1a - LAR1f -0.065467 0.0685 553 -0.956 1.0000 |
| LAR1c 0.000000 0.0641 601 -0.1258 0.126 | LAR1a - LAR2a -4.058719 0.0685 553 -59.258 <.0001 |
| LAR1d 0.055687 0.0641 601 -0.0701 0.181 | LAR1a - LAR2b -3.902292 0.0685 553 -56.974 <.0001 |
| LAR1e 0.002949 0.0641 601 -0.1229 0.129 | LAR1b - LAR1c -0.935405 0.0685 553 -13.657 <.0001 |
| LAR1f 0.002121 0.0641 601 -0.1237 0.128 | LAR1b - LAR1d -0.835510 0.0685 553 -12.199 <.0001 |
| LAR2a 1.567546 0.0641 601 1.4417 1.693 | LAR1b - LAR1e -0.018359 0.0685 553 -0.268 1.0000 |
| LAR2b 1.939493 0.0641 601 1.8137 2.065 | LAR1b - LAR1f -0.065140 0.0685 553 -0.951 1.0000 |
| | LAR1b - LAR2a -4.058392 0.0685 553 -59.253 <.0001 |

```
 LAR1b - LAR2b -3.901965 0.0685 553 -56.969  <.0001
 LAR1c - LAR1d  0.099896 0.0685 553   1.458  1.0000
 LAR1c - LAR1e  0.917046 0.0685 553  13.389  <.0001
 LAR1c - LAR1f  0.870265 0.0685 553  12.706  <.0001
 LAR1c - LAR2a -3.122987 0.0685 553 -45.596  <.0001
 LAR1c - LAR2b -2.966560 0.0685 553 -43.312  <.0001
 LAR1d - LAR1e  0.817150 0.0685 553  11.931  <.0001
 LAR1d - LAR1f  0.770370 0.0685 553  11.248  <.0001
 LAR1d - LAR2a -3.222882 0.0685 553 -47.055  <.0001
 LAR1d - LAR2b -3.066455 0.0685 553 -44.771  <.0001
 LAR1e - LAR1f -0.046780 0.0685 553  -0.683  1.0000
 LAR1e - LAR2a -4.040033 0.0685 553 -58.985  <.0001
 LAR1e - LAR2b -3.883605 0.0685 553 -56.701  <.0001
 LAR1f - LAR2a -3.993252 0.0685 553 -58.302  <.0001
 LAR1f - LAR2b -3.836825 0.0685 553 -56.018  <.0001
 LAR2a - LAR2b  0.156427 0.0685 553   2.284  0.6372
tissue = Whole seedling:
 contrast       estimate     SE  df t.ratio p.value
 LAR1a - LAR1b  0.603637 0.0969 553   6.232  <.0001
 LAR1a - LAR1c -0.356133 0.0969 553  -3.677  0.0073
 LAR1a - LAR1d  0.146475 0.0969 553   1.512  1.0000
 LAR1a - LAR1e -0.236725 0.0969 553  -2.444  0.4155
 LAR1a - LAR1f -0.435508 0.0969 553  -4.496  0.0002
 LAR1a - LAR2a -1.328200 0.0969 553 -13.712  <.0001
 LAR1a - LAR2b -2.228784 0.0969 553 -23.010  <.0001
 LAR1b - LAR1c -0.959770 0.0969 553  -9.909  <.0001
 LAR1b - LAR1d -0.457162 0.0969 553  -4.720  0.0001
 LAR1b - LAR1e -0.840362 0.0969 553  -8.676  <.0001
 LAR1b - LAR1f -1.039144 0.0969 553 -10.728  <.0001
 LAR1b - LAR2a -1.931837 0.0969 553 -19.944  <.0001
 LAR1b - LAR2b -2.832421 0.0969 553 -29.242  <.0001
 LAR1c - LAR1d  0.502608 0.0969 553   5.189  <.0001
 LAR1c - LAR1e  0.119408 0.0969 553   1.233  1.0000
 LAR1c - LAR1f -0.079374 0.0969 553  -0.819  1.0000
 LAR1c - LAR2a -0.972066 0.0969 553 -10.035  <.0001
 LAR1c - LAR2b -1.872651 0.0969 553 -19.333  <.0001
 LAR1d - LAR1e -0.383200 0.0969 553  -3.956  0.0024
 LAR1d - LAR1f -0.581983 0.0969 553  -6.008  <.0001
 LAR1d - LAR2a -1.474675 0.0969 553 -15.224  <.0001
 LAR1d - LAR2b -2.375259 0.0969 553 -24.522  <.0001
 LAR1e - LAR1f -0.198783 0.0969 553  -2.052  1.0000
 LAR1e - LAR2a -1.091475 0.0969 553 -11.268  <.0001
 LAR1e - LAR2b -1.992059 0.0969 553 -20.566  <.0001
 LAR1f - LAR2a -0.892692 0.0969 553  -9.216  <.0001
 LAR1f - LAR2b -1.793277 0.0969 553 -18.514  <.0001
 LAR2a - LAR2b -0.900584 0.0969 553  -9.298  <.0001
tissue = Wood core:
 contrast       estimate     SE  df t.ratio p.value
```

| | |
|---|---|
| | LAR1a - LAR1b 0.000000 0.0866 553 0.000 1.0000<br>LAR1a - LAR1c 0.000000 0.0866 553 0.000 1.0000<br>LAR1a - LAR1d -0.055687 0.0866 553 -0.643 1.0000<br>LAR1a - LAR1e -0.002949 0.0866 553 -0.034 1.0000<br>LAR1a - LAR1f -0.002121 0.0866 553 -0.024 1.0000<br>LAR1a - LAR2a -1.567546 0.0866 553 -18.093 <.0001<br>LAR1a - LAR2b -1.939493 0.0866 553 -22.386 <.0001<br>LAR1b - LAR1c 0.000000 0.0866 553 0.000 1.0000<br>LAR1b - LAR1d -0.055687 0.0866 553 -0.643 1.0000<br>LAR1b - LAR1e -0.002949 0.0866 553 -0.034 1.0000<br>LAR1b - LAR1f -0.002121 0.0866 553 -0.024 1.0000<br>LAR1b - LAR2a -1.567546 0.0866 553 -18.093 <.0001<br>LAR1b - LAR2b -1.939493 0.0866 553 -22.386 <.0001<br>LAR1c - LAR1d -0.055687 0.0866 553 -0.643 1.0000<br>LAR1c - LAR1e -0.002949 0.0866 553 -0.034 1.0000<br>LAR1c - LAR1f -0.002121 0.0866 553 -0.024 1.0000<br>LAR1c - LAR2a -1.567546 0.0866 553 -18.093 <.0001<br>LAR1c - LAR2b -1.939493 0.0866 553 -22.386 <.0001<br>LAR1d - LAR1e 0.052738 0.0866 553 0.609 1.0000<br>LAR1d - LAR1f 0.053566 0.0866 553 0.618 1.0000<br>LAR1d - LAR2a -1.511859 0.0866 553 -17.451 <.0001<br>LAR1d - LAR2b -1.883806 0.0866 553 -21.744 <.0001<br>LAR1e - LAR1f 0.000828 0.0866 553 0.010 1.0000<br>LAR1e - LAR2a -1.564597 0.0866 553 -18.059 <.0001<br>LAR1e - LAR2b -1.936544 0.0866 553 -22.352 <.0001<br>LAR1f - LAR2a -1.565425 0.0866 553 -18.069 <.0001<br>LAR1f - LAR2b -1.937372 0.0866 553 -22.362 <.0001<br>LAR2a - LAR2b -0.371947 0.0866 553 -4.293 0.0006 |

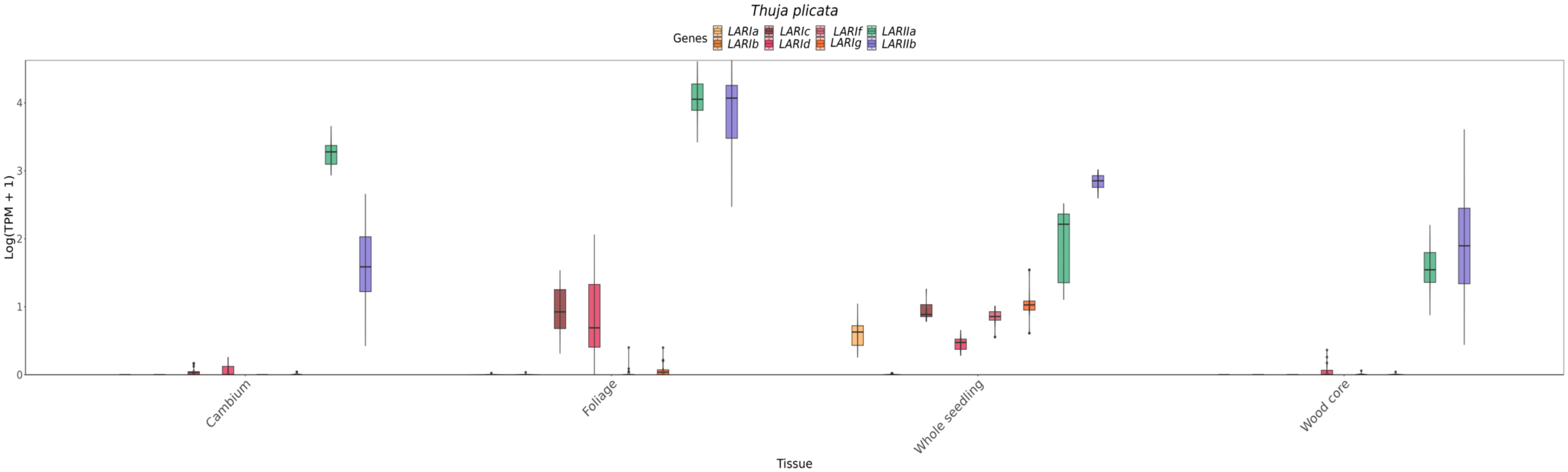

**Figure 2S.** Comparison of the expression of *LARIa* (yellow), *LARIb* (coral), *LARIc* (brown), *LARId* (pink), *LARIf* (red), *LARIg* (orange), *LARIIa* (green) and *LARIIb* (lila) genes in the species *Thuja plicata.*

### Dicotyledon species

#### *Vitis vinifera*

Linear mixed model fit by REML. t-tests use Satterthwaite's method ['lmerModLmerTest']
Formula: TPM_log ~ Genes * tissue + (1 | Sample)
Data: data_long

REML criterion at convergence: 2277.9

Scaled residuals:

| Min | 1Q | Median | 3Q | Max |
|---|---|---|---|---|
| -2.52487 | -0.66092 | -0.08172 | 0.50523 | 3.04447 |

Random effects:

| Groups | Name | Variance | Std.Dev. |
|---|---|---|---|
| Sample | (Intercept) | 0.3433 | 0.5859 |
| Residual | 1.4538 | 1.2058 | |
| Number of obs: | 674 | groups: | Sample, 337 |

Fixed effects:

| | Estimate | Std. Error | df | T-value | Pr(>\|t\|) | |
|---|---|---|---|---|---|---|
| (Intercept) | 1.49268 | 0.10374 | 630.97507 | 14.389 | < 2E-016 | *** |
| GenesLAR2 | 0.7286 | 0.13195 | 326.99998 | 5.522 | 6.85E-08 | *** |
| tissueCallus | -1.46912 | 0.67827 | 630.97507 | -2.166 | 0.03069 | * |
| tissueFlower | 1.5704 | 0.40065 | 630.97507 | 3.92 | 9.84E-05 | *** |
| tissueGrapevine | 2.32336 | 0.40065 | 630.97507 | 5.799 | 1.06E-08 | *** |
| tissueInflorescence | 0.97675 | 0.40065 | 630.97507 | 2.438 | 0.01505 | * |
| tissueLeaf | 0.31846 | 0.16696 | 630.97507 | 1.907 | 0.05693 | . |
| tissueOvule | 2.23736 | 0.45874 | 630.97507 | 4.877 | 1.36E-06 | *** |
| tissuePlant | 0.82036 | 0.55703 | 630.97507 | 1.473 | 0.14132 | |
| tissueSeedling | 2.26516 | 0.95359 | 630.97507 | 2.375 | 0.01783 | * |
| tissueStem | 0.79727 | 0.48519 | 630.97507 | 1.643 | 0.10084 | |
| GenesLAR2:tissueCallus | -0.7298 | 0.86275 | 326.99998 | -0.846 | 0.39822 | |
| GenesLAR2:tissueFlower | -0.8013 | 0.50963 | 326.99998 | -1.572 | 0.11684 | |
| GenesLAR2:tissueGrapevine | -0.9581 | 0.50963 | 326.99998 | -1.88 | 0.061 | . |
| GenesLAR2:tissueInflorescence | -0.61914 | 0.50963 | 326.99998 | -1.215 | 0.22528 | |
| GenesLAR2:tissueLeaf | 0.67119 | 0.21238 | 326.99998 | 3.16 | 0.00172 | ** |
| GenesLAR2:tissueOvule | -0.47409 | 0.58351 | 326.99998 | -0.812 | 0.41711 | |
| GenesLAR2:tissuePlant | -1.02363 | 0.70854 | 326.99998 | -1.445 | 0.1495 | |

| | | | | | | |
|---|---|---|---|---|---|---|
| *GenesLAR2:tissueSeedling* | *-2.09943* | *1.21296* | *326.99998* | *-1.731* | *0.08442* | *.* |
| *GenesLAR2:tissueStem* | *0.07115* | *0.61715* | *326.99998* | *0.115* | *0.90829* | |

Signif. codes: 0 ‘***’ 0.001 ‘**’ 0.01 ‘*’ 0.05 ‘.’ 0.1 ‘ ’ 1

**Table 6S.** Estimated marginal means and pairwise comparison for the species *Vitis vinifera*. Degrees-of-freedom method: kenward-roger. Confidence level used: 0.95.

| ***Estimated Marginal Means*** | ***Contrasts*** |
|---|---|
| tissue = Berry:<br>Genes emmean SE df lower.CL upper.CL<br>LAR1 1.4927 0.104 631 1.289 1.70<br>LAR2 2.2213 0.104 631 2.018 2.42<br><br>tissue = Callus:<br>Genes emmean SE df lower.CL upper.CL<br>LAR1 0.0236 0.670 631 -1.293 1.34<br>LAR2 0.0224 0.670 631 -1.294 1.34<br><br>tissue = Flower:<br>Genes emmean SE df lower.CL upper.CL<br>LAR1 3.0631 0.387 631 2.303 3.82<br>LAR2 2.9904 0.387 631 2.230 3.75<br><br>tissue = Grapevine:<br>Genes emmean SE df lower.CL upper.CL<br>LAR1 3.8160 0.387 631 3.056 4.58<br>LAR2 3.5865 0.387 631 2.827 4.35<br><br>tissue = Inflorescence:<br>Genes emmean SE df lower.CL upper.CL<br>LAR1 2.4694 0.387 631 1.709 3.23<br>LAR2 2.5789 0.387 631 1.819 3.34<br><br>tissue = Leaf:<br>Genes emmean SE df lower.CL upper.CL<br>LAR1 1.8111 0.131 631 1.554 2.07<br>LAR2 3.2109 0.131 631 2.954 3.47<br><br>tissue = Ovule:<br>Genes emmean SE df lower.CL upper.CL<br>LAR1 3.7300 0.447 631 2.853 4.61<br>LAR2 3.9845 0.447 631 3.107 4.86<br><br>tissue = Plant:<br>Genes emmean SE df lower.CL upper.CL<br>LAR1 2.3130 0.547 631 1.238 3.39<br>LAR2 2.0180 0.547 631 0.943 3.09<br><br>tissue = Seedling: | tissue = Berry:<br>contrast estimate SE df t.ratio p.value<br>LAR1 - LAR2 -0.72860 0.132 327 -5.522 <.0001<br><br>tissue = Callus:<br>contrast estimate SE df t.ratio p.value<br>LAR1 - LAR2 0.00121 0.853 327 0.001 0.9989<br><br>tissue = Flower:<br>contrast estimate SE df t.ratio p.value<br>LAR1 - LAR2 0.07270 0.492 327 0.148 0.8827<br><br>tissue = Grapevine:<br>contrast estimate SE df t.ratio p.value<br>LAR1 - LAR2 0.22951 0.492 327 0.466 0.6414<br><br>tissue = Inflorescence:<br>contrast estimate SE df t.ratio p.value<br>LAR1 - LAR2 -0.10946 0.492 327 -0.222 0.8242<br><br>tissue = Leaf:<br>contrast estimate SE df t.ratio p.value<br>LAR1 - LAR2 -1.39979 0.166 327 -8.412 <.0001<br><br>tissue = Ovule:<br>contrast estimate SE df t.ratio p.value<br>LAR1 - LAR2 -0.25451 0.568 327 -0.448 0.6546<br><br>tissue = Plant:<br>contrast estimate SE df t.ratio p.value<br>LAR1 - LAR2 0.29504 0.696 327 0.424 0.6720<br><br>tissue = Seedling:<br>contrast estimate SE df t.ratio p.value<br>LAR1 - LAR2 1.37084 1.210 327 1.137 0.2564<br><br>tissue = Stem:<br>contrast estimate SE df t.ratio p.value<br>LAR1 - LAR2 -0.79975 0.603 327 -1.327 0.1856<br><br>Degrees-of-freedom method: kenward-roger |

| |
|---|
| Genes emmean SE df lower.CL upper.CL<br>LAR1 3.7578 0.948 631 1.896 5.62<br>LAR2 2.3870 0.948 631 0.526 4.25<br><br>tissue = Stem:<br>Genes emmean SE df lower.CL upper.CL<br>LAR1 2.2900 0.474 631 1.359 3.22<br>LAR2 3.0897 0.474 631 2.159 4.02<br><br>Degrees-of-freedom method: kenward-roger<br>Confidence level used: 0.95 |

### *Theobroma cacao*

Linear mixed model fit by REML. t-tests use Satterthwaite's method ['lmerModLmerTest']
Formula: TPM_log ~ Genes * tissue + (1 | Sample)
Data: data_long

REML criterion at convergence: 2160.3

Scaled residuals:

| Min | 1Q | Median | 3Q | Max |
|---|---|---|---|---|
| -2.41237 | -0.50204 | 0.01226 | 0.51105 | 1.94826 |

Random effects:

| Groups | Name | Variance | Std.Dev. |
|---|---|---|---|
| Sample | (Intercept) | 1.2945 | 1.1378 |
| Residual | 0.4684 | 0.6844 | |
| Number of obs: | 716 | groups: | Sample, 358 |

Fixed effects:

| | *Estimate* | *Std. Error* | *df* | *T-value* | *Pr(>\|t\|)* | |
|---|---|---|---|---|---|---|
| *(Intercept)* | 3.92368 | 0.22443 | 454.78463 | 17.483 | <2E-016 | *** |
| *GenesLAR2* | -0.99649 | 0.16361 | 350.00001 | -6.091 | 2.96E-09 | *** |
| *tissueInflorescence* | 0.53744 | 0.47609 | 454.78463 | 1.129 | 0.25955 | |
| *tissueLeaf* | -0.96882 | 0.23975 | 454.78463 | -4.041 | 6.25E-05 | *** |
| *tissueMix-leaf-flower* | 0.31182 | 0.37218 | 454.78463 | 0.838 | 0.40257 | |
| *tissuePistil* | 0.1097 | 0.47609 | 454.78463 | 0.23 | 0.81788 | |
| *tissuePod* | -2.65025 | 0.44416 | 454.78463 | -5.967 | 4.87E-09 | *** |
| *tissueSeedling* | -0.05375 | 0.47609 | 454.78463 | -0.113 | 0.91015 | |
| *tissueYoung red leaves* | -0.45728 | 0.43126 | 454.78463 | -1.06 | 0.28955 | |
| *GenesLAR2:tissueInflorescence* | 0.25293 | 0.34706 | 350.00001 | 0.729 | 0.46662 | |
| *GenesLAR2:tissueLeaf* | 0.57051 | 0.17477 | 350.00001 | 3.264 | 0.00121 | ** |

| | | | | | | |
|---|---|---|---|---|---|---|
| *GenesLAR2:tissueMix-leaf-flower* | 0.37982 | 0.27131 | 350.00001 | 1.4 | 0.16242 | |
| *GenesLAR2:tissuePistil* | 2.27943 | 0.34706 | 350.00001 | 6.568 | 1.85E-10 | *** |
| *GenesLAR2:tissuePod* | 3.70218 | 0.32379 | 350.00001 | 11.434 | <2E-016 | *** |
| *GenesLAR2:tissueSeedling* | 0.27493 | 0.34706 | 350.00001 | 0.792 | 0.42881 | |
| *GenesLAR2:tissueYoung red leaves* | -0.08063 | 0.31438 | 350.00001 | -0.256 | 0.79774 | |

Signif. codes: 0 '***' 0.001 '**' 0.01 '*' 0.05 '.' 0.1 ' ' 1

**Table 7S.** Estimated marginal means and pairwise comparison for the species *Theobroma cacao*. Degrees-of-freedom method: kenward-roger. Confidence level used: 0.95.

| ***Estimated Marginal Means*** | ***Contrasts*** |
|---|---|
| tissue = Cells:<br>Genes emmean SE df lower.CL upper.CL<br>LAR1 3.92 0.2240 455 3.48 4.36<br>LAR2 2.93 0.2240 455 2.49 3.37<br>tissue = Inflorescence:<br>Genes emmean SE df lower.CL upper.CL<br>LAR1 4.46 0.4200 455 3.64 5.29<br>LAR2 3.72 0.4200 455 2.89 4.54<br>tissue = Leaf:<br>Genes emmean SE df lower.CL upper.CL<br>LAR1 2.95 0.0843 455 2.79 3.12<br>LAR2 2.53 0.0843 455 2.36 2.69<br>tissue = Mix-leaf-flower:<br>Genes emmean SE df lower.CL upper.CL<br>LAR1 4.24 0.2970 455 3.65 4.82<br>LAR2 3.62 0.2970 455 3.04 4.20<br>tissue = Pistil:<br>Genes emmean SE df lower.CL upper.CL<br>LAR1 4.03 0.4200 455 3.21 4.86<br>LAR2 5.32 0.4200 455 4.49 6.14<br>tissue = Pod:<br>Genes emmean SE df lower.CL upper.CL<br>LAR1 1.27 0.3830 455 0.52 2.03<br>LAR2 3.98 0.3830 455 3.23 4.73<br>tissue = Seedling:<br>Genes emmean SE df lower.CL upper.CL<br>LAR1 3.87 0.4200 455 3.04 4.70<br>LAR2 3.15 0.4200 455 2.32 3.97<br>tissue = Young red leaves:<br>Genes emmean SE df lower.CL upper.CL<br>LAR1 3.47 0.3680 455 2.74 4.19<br>LAR2 2.39 0.3680 455 1.67 3.11 | tissue = Cells:<br>contrast estimate SE df t.ratio p.value<br>LAR1 - LAR2 0.996 0.1640 350 6.091 <.0001<br>tissue = Inflorescence:<br>contrast estimate SE df t.ratio p.value<br>LAR1 - LAR2 0.744 0.3060 350 2.429 0.0156<br>tissue = Leaf:<br>contrast estimate SE df t.ratio p.value<br>LAR1 - LAR2 0.426 0.0615 350 6.931 <.0001<br>tissue = Mix-leaf-flower:<br>contrast estimate SE df t.ratio p.value<br>LAR1 - LAR2 0.617 0.2160 350 2.849 0.0046<br>tissue = Pistil:<br>contrast estimate SE df t.ratio p.value<br>LAR1 - LAR2 -1.283 0.3060 350 -4.192 <.0001<br>tissue = Pod:<br>contrast estimate SE df t.ratio p.value<br>LAR1 - LAR2 -2.706 0.2790 350 -9.684 <.0001<br>tissue = Seedling:<br>contrast estimate SE df t.ratio p.value<br>LAR1 - LAR2 0.722 0.3060 350 2.357 0.0190<br>tissue = Young red leaves:<br>contrast estimate SE df t.ratio p.value<br>LAR1 - LAR2 1.077 0.2680 350 4.012 0.0001<br>Degrees-of-freedom method: kenward-roger<br>Confidence level used: 0.95 |

### *Moringa oleifera*

Linear mixed model fit by REML. t-tests use Satterthwaite's method ['lmerModLmerTest']
Formula: TPM_log ~ Genes * tissue + (1 | Sample)
Data: data_long

REML criterion at convergence: 221.1

Scaled residuals:

| Min | 1Q | Median | 3Q | Max |
|---|---|---|---|---|
| -1.90128 | -0.47035 | -0.05556 | 0.55924 | 2.55943 |

Random effects:

| Groups | Name | Variance | Std.Dev. |
|---|---|---|---|
| Sample | (Intercept) | 0.1212 | 0.3482 |
| Residual | 0.6087 | 0.7802 | |
| Number of obs: | 92 | groups: | Sample, 46 |

Fixed effects:

| | *Estimate* | *Std. Error* | *df* | *T-value* | *Pr(>\|t\|)* | |
|---|---|---|---|---|---|---|
| *(Intercept)* | *2.48954* | *0.3488* | *77.85264* | *7.137* | *4.35E-10* | *** |
| *GenesLAR2* | *0.18026* | *0.45046* | *40* | *0.4* | *0.69116* | |
| *tissueFlower* | *-2.03704* | *0.60414* | *77.85264* | *-3.372* | *0.00117* | ** |
| *tissueLeaf* | *-2.03072* | *0.39166* | *77.85264* | *-5.185* | *1.66E-06* | *** |
| *tissuePod* | *-2.31756* | *0.6976* | *77.85264* | *-3.322* | *0.00136* | ** |
| *tissueRoot* | *0.71184* | *0.46142* | *77.85264* | *1.543* | *0.12696* | |
| *tissueStem* | *0.3158* | *0.5515* | *77.85264* | *0.573* | *0.56855* | |
| *GenesLAR2:tissueFlower* | *0.57236* | *0.78022* | *40* | *0.734* | *0.46748* | |
| *GenesLAR2:tissueLeaf* | *1.10037* | *0.50581* | *40* | *2.175* | *0.03556* | * |
| *GenesLAR2:tissuePod* | *0.02109* | *0.90092* | *40* | *0.023* | *0.98144* | |
| *GenesLAR2:tissueRoot* | *1.10062* | *0.5959* | *40* | *1.847* | *0.07216* | . |
| *GenesLAR2:tissueStem* | *1.47121* | *0.71224* | *40* | *2.066* | *0.04538* | * |

Signif. codes: 0 '***' 0.001 '**' 0.01 '*' 0.05 '.' 0.1 ' ' 1

**Table 8S.** Estimated marginal means and pairwise comparison for the species *Moringa oleifera*. Degrees-of-freedom method: kenward-roger. Confidence level used: 0.95.

| ***Estimated Marginal Means*** | ***Contrasts*** |
|---|---|
| tissue = Callus:<br>Genes emmean SE df lower.CL upper.CL<br>LAR1 2.490 0.349 77.8 1.795 3.184<br>LAR2 2.670 0.349 77.8 1.975 3.364<br>tissue = Flower:<br>Genes emmean SE df lower.CL upper.CL<br>LAR1 0.453 0.493 77.8 -0.530 1.435 | tissue = Callus:<br>contrast estimate SE df t.ratio p.value<br>LAR1 - LAR2 -0.180 0.450 40 -0.400 0.6912<br>tissue = Flower:<br>contrast estimate SE df t.ratio p.value<br>LAR1 - LAR2 -0.753 0.637 40 -1.181 0.2444<br>tissue = Leaf: |

| | |
|---|---|
| LAR2 1.205 0.493 77.8 0.223 2.187<br>tissue = Leaf:<br>Genes emmean SE df lower.CL upper.CL<br>LAR1 0.459 0.178 77.8 0.104 0.813<br>LAR2 1.739 0.178 77.8 1.385 2.094<br>tissue = Pod:<br>Genes emmean SE df lower.CL upper.CL<br>LAR1 0.172 0.604 77.8 -1.031 1.375<br>LAR2 0.373 0.604 77.8 -0.829 1.576<br>tissue = Root:<br>Genes emmean SE df lower.CL upper.CL<br>LAR1 3.201 0.302 77.8 2.600 3.803<br>LAR2 4.482 0.302 77.8 3.881 5.084<br>tissue = Stem:<br>Genes emmean SE df lower.CL upper.CL<br>LAR1 2.805 0.427 77.8 1.955 3.656<br>LAR2 4.457 0.427 77.8 3.606 5.307 | contrast estimate SE df t.ratio p.value<br>LAR1 - LAR2 -1.281 0.230 40 -5.566 <.0001<br>tissue = Pod:<br>contrast estimate SE df t.ratio p.value<br>LAR1 - LAR2 -0.201 0.780 40 -0.258 0.7977<br>tissue = Root:<br>contrast estimate SE df t.ratio p.value<br>LAR1 - LAR2 -1.281 0.390 40 -3.283 0.0021<br>tissue = Stem:<br>contrast estimate SE df t.ratio p.value<br>LAR1 - LAR2 -1.651 0.552 40 -2.993 0.0047<br><br>Degrees-of-freedom method: kenward-roger<br>Confidence level used: 0.95 |

### *Gossypium hirsutum*

Linear mixed model fit by REML. t-tests use Satterthwaite's method ['lmerModLmerTest']
Formula: TPM_log ~ Genes * tissue + (1 | Sample)
Data: data_long

REML criterion at convergence: 10769.4

Scaled residuals:

| Min | 1Q | Median | 3Q | Max |
|---|---|---|---|---|
| -3.7343 | -0.5257 | -0.0146 | 0.4647 | 4.155 |

Random effects:

| Groups | Name | Variance | Std.Dev. |
|---|---|---|---|
| Sample | (Intercept) | 0.8483 | 0.921 |
| Residual | 0.4139 | 0.6433 | |
| Number of obs: | 4316 | groups: | Sample, 1079 |

Fixed effects:

| | *Estimate* | *Std. Error* | *df* | *T-value* | *Pr(>\|t\|)* | |
|---|---|---|---|---|---|---|
| *(Intercept)* | 2.47E+00 | 1.26E-01 | 1.78E+03 | 19.695 | <2E-016 | *** |
| *GenesLAR1b* | 5.01E-01 | 1.02E-01 | 3.14E+03 | 4.924 | 8.92E-07 | *** |
| *GenesLAR2a* | -1.14E+00 | 1.02E-01 | 3.14E+03 | -11.196 | <2E-016 | *** |
| *GenesLAR2b* | -5.02E-02 | 1.02E-01 | 3.14E+03 | -0.493 | 0.622041 | |

| | | | | | | |
|---|---|---|---|---|---|---|
| *tissueBinucleate* | -5.22E-01 | 4.76E-01 | 1.78E+03 | -1.097 | 0.272634 | |
| *tissueBoll* | 1.06E-01 | 8.04E-01 | 1.78E+03 | 0.132 | 0.894907 | |
| *tissueBoll-Shell* | 1.94E+00 | 1.13E+00 | 1.78E+03 | 1.717 | 0.0862 | . |
| *tissueBracts* | 2.27E+00 | 1.13E+00 | 1.78E+03 | 2.01 | 0.044592 | * |
| *tissueBud* | 1.27E+00 | 2.93E-01 | 1.78E+03 | 4.325 | 1.61E-05 | *** |
| *tissueCallus* | -4.69E-01 | 3.48E-01 | 1.78E+03 | -1.349 | 0.177454 | |
| *tissueCalycle* | 2.82E+00 | 1.13E+00 | 1.78E+03 | 2.491 | 0.012836 | * |
| *tissueCalyxs* | 6.93E-01 | 1.13E+00 | 1.78E+03 | 0.613 | 0.539894 | |
| *tissueCell* | -1.34E+00 | 4.76E-01 | 1.78E+03 | -2.825 | 0.004784 | ** |
| *tissueCorollas* | -1.98E+00 | 1.13E+00 | 1.78E+03 | -1.751 | 0.080116 | . |
| *tissueCotyledon* | 1.64E+00 | 2.09E-01 | 1.78E+03 | 7.846 | 7.36E-15 | *** |
| *tissueEpidermal cells* | 1.66E-01 | 5.18E-01 | 1.78E+03 | 0.32 | 0.748766 | |
| *tissueFiber* | -4.90E-01 | 1.51E-01 | 1.78E+03 | -3.246 | 0.001193 | ** |
| *tissueFlower* | -7.94E-01 | 6.61E-01 | 1.78E+03 | -1.202 | 0.229578 | |
| *tissueGynoecium* | 6.15E-01 | 8.04E-01 | 1.78E+03 | 0.765 | 0.444663 | |
| *tissueHypocotyl* | 5.00E-01 | 3.36E-01 | 1.78E+03 | 1.489 | 0.136794 | |
| *tissueJunction* | 2.22E+00 | 5.18E-01 | 1.78E+03 | 4.279 | 1.97E-05 | *** |
| *tissueLeaf* | 2.01E+00 | 1.47E-01 | 1.78E+03 | 13.692 | <2E-016 | *** |
| *tissueMature seed* | -2.47E+00 | 8.04E-01 | 1.78E+03 | -3.076 | 0.002131 | ** |
| *tissueMeristem* | 1.82E+00 | 8.04E-01 | 1.78E+03 | 2.259 | 0.02403 | * |
| *tissueOvule* | 7.61E-01 | 1.58E-01 | 1.78E+03 | 4.823 | 1.53E-06 | *** |
| *tissuePetal* | -2.34E-01 | 3.36E-01 | 1.78E+03 | -0.695 | 0.487099 | |
| *tissuePollen* | -2.47E+00 | 4.76E-01 | 1.78E+03 | -5.202 | 2.20E-07 | *** |
| *tissueProtoplasts* | -2.02E+00 | 4.76E-01 | 1.78E+03 | -4.244 | 2.31E-05 | *** |
| *tissueRoots* | 5.12E-01 | 1.49E-01 | 1.78E+03 | 3.441 | 0.000592 | *** |
| *tissueSeed* | -8.33E-01 | 3.36E-01 | 1.78E+03 | -2.48 | 0.013241 | * |
| *tissueSeedlings* | -5.21E-02 | 4.76E-01 | 1.78E+03 | -0.11 | 0.912698 | |
| *tissueShoot Apical* | 1.93E+00 | 2.54E-01 | 1.78E+03 | 7.603 | 4.66E-14 | *** |
| *tissueStamen* | -2.56E-01 | 1.13E+00 | 1.78E+03 | -0.226 | 0.82108 | |
| *tissueStem* | 1.53E+00 | 1.95E-01 | 1.78E+03 | 7.835 | 8.01E-15 | *** |
| *tissueTetrads* | -9.47E-03 | 4.76E-01 | 1.78E+03 | -0.02 | 0.984118 | |
| *tissueTorus* | 3.71E+00 | 1.13E+00 | 1.78E+03 | 3.285 | 0.001039 | ** |
| *tissueUninucleate* | 3.42E-01 | 4.76E-01 | 1.78E+03 | 0.719 | 0.472033 | |
| *GenesLAR1b:tissueBinucleate* | -1.04E+00 | 3.85E-01 | 3.14E+03 | -2.695 | 0.007079 | ** |
| *GenesLAR2a:tissueBinucleate* | -6.32E-01 | 3.85E-01 | 3.14E+03 | -1.641 | 0.100989 | |
| *GenesLAR2b:tissueBinucleate* | -1.25E+00 | 3.85E-01 | 3.14E+03 | -3.246 | 0.001183 | ** |
| *GenesLAR1b:tissueBoll* | 7.23E-01 | 6.51E-01 | 3.14E+03 | 1.11 | 0.267139 | |
| *GenesLAR2a:tissueBoll* | 6.87E-01 | 6.51E-01 | 3.14E+03 | 1.055 | 0.291578 | |
| *GenesLAR2b:tissueBoll* | 1.93E+00 | 6.51E-01 | 3.14E+03 | 2.959 | 0.003114 | ** |
| *GenesLAR1b:tissueBoll-Shell* | 1.33E-01 | 9.16E-01 | 3.14E+03 | 0.145 | 0.884668 | |
| *GenesLAR2a:tissueBoll-Shell* | -2.67E-02 | 9.16E-01 | 3.14E+03 | -0.029 | 0.976767 | |
| *GenesLAR2b:tissueBoll-Shell* | 5.83E-03 | 9.16E-01 | 3.14E+03 | 0.006 | 0.99492 | |
| *GenesLAR1b:tissueBracts* | -2.22E+00 | 9.16E-01 | 3.14E+03 | -2.429 | 0.015199 | * |
| *GenesLAR2a:tissueBracts* | -2.54E+00 | 9.16E-01 | 3.14E+03 | -2.776 | 0.005531 | ** |
| *GenesLAR2b:tissueBracts* | 7.59E-01 | 9.16E-01 | 3.14E+03 | 0.829 | 0.407374 | |

| | | | | | | |
|---|---|---|---|---|---|---|
| *GenesLAR1b:tissueBud* | 6.53E-02 | 2.37E-01 | 3.14E+03 | 0.275 | 0.783215 | |
| *GenesLAR2a:tissueBud* | -1.15E-02 | 2.37E-01 | 3.14E+03 | -0.049 | 0.96123 | |
| *GenesLAR2b:tissueBud* | -1.37E-01 | 2.37E-01 | 3.14E+03 | -0.576 | 0.56447 | |
| *GenesLAR1b:tissueCallus* | 2.86E-01 | 2.82E-01 | 3.14E+03 | 1.014 | 0.310522 | |
| *GenesLAR2a:tissueCallus* | 6.34E-01 | 2.82E-01 | 3.14E+03 | 2.253 | 0.02435 | * |
| *GenesLAR2b:tissueCallus* | 1.41E-01 | 2.82E-01 | 3.14E+03 | 0.501 | 0.616634 | |
| *GenesLAR1b:tissueCalycle* | -1.82E+00 | 9.16E-01 | 3.14E+03 | -1.991 | 0.046587 | * |
| *GenesLAR2a:tissueCalycle* | -1.25E+00 | 9.16E-01 | 3.14E+03 | -1.367 | 0.171628 | |
| *GenesLAR2b:tissueCalycle* | 1.30E+00 | 9.16E-01 | 3.14E+03 | 1.42 | 0.155711 | |
| *GenesLAR1b:tissueCalyxs* | -1.31E+00 | 9.16E-01 | 3.14E+03 | -1.43 | 0.152906 | |
| *GenesLAR2a:tissueCalyxs* | -1.59E+00 | 9.16E-01 | 3.14E+03 | -1.733 | 0.083138 | . |
| *GenesLAR2b:tissueCalyxs* | 4.93E-01 | 9.16E-01 | 3.14E+03 | 0.539 | 0.590062 | |
| *GenesLAR1b:tissueCell* | 3.62E-01 | 3.85E-01 | 3.14E+03 | 0.941 | 0.346686 | |
| *GenesLAR2a:tissueCell* | 5.99E-01 | 3.85E-01 | 3.14E+03 | 1.556 | 0.119817 | |
| *GenesLAR2b:tissueCell* | -9.67E-02 | 3.85E-01 | 3.14E+03 | -0.251 | 0.801669 | |
| *GenesLAR1b:tissueCorollas* | -6.59E-01 | 9.16E-01 | 3.14E+03 | -0.72 | 0.471676 | |
| *GenesLAR2a:tissueCorollas* | 6.78E-01 | 9.16E-01 | 3.14E+03 | 0.741 | 0.458873 | |
| *GenesLAR2b:tissueCorollas* | 8.56E-01 | 9.16E-01 | 3.14E+03 | 0.935 | 0.34961 | |
| *GenesLAR1b:tissueCotyledon* | -6.54E-01 | 1.70E-01 | 3.14E+03 | -3.858 | 0.000117 | *** |
| *GenesLAR2a:tissueCotyledon* | -2.31E+00 | 1.70E-01 | 3.14E+03 | -13.629 | <2E-016 | *** |
| *GenesLAR2b:tissueCotyledon* | -1.52E+00 | 1.70E-01 | 3.14E+03 | -8.945 | <2E-016 | *** |
| *GenesLAR1b:tissueEpidermal Cells* | 6.13E-01 | 4.19E-01 | 3.14E+03 | 1.462 | 0.143956 | |
| *GenesLAR2a:tissueEpidermal Cells* | 4.13E-01 | 4.19E-01 | 3.14E+03 | 0.985 | 0.324945 | |
| *GenesLAR2b:tissueEpidermal Cells* | 1.87E+00 | 4.19E-01 | 3.14E+03 | 4.455 | 8.67E-06 | *** |
| *GenesLAR1b:tissueFiber* | 5.86E-01 | 1.22E-01 | 3.14E+03 | 4.794 | 1.71E-06 | *** |
| *GenesLAR2a:tissueFiber* | 7.86E-01 | 1.22E-01 | 3.14E+03 | 6.431 | 1.46E-10 | *** |
| *GenesLAR2b:tissueFiber* | 1.51E+00 | 1.22E-01 | 3.14E+03 | 12.387 | <2E-016 | *** |
| *GenesLAR1b:tissueFlower* | -4.75E-01 | 5.35E-01 | 3.14E+03 | -0.888 | 0.374655 | |
| *GenesLAR2a:tissueFlower* | 4.51E-01 | 5.35E-01 | 3.14E+03 | 0.842 | 0.399598 | |
| *GenesLAR2b:tissueFlower* | 1.25E+00 | 5.35E-01 | 3.14E+03 | 2.343 | 0.019213 | * |
| *GenesLAR1b:tissueGynoecium* | -3.98E-01 | 6.51E-01 | 3.14E+03 | -0.611 | 0.541297 | |
| *GenesLAR2a:tissueGynoecium* | -6.34E-01 | 6.51E-01 | 3.14E+03 | -0.973 | 0.330457 | |
| *GenesLAR2b:tissueGynoecium* | 1.11E+00 | 6.51E-01 | 3.14E+03 | 1.704 | 0.088439 | . |
| *GenesLAR1b:tissueHypocotyl* | 2.60E-01 | 2.72E-01 | 3.14E+03 | 0.957 | 0.338762 | |
| *GenesLAR2a:tissueHypocotyl* | 1.53E+00 | 2.72E-01 | 3.14E+03 | 5.612 | 2.17E-08 | *** |
| *GenesLAR2b:tissueHypocotyl* | 5.38E-01 | 2.72E-01 | 3.14E+03 | 1.979 | 0.047912 | * |
| *GenesLAR1b:tissueJunction* | -2.95E-01 | 4.19E-01 | 3.14E+03 | -0.704 | 0.481244 | |
| *GenesLAR2a:tissueJunction* | -1.48E+00 | 4.19E-01 | 3.14E+03 | -3.517 | 0.000443 | *** |
| *GenesLAR2b:tissueJunction* | -5.70E-01 | 4.19E-01 | 3.14E+03 | -1.359 | 0.174102 | |
| *GenesLAR1b:tissueLeaf* | -3.63E-01 | 1.19E-01 | 3.14E+03 | -3.045 | 0.002348 | ** |
| *GenesLAR2a:tissueLeaf* | -2.15E+00 | 1.19E-01 | 3.14E+03 | -18.021 | <2E-016 | *** |
| *GenesLAR2b:tissueLeaf* | -1.04E+00 | 1.19E-01 | 3.14E+03 | -8.724 | <2E-016 | *** |
| *GenesLAR1b:tissueMature Seed* | -5.01E-01 | 6.51E-01 | 3.14E+03 | -0.769 | 0.44195 | |
| *GenesLAR2a:tissueMature Seed* | 1.21E+00 | 6.51E-01 | 3.14E+03 | 1.865 | 0.062339 | |
| *GenesLAR2b:tissueMature Seed* | 5.02E-02 | 6.51E-01 | 3.14E+03 | 0.077 | 0.938633 | |

| | | | | | | |
|---|---|---|---|---|---|---|
| *GenesLAR1b:tissueMeristem* | -2.00E-01 | 6.51E-01 | 3.14E+03 | -0.308 | 0.758303 | |
| *GenesLAR2a:tissueMeristem* | -1.19E+00 | 6.51E-01 | 3.14E+03 | -1.83 | 0.067272 | . |
| *GenesLAR2b:tissueMeristem* | 5.35E-01 | 6.51E-01 | 3.14E+03 | 0.822 | 0.411118 | |
| *GenesLAR1b:tissueOvule* | 4.61E-01 | 1.28E-01 | 3.14E+03 | 3.602 | 0.00032 | *** |
| *GenesLAR2a:tissueOvule* | 2.90E-01 | 1.28E-01 | 3.14E+03 | 2.266 | 0.023514 | * |
| *GenesLAR2b:tissueOvule* | 2.73E-01 | 1.28E-01 | 3.14E+03 | 2.138 | 0.032556 | * |
| *GenesLAR1b:tissuePetal* | 8.95E-01 | 2.72E-01 | 3.14E+03 | 3.288 | 0.001018 | ** |
| *GenesLAR2a:tissuePetal* | -1.02E+00 | 2.72E-01 | 3.14E+03 | -3.731 | 0.000194 | *** |
| *GenesLAR2b:tissuePetal* | -4.60E-01 | 2.72E-01 | 3.14E+03 | -1.69 | 0.0912 | . |
| *GenesLAR1b:tissuePollen* | -5.01E-01 | 3.85E-01 | 3.14E+03 | -1.301 | 0.193489 | |
| *GenesLAR2a:tissuePollen* | 1.15E+00 | 3.85E-01 | 3.14E+03 | 2.989 | 0.002825 | ** |
| *GenesLAR2b:tissuePollen* | 5.02E-02 | 3.85E-01 | 3.14E+03 | 0.13 | 0.8964 | |
| *GenesLAR1b:tissueProtoplasts* | 3.20E-02 | 3.85E-01 | 3.14E+03 | 0.083 | 0.933774 | |
| *GenesLAR2a:tissueProtoplasts* | 6.86E-01 | 3.85E-01 | 3.14E+03 | 1.782 | 0.074806 | . |
| *GenesLAR2b:tissueProtoplasts* | 5.03E-01 | 3.85E-01 | 3.14E+03 | 1.307 | 0.1912 | |
| *GenesLAR1b:tissueRoots* | 2.42E-01 | 1.21E-01 | 3.14E+03 | 2.006 | 0.044918 | * |
| *GenesLAR2a:tissueRoots* | -2.85E-01 | 1.21E-01 | 3.14E+03 | -2.366 | 0.01806 | * |
| *GenesLAR2b:tissueRoots* | -6.33E-01 | 1.21E-01 | 3.14E+03 | -5.248 | 1.64E-07 | *** |
| *GenesLAR1b:tissueSeed* | 3.71E-02 | 2.72E-01 | 3.14E+03 | 0.136 | 0.891652 | |
| *GenesLAR2a:tissueSeed* | 4.35E-01 | 2.72E-01 | 3.14E+03 | 1.597 | 0.110266 | |
| *GenesLAR2b:tissueSeed* | -4.78E-01 | 2.72E-01 | 3.14E+03 | -1.756 | 0.079215 | . |
| *GenesLAR1b:tissueSeedlings* | -1.39E-01 | 3.85E-01 | 3.14E+03 | -0.362 | 0.717453 | |
| *GenesLAR2a:tissueSeedlings* | -1.09E+00 | 3.85E-01 | 3.14E+03 | -2.833 | 0.004636 | ** |
| *GenesLAR2b:tissueSeedlings* | -1.89E+00 | 3.85E-01 | 3.14E+03 | -4.916 | 9.31E-07 | *** |
| *GenesLAR1b:tissueShoot Apical* | 1.48E-01 | 2.05E-01 | 3.14E+03 | 0.72 | 0.471878 | |
| *GenesLAR2a:tissueShoot Apical* | -8.26E-01 | 2.05E-01 | 3.14E+03 | -4.019 | 5.97E-05 | *** |
| *GenesLAR2b:tissueShoot Apical* | -3.14E-01 | 2.05E-01 | 3.14E+03 | -1.529 | 0.126367 | |
| *GenesLAR1b:tissueStamen* | -6.84E-02 | 9.16E-01 | 3.14E+03 | -0.075 | 0.940462 | |
| *GenesLAR2a:tissueStamen* | -8.93E-01 | 9.16E-01 | 3.14E+03 | -0.975 | 0.329537 | |
| *GenesLAR2b:tissueStamen* | -1.75E+00 | 9.16E-01 | 3.14E+03 | -1.916 | 0.055438 | . |
| *GenesLAR1b:tissueStem* | -7.41E-02 | 1.58E-01 | 3.14E+03 | -0.47 | 0.638608 | |
| *GenesLAR2a:tissueStem* | -1.26E+00 | 1.58E-01 | 3.14E+03 | -7.964 | 2.31E-15 | *** |
| *GenesLAR2b:tissueStem* | -5.18E-01 | 1.58E-01 | 3.14E+03 | -3.283 | 0.001038 | ** |
| *GenesLAR1b:tissueTetrads* | 1.56E-01 | 3.85E-01 | 3.14E+03 | 0.405 | 0.68521 | |
| *GenesLAR2a:tissueTetrads* | -2.04E-01 | 3.85E-01 | 3.14E+03 | -0.531 | 0.59561 | |
| *GenesLAR2b:tissueTetrads* | -7.22E-01 | 3.85E-01 | 3.14E+03 | -1.875 | 0.060919 | . |
| *GenesLAR1b:tissueTorus* | -3.43E-01 | 9.16E-01 | 3.14E+03 | -0.374 | 0.708182 | |
| *GenesLAR2a:tissueTorus* | -1.70E+00 | 9.16E-01 | 3.14E+03 | -1.852 | 0.064133 | . |
| *GenesLAR2b:tissueTorus* | -1.65E+00 | 9.16E-01 | 3.14E+03 | -1.806 | 0.071054 | . |
| *GenesLAR1b:tissueUninucleate* | 1.73E-01 | 3.85E-01 | 3.14E+03 | 0.45 | 0.65248 | |
| *GenesLAR2a:tissueUninucleate* | -1.32E-01 | 3.85E-01 | 3.14E+03 | -0.344 | 0.731145 | |
| *GenesLAR2b:tissueUninucleate* | -5.43E-01 | 3.85E-01 | 3.14E+03 | -1.411 | 0.158378 | |

**Table 9S.** Estimated marginal means and pairwise comparison for the species *Gossypium hirsutum*. Degrees-of-freedom method: asymptotic. P value adjustment: bonferroni method for 6 tests

| ***Estimated Marginal Means*** | ***Contrasts*** |
|---|---|
| tissue = Anther: | tissue = Anther: |
| Genes emmean SE df asymp.LCL asymp.UCL | contrast estimate SE df z.ratio p.value |
| LAR1a 2.47375 0.1260 Inf 2.2276 2.720 | LAR1a - LAR1b -0.5009 0.1020 Inf -4.924 <.0001 |
| LAR1b 2.97461 0.1260 Inf 2.7284 3.221 | LAR1a - LAR2a 1.1388 0.1020 Inf 11.196 <.0001 |
| LAR2a 1.33492 0.1260 Inf 1.0887 1.581 | LAR1a - LAR2b 0.0501 0.1020 Inf 0.493 1.0000 |
| LAR2b 2.42360 0.1260 Inf 2.1774 2.670 | LAR1b - LAR2a 1.6397 0.1020 Inf 16.120 <.0001 |
| | LAR1b - LAR2b 0.5510 0.1020 Inf 5.417 <.0001 |
| tissue = Binucleate: | LAR2a - LAR2b -1.0887 0.1020 Inf -10.703 <.0001 |
| Genes emmean SE df asymp.LCL asymp.UCL | |
| LAR1a 1.95192 0.4590 Inf 1.0530 2.851 | tissue = Binucleate: |
| LAR1b 1.41500 0.4590 Inf 0.5161 2.314 | contrast estimate SE df z.ratio p.value |
| LAR2a 0.18132 0.4590 Inf -0.7176 1.080 | LAR1a - LAR1b 0.5369 0.3710 Inf 1.446 0.8897 |
| LAR2b 0.65175 0.4590 Inf -0.2472 1.551 | LAR1a - LAR2a 1.7706 0.3710 Inf 4.767 <.0001 |
| | LAR1a - LAR2b 1.3002 0.3710 Inf 3.501 0.0028 |
| tissue = Boll: | LAR1b - LAR2a 1.2337 0.3710 Inf 3.322 0.0054 |
| Genes emmean SE df asymp.LCL asymp.UCL | LAR1b - LAR2b 0.7632 0.3710 Inf 2.055 0.2393 |
| LAR1a 2.58000 0.7940 Inf 1.0230 4.137 | LAR2a - LAR2b -0.4704 0.3710 Inf -1.267 1.0000 |
| LAR1b 3.80374 0.7940 Inf 2.2468 5.361 | |
| LAR2a 2.12821 0.7940 Inf 0.5712 3.685 | tissue = Boll: |
| LAR2b 4.45679 0.7940 Inf 2.8998 6.014 | contrast estimate SE df z.ratio p.value |
| | LAR1a - LAR1b -1.2237 0.6430 Inf -1.902 0.3429 |
| tissue = Boll-Shell: | LAR1a - LAR2a 0.4518 0.6430 Inf 0.702 1.0000 |
| Genes emmean SE df asymp.LCL asymp.UCL | LAR1a - LAR2b -1.8768 0.6430 Inf -2.917 0.0212 |
| LAR1a 4.41443 1.1200 Inf 2.2125 6.616 | LAR1b - LAR2a 1.6755 0.6430 Inf 2.604 0.0552 |
| LAR1b 5.04809 1.1200 Inf 2.8462 7.250 | LAR1b - LAR2b -0.6530 0.6430 Inf -1.015 1.0000 |
| LAR2a 3.24894 1.1200 Inf 1.0470 5.451 | LAR2a - LAR2b -2.3286 0.6430 Inf -3.620 0.0018 |
| LAR2b 4.37011 1.1200 Inf 2.1682 6.572 | |
| | tissue = Boll-Shell: |
| tissue = Bracts: | contrast estimate SE df z.ratio p.value |
| Genes emmean SE df asymp.LCL asymp.UCL | LAR1a - LAR1b -0.6337 0.9100 Inf -0.696 1.0000 |
| LAR1a 4.74583 1.1200 Inf 2.5439 6.948 | LAR1a - LAR2a 1.1655 0.9100 Inf 1.281 1.0000 |
| LAR1b 3.02310 1.1200 Inf 0.8212 5.225 | LAR1a - LAR2b 0.0443 0.9100 Inf 0.049 1.0000 |
| LAR2a 1.06540 1.1200 Inf -1.1365 3.267 | LAR1b - LAR2a 1.7991 0.9100 Inf 1.978 0.2879 |
| LAR2b 5.45426 1.1200 Inf 3.2524 7.656 | LAR1b - LAR2b 0.6780 0.9100 Inf 0.745 1.0000 |
| | LAR2a - LAR2b -1.1212 0.9100 Inf -1.232 1.0000 |
| tissue = Bud: | |
| Genes emmean SE df asymp.LCL asymp.UCL | tissue = Bracts: |
| LAR1a 3.74127 0.2650 Inf 3.2223 4.260 | contrast estimate SE df z.ratio p.value |
| LAR1b 4.30744 0.2650 Inf 3.7884 4.826 | LAR1a - LAR1b 1.7227 0.9100 Inf 1.894 0.3497 |
| LAR2a 2.59091 0.2650 Inf 2.0719 3.110 | LAR1a - LAR2a 3.6804 0.9100 Inf 4.045 0.0003 |
| LAR2b 3.55435 0.2650 Inf 3.0354 4.073 | LAR1a - LAR2b -0.7084 0.9100 Inf -0.779 1.0000 |
| | LAR1b - LAR2a 1.9577 0.9100 Inf 2.152 0.1885 |
| tissue = Callus: | LAR1b - LAR2b -2.4312 0.9100 Inf -2.672 0.0452 |
| Genes emmean SE df asymp.LCL asymp.UCL | LAR2a - LAR2b -4.3889 0.9100 Inf -4.824 <.0001 |
| LAR1a 2.00453 0.3240 Inf 1.3689 2.640 | |
| LAR1b 2.79106 0.3240 Inf 2.1554 3.427 | tissue = Bud: |

```
 LAR2a 1.50015 0.3240 Inf    0.8645     2.136
 LAR2b 2.09539 0.3240 Inf    1.4598     2.731

tissue = Calycle:
 Genes  emmean     SE  df asymp.LCL asymp.UCL
 LAR1a 5.28946 1.1200 Inf    3.0876     7.491
 LAR1b 3.96780 1.1200 Inf    1.7659     6.170
 LAR2a 2.89892 1.1200 Inf    0.6970     5.101
 LAR2b 6.53925 1.1200 Inf    4.3373     8.741

tissue = Calyxs:
 Genes  emmean     SE  df asymp.LCL asymp.UCL
 LAR1a 3.16681 1.1200 Inf    0.9649     5.369
 LAR1b 2.35885 1.1200 Inf    0.1569     4.561
 LAR2a 0.44121 1.1200 Inf   -1.7607     2.643
 LAR2b 3.60991 1.1200 Inf    1.4080     5.812

tissue = Cell:
 Genes  emmean     SE  df asymp.LCL asymp.UCL
 LAR1a 1.13046 0.4590 Inf    0.2315     2.029
 LAR1b 1.99376 0.4590 Inf    1.0948     2.893
 LAR2a 0.59083 0.4590 Inf   -0.3081     1.490
 LAR2b 0.98357 0.4590 Inf    0.0846     1.882

tissue = Corollas:
 Genes  emmean     SE  df asymp.LCL asymp.UCL
 LAR1a 0.49433 1.1200 Inf   -1.7076     2.696
 LAR1b 0.33621 1.1200 Inf   -1.8657     2.538
 LAR2a 0.03368 1.1200 Inf   -2.1682     2.236
 LAR2b 1.30058 1.1200 Inf   -0.9013     3.502

tissue = Cotyledon:
 Genes  emmean     SE  df asymp.LCL asymp.UCL
 LAR1a 4.11625 0.1670 Inf    3.7880     4.444
 LAR1b 3.96312 0.1670 Inf    3.6349     4.291
 LAR2a 0.66690 0.1670 Inf    0.3387     0.995
 LAR2b 2.54959 0.1670 Inf    2.2214     2.878

tissue = Epidermal Cells:
 Genes  emmean     SE  df asymp.LCL asymp.UCL
 LAR1a 2.63963 0.5020 Inf    1.6549     3.624
 LAR1b 3.75347 0.5020 Inf    2.7687     4.738
 LAR2a 1.91370 0.5020 Inf    0.9290     2.898
 LAR2b 4.45801 0.5020 Inf    3.4733     5.443

tissue = Fiber:
 Genes  emmean     SE  df asymp.LCL asymp.UCL
 LAR1a 1.98378 0.0837 Inf    1.8197     2.148
```

```
 contrast      estimate     SE  df z.ratio p.value
 LAR1a - LAR1b  -0.5662 0.2140 Inf  -2.640  0.0497
 LAR1a - LAR2a   1.1504 0.2140 Inf   5.364  <.0001
 LAR1a - LAR2b   0.1869 0.2140 Inf   0.872  1.0000
 LAR1b - LAR2a   1.7165 0.2140 Inf   8.005  <.0001
 LAR1b - LAR2b   0.7531 0.2140 Inf   3.512  0.0027
 LAR2a - LAR2b  -0.9634 0.2140 Inf  -4.493  <.0001

tissue = Callus:
 contrast      estimate     SE  df z.ratio p.value
 LAR1a - LAR1b  -0.7865 0.2630 Inf  -2.995  0.0165
 LAR1a - LAR2a   0.5044 0.2630 Inf   1.920  0.3288
 LAR1a - LAR2b  -0.0909 0.2630 Inf  -0.346  1.0000
 LAR1b - LAR2a   1.2909 0.2630 Inf   4.915  <.0001
 LAR1b - LAR2b   0.6957 0.2630 Inf   2.649  0.0485
 LAR2a - LAR2b  -0.5952 0.2630 Inf  -2.266  0.1405

tissue = Calycle:
 contrast      estimate     SE  df z.ratio p.value
 LAR1a - LAR1b   1.3217 0.9100 Inf   1.453  0.8778
 LAR1a - LAR2a   2.3905 0.9100 Inf   2.628  0.0516
 LAR1a - LAR2b  -1.2498 0.9100 Inf  -1.374  1.0000
 LAR1b - LAR2a   1.0689 0.9100 Inf   1.175  1.0000
 LAR1b - LAR2b  -2.5714 0.9100 Inf  -2.826  0.0282
 LAR2a - LAR2b  -3.6403 0.9100 Inf  -4.001  0.0004

tissue = Calyxs:
 contrast      estimate     SE  df z.ratio p.value
 LAR1a - LAR1b   0.8080 0.9100 Inf   0.888  1.0000
 LAR1a - LAR2a   2.7256 0.9100 Inf   2.996  0.0164
 LAR1a - LAR2b  -0.4431 0.9100 Inf  -0.487  1.0000
 LAR1b - LAR2a   1.9176 0.9100 Inf   2.108  0.2103
 LAR1b - LAR2b  -1.2511 0.9100 Inf  -1.375  1.0000
 LAR2a - LAR2b  -3.1687 0.9100 Inf  -3.483  0.0030

tissue = Cell:
 contrast      estimate     SE  df z.ratio p.value
 LAR1a - LAR1b  -0.8633 0.3710 Inf  -2.324  0.1206
 LAR1a - LAR2a   0.5396 0.3710 Inf   1.453  0.8776
 LAR1a - LAR2b   0.1469 0.3710 Inf   0.395  1.0000
 LAR1b - LAR2a   1.4029 0.3710 Inf   3.777  0.0010
 LAR1b - LAR2b   1.0102 0.3710 Inf   2.720  0.0392
 LAR2a - LAR2b  -0.3927 0.3710 Inf  -1.057  1.0000

tissue = Corollas:
 contrast      estimate     SE  df z.ratio p.value
 LAR1a - LAR1b   0.1581 0.9100 Inf   0.174  1.0000
 LAR1a - LAR2a   0.4607 0.9100 Inf   0.506  1.0000
```

```
 LAR1b 3.07076 0.0837 Inf    2.9066     3.235
 LAR2a 1.63119 0.0837 Inf    1.4671     1.795
 LAR2b 3.44798 0.0837 Inf    3.2839     3.612

tissue = Flower:
 Genes  emmean     SE  df asymp.LCL asymp.UCL
 LAR1a 1.67972 0.6490 Inf    0.4084     2.951
 LAR1b 1.70552 0.6490 Inf    0.4342     2.977
 LAR2a 0.99163 0.6490 Inf   -0.2796     2.263
 LAR2b 2.88291 0.6490 Inf    1.6116     4.154

tissue = Gynoecium:
 Genes  emmean     SE  df asymp.LCL asymp.UCL
 LAR1a 3.08862 0.7940 Inf    1.5316     4.646
 LAR1b 3.19158 0.7940 Inf    1.6346     4.749
 LAR2a 1.31584 0.7940 Inf   -0.2411     2.873
 LAR2b 4.14845 0.7940 Inf    2.5915     5.705

tissue = Hypocotyl:
 Genes  emmean     SE  df asymp.LCL asymp.UCL
 LAR1a 2.97381 0.3120 Inf    2.3631     3.585
 LAR1b 3.73497 0.3120 Inf    3.1243     4.346
 LAR2a 3.36181 0.3120 Inf    2.7511     3.973
 LAR2b 3.46205 0.3120 Inf    2.8514     4.073

tissue = Junction:
 Genes  emmean     SE  df asymp.LCL asymp.UCL
 LAR1a 4.68992 0.5020 Inf    3.7052     5.675
 LAR1b 4.89537 0.5020 Inf    3.9106     5.880
 LAR2a 2.07620 0.5020 Inf    1.0915     3.061
 LAR2b 4.06963 0.5020 Inf    3.0849     5.054

tissue = Leaf:
 Genes  emmean     SE  df asymp.LCL asymp.UCL
 LAR1a 4.48691 0.0764 Inf    4.3371     4.637
 LAR1b 4.62522 0.0764 Inf    4.4754     4.775
 LAR2a 1.20225 0.0764 Inf    1.0524     1.352
 LAR2b 3.39796 0.0764 Inf    3.2481     3.548

tissue = Mature Seed:
 Genes  emmean     SE  df asymp.LCL asymp.UCL
 LAR1a 0.00000 0.7940 Inf   -1.5570     1.557
 LAR1b 0.00000 0.7940 Inf   -1.5570     1.557
 LAR2a 0.07557 0.7940 Inf   -1.4814     1.633
 LAR2b 0.00000 0.7940 Inf   -1.5570     1.557

tissue = Meristem:
 Genes  emmean     SE  df asymp.LCL asymp.UCL
```

```
 LAR1a - LAR2b  -0.8063 0.9100 Inf  -0.886  1.0000
 LAR1b - LAR2a   0.3025 0.9100 Inf   0.333  1.0000
 LAR1b - LAR2b  -0.9644 0.9100 Inf  -1.060  1.0000
 LAR2a - LAR2b  -1.2669 0.9100 Inf  -1.393  0.9826

tissue = Cotyledon:
 contrast      estimate     SE  df z.ratio p.value
 LAR1a - LAR1b   0.1531 0.1360 Inf   1.129  1.0000
 LAR1a - LAR2a   3.4494 0.1360 Inf  25.433  <.0001
 LAR1a - LAR2b   1.5667 0.1360 Inf  11.552  <.0001
 LAR1b - LAR2a   3.2962 0.1360 Inf  24.304  <.0001
 LAR1b - LAR2b   1.4135 0.1360 Inf  10.422  <.0001
 LAR2a - LAR2b  -1.8827 0.1360 Inf -13.882  <.0001

tissue = Epidermal Cells:
 contrast      estimate     SE  df z.ratio p.value
 LAR1a - LAR1b  -1.1138 0.4070 Inf  -2.738  0.0371
 LAR1a - LAR2a   0.7259 0.4070 Inf   1.784  0.4464
 LAR1a - LAR2b  -1.8184 0.4070 Inf  -4.469  <.0001
 LAR1b - LAR2a   1.8398 0.4070 Inf   4.522  <.0001
 LAR1b - LAR2b  -0.7045 0.4070 Inf  -1.732  0.5001
 LAR2a - LAR2b  -2.5443 0.4070 Inf  -6.253  <.0001

tissue = Fiber:
 contrast      estimate     SE  df z.ratio p.value
 LAR1a - LAR1b  -1.0870 0.0678 Inf -16.029  <.0001
 LAR1a - LAR2a   0.3526 0.0678 Inf   5.200  <.0001
 LAR1a - LAR2b  -1.4642 0.0678 Inf -21.592  <.0001
 LAR1b - LAR2a   1.4396 0.0678 Inf  21.229  <.0001
 LAR1b - LAR2b  -0.3772 0.0678 Inf  -5.563  <.0001
 LAR2a - LAR2b  -1.8168 0.0678 Inf -26.792  <.0001

tissue = Flower:
 contrast      estimate     SE  df z.ratio p.value
 LAR1a - LAR1b  -0.0258 0.5250 Inf  -0.049  1.0000
 LAR1a - LAR2a   0.6881 0.5250 Inf   1.310  1.0000
 LAR1a - LAR2b  -1.2032 0.5250 Inf  -2.291  0.1319
 LAR1b - LAR2a   0.7139 0.5250 Inf   1.359  1.0000
 LAR1b - LAR2b  -1.1774 0.5250 Inf  -2.242  0.1500
 LAR2a - LAR2b  -1.8913 0.5250 Inf  -3.601  0.0019

tissue = Gynoecium:
 contrast      estimate     SE  df z.ratio p.value
 LAR1a - LAR1b  -0.1030 0.6430 Inf  -0.160  1.0000
 LAR1a - LAR2a   1.7728 0.6430 Inf   2.756  0.0351
 LAR1a - LAR2b  -1.0598 0.6430 Inf  -1.647  0.5968
 LAR1b - LAR2a   1.8757 0.6430 Inf   2.916  0.0213
 LAR1b - LAR2b  -0.9569 0.6430 Inf  -1.487  0.8215
```

```
 LAR1a 4.29025 0.7940 Inf    2.7333     5.847
 LAR1b 4.59068 0.7940 Inf    3.0337     6.148
 LAR2a 1.95921 0.7940 Inf    0.4022     3.516
 LAR2b 4.77550 0.7940 Inf    3.2185     6.332

tissue = Ovule:
 Genes  emmean     SE  df asymp.LCL asymp.UCL
 LAR1a 3.23517 0.0956 Inf    3.0477     3.423
 LAR1b 4.19656 0.0956 Inf    4.0091     4.384
 LAR2a 2.38605 0.0956 Inf    2.1986     2.573
 LAR2b 3.45841 0.0956 Inf    3.2710     3.646

tissue = Petal:
 Genes  emmean     SE  df asymp.LCL asymp.UCL
 LAR1a 2.24024 0.3120 Inf    1.6295     2.851
 LAR1b 3.63577 0.3120 Inf    3.0251     4.246
 LAR2a 0.08645 0.3120 Inf   -0.5242     0.697
 LAR2b 1.73041 0.3120 Inf    1.1197     2.341

tissue = Pollen:
 Genes  emmean     SE  df asymp.LCL asymp.UCL
 LAR1a 0.00000 0.4590 Inf   -0.8989     0.899
 LAR1b 0.00000 0.4590 Inf   -0.8989     0.899
 LAR2a 0.01204 0.4590 Inf   -0.8869     0.911
 LAR2b 0.00000 0.4590 Inf   -0.8989     0.899

tissue = Protoplasts:
 Genes  emmean     SE  df asymp.LCL asymp.UCL
 LAR1a 0.45580 0.4590 Inf   -0.4431     1.355
 LAR1b 0.98866 0.4590 Inf    0.0897     1.888
 LAR2a 0.00331 0.4590 Inf   -0.8956     0.902
 LAR2b 0.90909 0.4590 Inf    0.0102     1.808

tissue = Roots:
 Genes  emmean     SE  df asymp.LCL asymp.UCL
 LAR1a 2.98593 0.0798 Inf    2.8295     3.142
 LAR1b 3.72860 0.0798 Inf    3.5721     3.885
 LAR2a 1.56199 0.0798 Inf    1.4055     1.718
 LAR2b 2.30328 0.0798 Inf    2.1468     2.460

tissue = Seed:
 Genes  emmean     SE  df asymp.LCL asymp.UCL
 LAR1a 1.64069 0.3120 Inf    1.0300     2.251
 LAR1b 2.17861 0.3120 Inf    1.5679     2.789
 LAR2a 0.93647 0.3120 Inf    0.3258     1.547
 LAR2b 1.11285 0.3120 Inf    0.5021     1.724

tissue = Seedlings:
```

```
 LAR2a - LAR2b  -2.8326 0.6430 Inf  -4.403  0.0001

tissue = Hypocotyl:
 contrast      estimate     SE  df z.ratio p.value
 LAR1a - LAR1b  -0.7612 0.2520 Inf  -3.017  0.0153
 LAR1a - LAR2a  -0.3880 0.2520 Inf  -1.538  0.7448
 LAR1a - LAR2b  -0.4882 0.2520 Inf  -1.935  0.3180
 LAR1b - LAR2a   0.3732 0.2520 Inf   1.479  0.8351
 LAR1b - LAR2b   0.2729 0.2520 Inf   1.082  1.0000
 LAR2a - LAR2b  -0.1002 0.2520 Inf  -0.397  1.0000

tissue = Junction:
 contrast      estimate     SE  df z.ratio p.value
 LAR1a - LAR1b  -0.2054 0.4070 Inf  -0.505  1.0000
 LAR1a - LAR2a   2.6137 0.4070 Inf   6.424  <.0001
 LAR1a - LAR2b   0.6203 0.4070 Inf   1.525  0.7642
 LAR1b - LAR2a   2.8192 0.4070 Inf   6.929  <.0001
 LAR1b - LAR2b   0.8257 0.4070 Inf   2.029  0.2545
 LAR2a - LAR2b  -1.9934 0.4070 Inf  -4.899  <.0001

tissue = Leaf:
 contrast      estimate     SE  df z.ratio p.value
 LAR1a - LAR1b  -0.1383 0.0619 Inf  -2.234  0.1528
 LAR1a - LAR2a   3.2847 0.0619 Inf  53.061  <.0001
 LAR1a - LAR2b   1.0889 0.0619 Inf  17.591  <.0001
 LAR1b - LAR2a   3.4230 0.0619 Inf  55.295  <.0001
 LAR1b - LAR2b   1.2273 0.0619 Inf  19.825  <.0001
 LAR2a - LAR2b  -2.1957 0.0619 Inf -35.470  <.0001

tissue = Mature Seed:
 contrast      estimate     SE  df z.ratio p.value
 LAR1a - LAR1b   0.0000 0.6430 Inf   0.000  1.0000
 LAR1a - LAR2a  -0.0756 0.6430 Inf  -0.117  1.0000
 LAR1a - LAR2b   0.0000 0.6430 Inf   0.000  1.0000
 LAR1b - LAR2a  -0.0756 0.6430 Inf  -0.117  1.0000
 LAR1b - LAR2b   0.0000 0.6430 Inf   0.000  1.0000
 LAR2a - LAR2b   0.0756 0.6430 Inf   0.117  1.0000

tissue = Meristem:
 contrast      estimate     SE  df z.ratio p.value
 LAR1a - LAR1b  -0.3004 0.6430 Inf  -0.467  1.0000
 LAR1a - LAR2a   2.3310 0.6430 Inf   3.623  0.0017
 LAR1a - LAR2b  -0.4853 0.6430 Inf  -0.754  1.0000
 LAR1b - LAR2a   2.6315 0.6430 Inf   4.090  0.0003
 LAR1b - LAR2b  -0.1848 0.6430 Inf  -0.287  1.0000
 LAR2a - LAR2b  -2.8163 0.6430 Inf  -4.378  0.0001

tissue = Ovule:
```

```
 Genes  emmean     SE  df asymp.LCL asymp.UCL
 LAR1a 2.42161 0.4590 Inf    1.5227     3.321
 LAR1b 2.78310 0.4590 Inf    1.8842     3.682
 LAR2a 0.19167 0.4590 Inf   -0.7072     1.091
 LAR2b 0.47849 0.4590 Inf   -0.4204     1.377

tissue = Shoot Apical:
 Genes  emmean     SE  df asymp.LCL asymp.UCL
 LAR1a 4.40200 0.2200 Inf    3.9702     4.834
 LAR1b 5.05063 0.2200 Inf    4.6188     5.482
 LAR2a 2.43764 0.2200 Inf    2.0058     2.869
 LAR2b 4.03782 0.2200 Inf    3.6060     4.470

tissue = Stamen:
 Genes  emmean     SE  df asymp.LCL asymp.UCL
 LAR1a 2.21806 1.1200 Inf    0.0162     4.420
 LAR1b 2.65053 1.1200 Inf    0.4486     4.852
 LAR2a 0.18648 1.1200 Inf   -2.0154     2.388
 LAR2b 0.41376 1.1200 Inf   -1.7881     2.616

tissue = Stem:
 Genes  emmean     SE  df asymp.LCL asymp.UCL
 LAR1a 3.99944 0.1490 Inf    3.7078     4.291
 LAR1b 4.42623 0.1490 Inf    4.1346     4.718
 LAR2a 1.60470 0.1490 Inf    1.3131     1.896
 LAR2b 3.43154 0.1490 Inf    3.1399     3.723

tissue = Tetrads:
 Genes  emmean     SE  df asymp.LCL asymp.UCL
 LAR1a 2.46428 0.4590 Inf    1.5654     3.363
 LAR1b 3.12126 0.4590 Inf    2.2223     4.020
 LAR2a 1.12105 0.4590 Inf    0.2221     2.020
 LAR2b 1.69217 0.4590 Inf    0.7932     2.591

tissue = Torus:
 Genes  emmean     SE  df asymp.LCL asymp.UCL
 LAR1a 6.18732 1.1200 Inf    3.9854     8.389
 LAR1b 6.34549 1.1200 Inf    4.1436     8.547
 LAR2a 3.35315 1.1200 Inf    1.1512     5.555
 LAR2b 4.48408 1.1200 Inf    2.2822     6.686

tissue = Uninucleate:
 Genes  emmean     SE  df asymp.LCL asymp.UCL
 LAR1a 2.81581 0.4590 Inf    1.9169     3.715
 LAR1b 3.49010 0.4590 Inf    2.5912     4.389
 LAR2a 1.54465 0.4590 Inf    0.6457     2.444
 LAR2b 2.22234 0.4590 Inf    1.3234     3.121
```

```
 contrast      estimate     SE  df z.ratio p.value
 LAR1a - LAR1b  -0.9614 0.0774 Inf -12.414  <.0001
 LAR1a - LAR2a   0.8491 0.0774 Inf  10.964  <.0001
 LAR1a - LAR2b  -0.2232 0.0774 Inf  -2.883  0.0237
 LAR1b - LAR2a   1.8105 0.0774 Inf  23.378  <.0001
 LAR1b - LAR2b   0.7382 0.0774 Inf   9.531  <.0001
 LAR2a - LAR2b  -1.0724 0.0774 Inf -13.846  <.0001

tissue = Petal:
 contrast      estimate     SE  df z.ratio p.value
 LAR1a - LAR1b  -1.3955 0.2520 Inf  -5.531  <.0001
 LAR1a - LAR2a   2.1538 0.2520 Inf   8.536  <.0001
 LAR1a - LAR2b   0.5098 0.2520 Inf   2.020  0.2600
 LAR1b - LAR2a   3.5493 0.2520 Inf  14.066  <.0001
 LAR1b - LAR2b   1.9054 0.2520 Inf   7.551  <.0001
 LAR2a - LAR2b  -1.6440 0.2520 Inf  -6.515  <.0001

tissue = Pollen:
 contrast      estimate     SE  df z.ratio p.value
 LAR1a - LAR1b   0.0000 0.3710 Inf   0.000  1.0000
 LAR1a - LAR2a  -0.0120 0.3710 Inf  -0.032  1.0000
 LAR1a - LAR2b   0.0000 0.3710 Inf   0.000  1.0000
 LAR1b - LAR2a  -0.0120 0.3710 Inf  -0.032  1.0000
 LAR1b - LAR2b   0.0000 0.3710 Inf   0.000  1.0000
 LAR2a - LAR2b   0.0120 0.3710 Inf   0.032  1.0000

tissue = Protoplasts:
 contrast      estimate     SE  df z.ratio p.value
 LAR1a - LAR1b  -0.5329 0.3710 Inf  -1.435  0.9083
 LAR1a - LAR2a   0.4525 0.3710 Inf   1.218  1.0000
 LAR1a - LAR2b  -0.4533 0.3710 Inf  -1.220  1.0000
 LAR1b - LAR2a   0.9853 0.3710 Inf   2.653  0.0479
 LAR1b - LAR2b   0.0796 0.3710 Inf   0.214  1.0000
 LAR2a - LAR2b  -0.9058 0.3710 Inf  -2.439  0.0884

tissue = Roots:
 contrast      estimate     SE  df z.ratio p.value
 LAR1a - LAR1b  -0.7427 0.0647 Inf -11.486  <.0001
 LAR1a - LAR2a   1.4239 0.0647 Inf  22.023  <.0001
 LAR1a - LAR2b   0.6827 0.0647 Inf  10.558  <.0001
 LAR1b - LAR2a   2.1666 0.0647 Inf  33.510  <.0001
 LAR1b - LAR2b   1.4253 0.0647 Inf  22.045  <.0001
 LAR2a - LAR2b  -0.7413 0.0647 Inf -11.465  <.0001

tissue = Seed:
 contrast      estimate     SE  df z.ratio p.value
 LAR1a - LAR1b  -0.5379 0.2520 Inf  -2.132  0.1981
 LAR1a - LAR2a   0.7042 0.2520 Inf   2.791  0.0315
```

| | |
|---|---|
| Degrees-of-freedom method: asymptotic<br>Confidence level used: 0.95 | LAR1a - LAR2b 0.5278 0.2520 Inf 2.092 0.2187<br>LAR1b - LAR2a 1.2421 0.2520 Inf 4.923 <.0001<br>LAR1b - LAR2b 1.0658 0.2520 Inf 4.224 0.0001<br>LAR2a - LAR2b -0.1764 0.2520 Inf -0.699 1.0000<br><br>tissue = Seedlings:<br>contrast estimate SE df z.ratio p.value<br>LAR1a - LAR1b -0.3615 0.3710 Inf -0.973 1.0000<br>LAR1a - LAR2a 2.2299 0.3710 Inf 6.004 <.0001<br>LAR1a - LAR2b 1.9431 0.3710 Inf 5.232 <.0001<br>LAR1b - LAR2a 2.5914 0.3710 Inf 6.977 <.0001<br>LAR1b - LAR2b 2.3046 0.3710 Inf 6.205 <.0001<br>LAR2a - LAR2b -0.2868 0.3710 Inf -0.772 1.0000<br><br>tissue = Shoot Apical:<br>contrast estimate SE df z.ratio p.value<br>LAR1a - LAR1b -0.6486 0.1780 Inf -3.635 0.0017<br>LAR1a - LAR2a 1.9644 0.1780 Inf 11.009 <.0001<br>LAR1a - LAR2b 0.3642 0.1780 Inf 2.041 0.2475<br>LAR1b - LAR2a 2.6130 0.1780 Inf 14.645 <.0001<br>LAR1b - LAR2b 1.0128 0.1780 Inf 5.676 <.0001<br>LAR2a - LAR2b -1.6002 0.1780 Inf -8.968 <.0001<br><br>tissue = Stamen:<br>contrast estimate SE df z.ratio p.value<br>LAR1a - LAR1b -0.4325 0.9100 Inf -0.475 1.0000<br>LAR1a - LAR2a 2.0316 0.9100 Inf 2.233 0.1533<br>LAR1a - LAR2b 1.8043 0.9100 Inf 1.983 0.2841<br>LAR1b - LAR2a 2.4641 0.9100 Inf 2.708 0.0406<br>LAR1b - LAR2b 2.2368 0.9100 Inf 2.459 0.0837<br>LAR2a - LAR2b -0.2273 0.9100 Inf -0.250 1.0000<br><br>tissue = Stem:<br>contrast estimate SE df z.ratio p.value<br>LAR1a - LAR1b -0.4268 0.1210 Inf -3.542 0.0024<br>LAR1a - LAR2a 2.3947 0.1210 Inf 19.873 <.0001<br>LAR1a - LAR2b 0.5679 0.1210 Inf 4.713 <.0001<br>LAR1b - LAR2a 2.8215 0.1210 Inf 23.414 <.0001<br>LAR1b - LAR2b 0.9947 0.1210 Inf 8.254 <.0001<br>LAR2a - LAR2b -1.8268 0.1210 Inf -15.160 <.0001<br><br>tissue = Tetrads:<br>contrast estimate SE df z.ratio p.value<br>LAR1a - LAR1b -0.6570 0.3710 Inf -1.769 0.4615<br>LAR1a - LAR2a 1.3432 0.3710 Inf 3.616 0.0018<br>LAR1a - LAR2b 0.7721 0.3710 Inf 2.079 0.2258<br>LAR1b - LAR2a 2.0002 0.3710 Inf 5.385 <.0001<br>LAR1b - LAR2b 1.4291 0.3710 Inf 3.848 0.0007 |

| | |
|---|---|
| | LAR2a - LAR2b -0.5711 0.3710 Inf -1.538 0.7448<br>tissue = Torus:<br>contrast estimate SE df z.ratio p.value<br>LAR1a - LAR1b -0.1582 0.9100 Inf -0.174 1.0000<br>LAR1a - LAR2a 2.8342 0.9100 Inf 3.115 0.0110<br>LAR1a - LAR2b 1.7032 0.9100 Inf 1.872 0.3671<br>LAR1b - LAR2a 2.9923 0.9100 Inf 3.289 0.0060<br>LAR1b - LAR2b 1.8614 0.9100 Inf 2.046 0.2446<br>LAR2a - LAR2b -1.1309 0.9100 Inf -1.243 1.0000<br>tissue = Uninucleate:<br>contrast estimate SE df z.ratio p.value<br>LAR1a - LAR1b -0.6743 0.3710 Inf -1.815 0.4167<br>LAR1a - LAR2a 1.2712 0.3710 Inf 3.422 0.0037<br>LAR1a - LAR2b 0.5935 0.3710 Inf 1.598 0.6605<br>LAR1b - LAR2a 1.9454 0.3710 Inf 5.238 <.0001<br>LAR1b - LAR2b 1.2678 0.3710 Inf 3.413 0.0039<br>LAR2a - LAR2b -0.6777 0.3710 Inf -1.825 0.4084 |

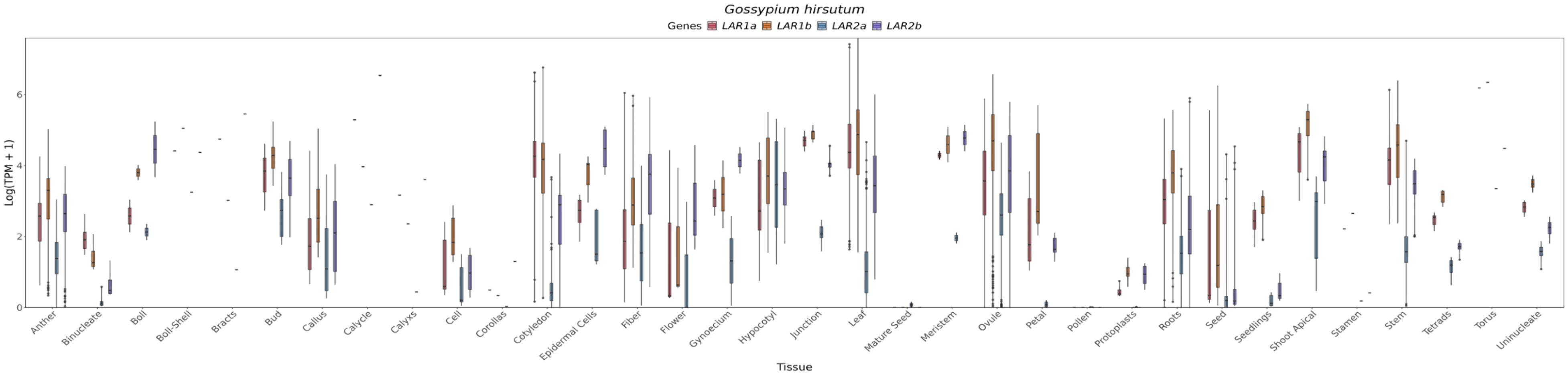

**Figure 3S.** Comparison of the expression of *LAR1a* (red), *LAR1b* (yellow), *LAR2a* (blue) and *LAR2b* (purple) genes in the species *Gossypium hirsutum*.

#### *Actinidia chinensis*

Linear mixed model fit by REML. t-tests use Satterthwaite's method ['lmerModLmerTest']
Formula: TPM_log ~ Genes * tissue + (1 | Sample)
Data: data_long

REML criterion at convergence: 2095.6

Scaled residuals:

| Min | 1Q | Median | 3Q | Max |
|---|---|---|---|---|
| -3.1345 | -0.4801 | -0.0858 | 0.3651 | 5.7218 |

Random effects:

| Groups | Name | Variance | Std.Dev. |
|---|---|---|---|
| Sample | (Intercept) | 0.09163 | 0.3027 |
| Residual | 0.34866 | 0.5905 | |
| Number of obs: | 1080 | groups: | Sample, 270 |

Fixed effects:

| | *Estimate* | *Std. Error* | *df* | *T value* | *Pr(>\|t\|)* | |
|---|---|---|---|---|---|---|
| *(Intercept)* | *3.62247* | *0.16589* | *881.46223* | *21.837* | *<2E-016* | *** |
| *GenesLAR1b* | *-3.53968* | *0.20876* | *747* | *-16.955* | *<2E-016* | *** |
| *GenesLAR1c* | *-2.12177* | *0.20876* | *747* | *-10.164* | *<2E-016* | *** |
| *GenesLAR2* | *-0.7778* | *0.20876* | *747* | *-3.726* | *0.000209* | *** |
| *tissueAxillary bud* | *1.64506* | *0.26748* | *881.46223* | *6.15* | *1.17E-09* | *** |
| *tissueCane* | *1.32631* | *0.68397* | *881.46223* | *1.939* | *0.052802* | . |
| *tissueFlesh* | *-2.35286* | *0.19854* | *881.46223* | *-11.851* | *<2E-016* | *** |
| *tissueFloral bud* | *-0.40476* | *0.27648* | *881.46223* | *-1.464* | *0.143556* | |
| *tissueFlower* | *0.83305* | *0.68397* | *881.46223* | *1.218* | *0.223559* | |
| *tissueFlower bud* | *0.97416* | *0.68397* | *881.46223* | *1.424* | *0.154719* | |
| *tissueFruit* | *-2.41667* | *0.17931* | *881.46223* | *-13.477* | *<2E-016* | *** |
| *tissueFull fruit* | *-2.83403* | *0.31765* | *881.46223* | *-8.922* | *<2E-016* | *** |
| *tissueImmature fruit* | *-1.50145* | *0.49766* | *881.46223* | *-3.017* | *0.002626* | ** |
| *tissueLeaf* | *0.47633* | *0.20795* | *881.46223* | *2.291* | *0.022223* | * |
| *tissueLeaf blade* | *0.37852* | *0.21416* | *881.46223* | *1.767* | *0.077495* | . |
| *tissueMature fruit* | *-1.30208* | *0.41747* | *881.46223* | *-3.119* | *0.001874* | ** |
| *tissuePlant root* | *0.14609* | *0.27648* | *881.46223* | *0.528* | *0.597351* | |
| *tissuePulp* | *-2.10614* | *0.31765* | *881.46223* | *-6.63* | *5.83E-11* | *** |
| *tissueRipe fruit* | *-2.0808* | *0.41747* | *881.46223* | *-4.984* | *7.49E-07* | *** |
| *tissueRoot* | *-0.95254* | *0.2534* | *881.46223* | *-3.759* | *0.000182* | *** |
| *tissueShoot* | *2.0984* | *0.68397* | *881.46223* | *3.068* | *0.002221* | ** |
| *tissueStem* | *-1.17261* | *0.49766* | *881.46223* | *-2.356* | *0.018679* | * |

| | | | | | | |
|---|---|---|---|---|---|---|
| *tissueTerminal bud* | *1.63388* | *0.41747* | *881.46223* | *3.914* | *9.79E-05* | ***** |
| *tissueYoung leaves* | *0.7751* | *0.68397* | *881.46223* | *1.133* | *0.257422* | |
| *GenesLAR1b:tissueAxillary bud* | *-1.59738* | *0.33662* | *747* | *-4.745* | *2.49E-06* | ***** |
| *GenesLAR1c:tissueAxillary bud* | *-0.83576* | *0.33662* | *747* | *-2.483* | *0.013255* | *** |
| *GenesLAR2:tissueAxillary bud* | *-0.23178* | *0.33662* | *747* | *-0.689* | *0.491329* | |
| *GenesLAR1b:tissueCane* | *0.07015* | *0.86075* | *747* | *0.081* | *0.935068* | |
| *GenesLAR1c:tissueCane* | *0.16149* | *0.86075* | *747* | *0.188* | *0.851226* | |
| *GenesLAR2:tissueCane* | *0.09901* | *0.86075* | *747* | *0.115* | *0.908456* | |
| *GenesLAR1b:tissueFlesh* | *2.28408* | *0.24986* | *747* | *9.142* | *<2E-016* | ***** |
| *GenesLAR1c:tissueFlesh* | *1.36766* | *0.24986* | *747* | *5.474* | *6.02E-08* | ***** |
| *GenesLAR2:tissueFlesh* | *0.28253* | *0.24986* | *747* | *1.131* | *0.258521* | |
| *GenesLAR1b:tissueFloral bud* | *0.79027* | *0.34794* | *747* | *2.271* | *0.023414* | *** |
| *GenesLAR1c:tissueFloral bud* | *1.55397* | *0.34794* | *747* | *4.466* | *9.19E-06* | ***** |
| *GenesLAR2:tissueFloral bud* | *0.41707* | *0.34794* | *747* | *1.199* | *0.231032* | |
| *GenesLAR1b:tissueFlower* | *-0.32613* | *0.86075* | *747* | *-0.379* | *0.704881* | |
| *GenesLAR1c:tissueFlower* | *1.68435* | *0.86075* | *747* | *1.957* | *0.05074* | . |
| *GenesLAR2:tissueFlower* | *0.23481* | *0.86075* | *747* | *0.273* | *0.785084* | |
| *GenesLAR1b:tissueFlower bud* | *-0.92048* | *0.86075* | *747* | *-1.069* | *0.285241* | |
| *GenesLAR1c:tissueFlower bud* | *1.72006* | *0.86075* | *747* | *1.998* | *0.046045* | *** |
| *GenesLAR2:tissueFlower bud* | *-0.1377* | *0.86075* | *747* | *-0.16* | *0.872943* | |
| *GenesLAR1b:tissueFruit* | *2.51757* | *0.22566* | *747* | *11.156* | *<2E-016* | ***** |
| *GenesLAR1c:tissueFruit* | *1.31038* | *0.22566* | *747* | *5.807* | *9.41E-09* | ***** |
| *GenesLAR2:tissueFruit* | *0.7733* | *0.22566* | *747* | *3.427* | *0.000644* | ***** |
| *GenesLAR1b:tissueFull fruit* | *2.75124* | *0.39975* | *747* | *6.882* | *1.25E-11* | ***** |
| *GenesLAR1c:tissueFull fruit* | *3.68874* | *0.39975* | *747* | *9.228* | *<2E-016* | ***** |
| *GenesLAR2:tissueFull fruit* | *1.14855* | *0.39975* | *747* | *2.873* | *0.004179* | **** |
| *GenesLAR1b:tissueImmature fruit* | *1.41865* | *0.62629* | *747* | *2.265* | *0.023788* | *** |
| *GenesLAR1c:tissueImmature fruit* | *3.00364* | *0.62629* | *747* | *4.796* | *1.96E-06* | ***** |
| *GenesLAR2:tissueImmature fruit* | *0.42284* | *0.62629* | *747* | *0.675* | *0.499789* | |
| *GenesLAR1b:tissueLeaf* | *-0.3111* | *0.2617* | *747* | *-1.189* | *0.23491* | |
| *GenesLAR1c:tissueLeaf* | *0.47117* | *0.2617* | *747* | *1.8* | *0.072197* | . |
| *GenesLAR2:tissueLeaf* | *-0.43898* | *0.2617* | *747* | *-1.677* | *0.093881* | . |
| *GenesLAR1b:tissueLeaf blade* | *-0.39489* | *0.26951* | *747* | *-1.465* | *0.143283* | |
| *GenesLAR1c:tissueLeaf blade* | *0.2207* | *0.26951* | *747* | *0.819* | *0.413123* | |
| *GenesLAR2:tissueLeaf blade* | *-1.45963* | *0.26951* | *747* | *-5.416* | *8.23E-08* | ***** |
| *GenesLAR1b:tissueMature fruit* | *1.22347* | *0.52538* | *747* | *2.329* | *0.020139* | *** |
| *GenesLAR1c:tissueMature fruit* | *1.19436* | *0.52538* | *747* | *2.273* | *0.023289* | *** |
| *GenesLAR2:tissueMature fruit* | *0.55707* | *0.52538* | *747* | *1.06* | *0.289344* | |
| *GenesLAR1b:tissuePlant root* | *1.22912* | *0.34794* | *747* | *3.533* | *0.000437* | ***** |
| *GenesLAR1c:tissuePlant root* | *0.22126* | *0.34794* | *747* | *0.636* | *0.525019* | |
| *GenesLAR2:tissuePlant root* | *-0.6733* | *0.34794* | *747* | *-1.935* | *0.053354* | . |
| *GenesLAR1b:tissuePulp* | *2.02335* | *0.39975* | *747* | *5.062* | *5.24E-07* | ***** |
| *GenesLAR1c:tissuePulp* | *1.00726* | *0.39975* | *747* | *2.52* | *0.011953* | *** |
| *GenesLAR2:tissuePulp* | *-0.27943* | *0.39975* | *747* | *-0.699* | *0.484761* | |
| *GenesLAR1b:tissueRipe fruit* | *2.00368* | *0.52538* | *747* | *3.814* | *0.000148* | ***** |
| *GenesLAR1c:tissueRipe fruit* | *0.60233* | *0.52538* | *747* | *1.146* | *0.251964* | |

| | | | | | | |
|---|---|---|---|---|---|---|
| *GenesLAR2:tissueRipe fruit* | *0.53542* | *0.52538* | *747* | *1.019* | *0.308476* | |
| *GenesLAR1b:tissueRoot* | *1.85218* | *0.31889* | *747* | *5.808* | *9.34E-09* | ***** |
| *GenesLAR1c:tissueRoot* | *1.00611* | *0.31889* | *747* | *3.155* | *0.001669* | **** |
| *GenesLAR2:tissueRoot* | *0.44656* | *0.31889* | *747* | *1.4* | *0.161824* | |
| *GenesLAR1b:tissueShoot* | *-0.27922* | *0.86075* | *747* | *-0.324* | *0.745731* | |
| *GenesLAR1c:tissueShoot* | *-0.13677* | *0.86075* | *747* | *-0.159* | *0.87379* | |
| *GenesLAR2:tissueShoot* | *-0.94476* | *0.86075* | *747* | *-1.098* | *0.272737* | |
| *GenesLAR1b:tissueStem* | *1.10002* | *0.62629* | *747* | *1.756* | *0.07943* | . |
| *GenesLAR1c:tissueStem* | *0.04657* | *0.62629* | *747* | *0.074* | *0.94074* | |
| *GenesLAR2:tissueStem* | *0.55313* | *0.62629* | *747* | *0.883* | *0.377417* | |
| *GenesLAR1b:tissueTerminal bud* | *-1.07634* | *0.52538* | *747* | *-2.049* | *0.04084* | *** |
| *GenesLAR1c:tissueTerminal bud* | *1.01347* | *0.52538* | *747* | *1.929* | *0.054105* | . |
| *GenesLAR2:tissueTerminal bud* | *-0.30105* | *0.52538* | *747* | *-0.573* | *0.566803* | |
| *GenesLAR1b:tissueYoung leaves* | *-0.10578* | *0.86075* | *747* | *-0.123* | *0.902225* | |
| *GenesLAR1c:tissueYoung leaves* | *1.27785* | *0.86075* | *747* | *1.485* | *0.138079* | |
| *GenesLAR2:tissueYoung leaves* | *-1.30573* | *0.86075* | *747* | *-1.517* | *0.1297* | |

Signif. codes: 0 ‘***’ 0.001 ‘**’ 0.01 ‘*’ 0.05 ‘.’ 0.1 ‘ ’ 1

**Table 10S.** Estimated marginal means and pairwise comparison for the species *Actinidia chinensis*. Degrees-of-freedom method: asymptotic
P value adjustment: bonferroni method for 6 tests

| ***Estimated Marginal Means*** | ***Contrasts*** |
|---|---|
| tissue = All plant: | tissue = All plant: |
| Genes emmean SE df lower.CL upper.CL | contrast estimate SE df t.ratio p.value |
| LAR1a 3.62247 0.1660 881 3.29689 3.948 | LAR1a - LAR1b 3.53968 0.2090 747 16.955 <.0001 |
| LAR1b 0.08279 0.1660 881 -0.24278 0.408 | LAR1a - LAR1c 2.12177 0.2090 747 10.164 <.0001 |
| LAR1c 1.50070 0.1660 881 1.17512 1.826 | LAR1a - LAR2 0.77780 0.2090 747 3.726 0.0013 |
| LAR2 2.84467 0.1660 881 2.51910 3.170 | LAR1b - LAR1c -1.41790 0.2090 747 -6.792 <.0001 |
| | LAR1b - LAR2 -2.76188 0.2090 747 -13.230 <.0001 |
| tissue = Axillary bud: | LAR1c - LAR2 -1.34398 0.2090 747 -6.438 <.0001 |
| Genes emmean SE df lower.CL upper.CL | |
| LAR1a 5.26753 0.2100 881 4.85570 5.679 | tissue = Axillary bud: |
| LAR1b 0.13048 0.2100 881 -0.28135 0.542 | contrast estimate SE df t.ratio p.value |
| LAR1c 2.31000 0.2100 881 1.89817 2.722 | LAR1a - LAR1b 5.13705 0.2640 747 19.454 <.0001 |
| LAR2 4.25796 0.2100 881 3.84613 4.670 | LAR1a - LAR1c 2.95753 0.2640 747 11.200 <.0001 |
| | LAR1a - LAR2 1.00957 0.2640 747 3.823 0.0009 |
| tissue = Cane: | LAR1b - LAR1c -2.17952 0.2640 747 -8.254 <.0001 |
| Genes emmean SE df lower.CL upper.CL | LAR1b - LAR2 -4.12748 0.2640 747 -15.630 <.0001 |
| LAR1a 4.94878 0.6640 881 3.64647 6.251 | LAR1c - LAR2 -1.94796 0.2640 747 -7.377 <.0001 |
| LAR1b 1.47925 0.6640 881 0.17694 2.782 | |
| LAR1c 2.98850 0.6640 881 1.68619 4.291 | tissue = Cane: |
| LAR2 4.26999 0.6640 881 2.96768 5.572 | contrast estimate SE df t.ratio p.value |
| | LAR1a - LAR1b 3.46953 0.8350 747 4.155 0.0002 |
| tissue = Flesh: | LAR1a - LAR1c 1.96028 0.8350 747 2.347 0.1150 |
| Genes emmean SE df lower.CL upper.CL | LAR1a - LAR2 0.67879 0.8350 747 0.813 1.0000 |
| LAR1a 1.26961 0.1090 881 1.05551 1.484 | LAR1b - LAR1c -1.50925 0.8350 747 -1.807 0.4266 |
| LAR1b 0.01401 0.1090 881 -0.20009 0.228 | LAR1b - LAR2 -2.79074 0.8350 747 -3.342 0.0052 |
| LAR1c 0.51550 0.1090 881 0.30140 0.730 | LAR1c - LAR2 -1.28149 0.8350 747 -1.535 0.7518 |

```
 LAR2  0.77434 0.1090 881  0.56024    0.988

tissue = Floral bud:
 Genes  emmean     SE  df lower.CL upper.CL
 LAR1a 3.21771 0.2210 881  2.78361    3.652
 LAR1b 0.46831 0.2210 881  0.03420    0.902
 LAR1c 2.64991 0.2210 881  2.21580    3.084
 LAR2  2.85698 0.2210 881  2.42288    3.291

tissue = Flower:
 Genes  emmean     SE  df lower.CL upper.CL
 LAR1a 4.45552 0.6640 881  3.15321    5.758
 LAR1b 0.58972 0.6640 881 -0.71259    1.892
 LAR1c 4.01810 0.6640 881  2.71579    5.320
 LAR2  3.91254 0.6640 881  2.61023    5.215

tissue = Flower bud:
 Genes  emmean     SE  df lower.CL upper.CL
 LAR1a 4.59663 0.6640 881  3.29432    5.899
 LAR1b 0.13647 0.6640 881 -1.16584    1.439
 LAR1c 4.19491 0.6640 881  2.89260    5.497
 LAR2  3.68113 0.6640 881  2.37882    4.983

tissue = Fruit:
 Genes  emmean     SE  df lower.CL upper.CL
 LAR1a 1.20580 0.0681 881  1.07219    1.339
 LAR1b 0.18369 0.0681 881  0.05008    0.317
 LAR1c 0.39441 0.0681 881  0.26079    0.528
 LAR2  1.20131 0.0681 881  1.06769    1.335

tissue = Full fruit:
 Genes  emmean     SE  df lower.CL upper.CL
 LAR1a 0.78844 0.2710 881  0.25677    1.320
 LAR1b 0.00000 0.2710 881 -0.53167    0.532
 LAR1c 2.35541 0.2710 881  1.82374    2.887
 LAR2  1.15919 0.2710 881  0.62753    1.691

tissue = Immature fruit:
 Genes  emmean     SE  df lower.CL upper.CL
 LAR1a 2.12102 0.4690 881  1.20015    3.042
 LAR1b 0.00000 0.4690 881 -0.92087    0.921
 LAR1c 3.00289 0.4690 881  2.08202    3.924
 LAR2  1.76607 0.4690 881  0.84519    2.687

tissue = Leaf:
 Genes  emmean     SE  df lower.CL upper.CL
 LAR1a 4.09880 0.1250 881  3.85268    4.345
 LAR1b 0.24802 0.1250 881  0.00191    0.494
```

```
tissue = Flesh:
 contrast      estimate     SE  df t.ratio p.value
 LAR1a - LAR1b  1.25560 0.1370 747   9.146  <.0001
 LAR1a - LAR1c  0.75411 0.1370 747   5.493  <.0001
 LAR1a - LAR2   0.49527 0.1370 747   3.608  0.0020
 LAR1b - LAR1c -0.50149 0.1370 747  -3.653  0.0017
 LAR1b - LAR2  -0.76033 0.1370 747  -5.538  <.0001
 LAR1c - LAR2  -0.25884 0.1370 747  -1.885  0.3585

tissue = Floral bud:
 contrast      estimate     SE  df t.ratio p.value
 LAR1a - LAR1b  2.74941 0.2780 747   9.877  <.0001
 LAR1a - LAR1c  0.56781 0.2780 747   2.040  0.2503
 LAR1a - LAR2   0.36073 0.2780 747   1.296  1.0000
 LAR1b - LAR1c -2.18160 0.2780 747  -7.838  <.0001
 LAR1b - LAR2  -2.38868 0.2780 747  -8.582  <.0001
 LAR1c - LAR2  -0.20708 0.2780 747  -0.744  1.0000

tissue = Flower:
 contrast      estimate     SE  df t.ratio p.value
 LAR1a - LAR1b  3.86580 0.8350 747   4.629  <.0001
 LAR1a - LAR1c  0.43743 0.8350 747   0.524  1.0000
 LAR1a - LAR2   0.54299 0.8350 747   0.650  1.0000
 LAR1b - LAR1c -3.42838 0.8350 747  -4.106  0.0003
 LAR1b - LAR2  -3.32282 0.8350 747  -3.979  0.0005
 LAR1c - LAR2   0.10556 0.8350 747   0.126  1.0000

tissue = Flower bud:
 contrast      estimate     SE  df t.ratio p.value
 LAR1a - LAR1b  4.46016 0.8350 747   5.341  <.0001
 LAR1a - LAR1c  0.40172 0.8350 747   0.481  1.0000
 LAR1a - LAR2   0.91550 0.8350 747   1.096  1.0000
 LAR1b - LAR1c -4.05844 0.8350 747  -4.860  <.0001
 LAR1b - LAR2  -3.54466 0.8350 747  -4.245  0.0001
 LAR1c - LAR2   0.51378 0.8350 747   0.615  1.0000

tissue = Fruit:
 contrast      estimate     SE  df t.ratio p.value
 LAR1a - LAR1b  1.02211 0.0857 747  11.930  <.0001
 LAR1a - LAR1c  0.81139 0.0857 747   9.471  <.0001
 LAR1a - LAR2   0.00449 0.0857 747   0.052  1.0000
 LAR1b - LAR1c -0.21072 0.0857 747  -2.459  0.0848
 LAR1b - LAR2  -1.01761 0.0857 747 -11.878  <.0001
 LAR1c - LAR2  -0.80690 0.0857 747  -9.418  <.0001

tissue = Full fruit:
 contrast      estimate     SE  df t.ratio p.value
```

```
 LAR1c 2.44819 0.1250 881  2.20208    2.694
 LAR2  2.88202 0.1250 881  2.63591    3.128

tissue = Leaf blade:
 Genes  emmean     SE  df lower.CL upper.CL
 LAR1a 4.00099 0.1350 881  3.73516    4.267
 LAR1b 0.06642 0.1350 881 -0.19942    0.332
 LAR1c 2.09991 0.1350 881  1.83408    2.366
 LAR2  1.76356 0.1350 881  1.49773    2.029

tissue = Mature fruit:
 Genes  emmean     SE  df lower.CL upper.CL
 LAR1a 2.32039 0.3830 881  1.56850    3.072
 LAR1b 0.00418 0.3830 881 -0.74771    0.756
 LAR1c 1.39297 0.3830 881  0.64108    2.145
 LAR2  2.09966 0.3830 881  1.34777    2.852

tissue = Plant root:
 Genes  emmean     SE  df lower.CL upper.CL
 LAR1a 3.76856 0.2210 881  3.33446    4.203
 LAR1b 1.45801 0.2210 881  1.02391    1.892
 LAR1c 1.86805 0.2210 881  1.43395    2.302
 LAR2  2.31746 0.2210 881  1.88336    2.752

tissue = Pulp:
 Genes  emmean     SE  df lower.CL upper.CL
 LAR1a 1.51633 0.2710 881  0.98466    2.048
 LAR1b 0.00000 0.2710 881 -0.53167    0.532
 LAR1c 0.40181 0.2710 881 -0.12985    0.933
 LAR2  0.45910 0.2710 881 -0.07257    0.991

tissue = Ripe fruit:
 Genes  emmean     SE  df lower.CL upper.CL
 LAR1a 1.54167 0.3830 881  0.78978    2.294
 LAR1b 0.00568 0.3830 881 -0.74621    0.758
 LAR1c 0.02223 0.3830 881 -0.72966    0.774
 LAR2  1.29929 0.3830 881  0.54741    2.051

tissue = Root:
 Genes  emmean     SE  df lower.CL upper.CL
 LAR1a 2.66993 0.1920 881  2.29398    3.046
 LAR1b 0.98243 0.1920 881  0.60648    1.358
 LAR1c 1.55426 0.1920 881  1.17831    1.930
 LAR2  2.33869 0.1920 881  1.96274    2.715

tissue = Shoot:
 Genes  emmean     SE  df lower.CL upper.CL
 LAR1a 5.72087 0.6640 881  4.41856    7.023
```

```
 LAR1a - LAR1b  0.78844 0.3410 747   2.313  0.1260
 LAR1a - LAR1c -1.56696 0.3410 747  -4.596  <.0001
 LAR1a - LAR2  -0.37075 0.3410 747  -1.088  1.0000
 LAR1b - LAR1c -2.35541 0.3410 747  -6.909  <.0001
 LAR1b - LAR2  -1.15919 0.3410 747  -3.400  0.0043
 LAR1c - LAR2   1.19621 0.3410 747   3.509  0.0029

tissue = Immature fruit:
 contrast      estimate     SE  df t.ratio p.value
 LAR1a - LAR1b  2.12102 0.5900 747   3.592  0.0021
 LAR1a - LAR1c -0.88187 0.5900 747  -1.493  0.8144
 LAR1a - LAR2   0.35496 0.5900 747   0.601  1.0000
 LAR1b - LAR1c -3.00289 0.5900 747  -5.086  <.0001
 LAR1b - LAR2  -1.76607 0.5900 747  -2.991  0.0172
 LAR1c - LAR2   1.23683 0.5900 747   2.095  0.2192

tissue = Leaf:
 contrast      estimate     SE  df t.ratio p.value
 LAR1a - LAR1b  3.85078 0.1580 747  24.401  <.0001
 LAR1a - LAR1c  1.65061 0.1580 747  10.459  <.0001
 LAR1a - LAR2   1.21677 0.1580 747   7.710  <.0001
 LAR1b - LAR1c -2.20017 0.1580 747 -13.942  <.0001
 LAR1b - LAR2  -2.63400 0.1580 747 -16.691  <.0001
 LAR1c - LAR2  -0.43383 0.1580 747  -2.749  0.0367

tissue = Leaf blade:
 contrast      estimate     SE  df t.ratio p.value
 LAR1a - LAR1b  3.93457 0.1700 747  23.083  <.0001
 LAR1a - LAR1c  1.90108 0.1700 747  11.153  <.0001
 LAR1a - LAR2   2.23743 0.1700 747  13.126  <.0001
 LAR1b - LAR1c -2.03349 0.1700 747 -11.930  <.0001
 LAR1b - LAR2  -1.69714 0.1700 747  -9.957  <.0001
 LAR1c - LAR2   0.33635 0.1700 747   1.973  0.2930

tissue = Mature fruit:
 contrast      estimate     SE  df t.ratio p.value
 LAR1a - LAR1b  2.31621 0.4820 747   4.804  <.0001
 LAR1a - LAR1c  0.92742 0.4820 747   1.924  0.3287
 LAR1a - LAR2   0.22073 0.4820 747   0.458  1.0000
 LAR1b - LAR1c -1.38879 0.4820 747  -2.881  0.0245
 LAR1b - LAR2  -2.09548 0.4820 747  -4.346  0.0001
 LAR1c - LAR2  -0.70669 0.4820 747  -1.466  0.8588

tissue = Plant root:
 contrast      estimate     SE  df t.ratio p.value
 LAR1a - LAR1b  2.31055 0.2780 747   8.301  <.0001
 LAR1a - LAR1c  1.90051 0.2780 747   6.828  <.0001
 LAR1a - LAR2   1.45110 0.2780 747   5.213  <.0001
```

```
 LAR1b 1.90197 0.6640 881  0.59966    3.204
 LAR1c 3.46232 0.6640 881  2.16001    4.765
 LAR2  3.99832 0.6640 881  2.69601    5.301

tissue = Stem:
 Genes  emmean     SE  df lower.CL upper.CL
 LAR1a 2.44986 0.4690 881  1.52899    3.371
 LAR1b 0.01020 0.4690 881 -0.91067    0.931
 LAR1c 0.37466 0.4690 881 -0.54621    1.296
 LAR2  2.22520 0.4690 881  1.30433    3.146

tissue = Terminal bud:
 Genes  emmean     SE  df lower.CL upper.CL
 LAR1a 5.25635 0.3830 881  4.50446    6.008
 LAR1b 0.64033 0.3830 881 -0.11156    1.392
 LAR1c 4.14805 0.3830 881  3.39616    4.900
 LAR2  4.17750 0.3830 881  3.42561    4.929

tissue = Young leaves:
 Genes  emmean     SE  df lower.CL upper.CL
 LAR1a 4.39757 0.6640 881  3.09525    5.700
 LAR1b 0.75211 0.6640 881 -0.55020    2.054
 LAR1c 3.55364 0.6640 881  2.25133    4.856
 LAR2  2.31404 0.6640 881  1.01173    3.616

Degrees-of-freedom method: kenward-roger
Confidence level used: 0.95
```

```
 LAR1b - LAR1c -0.41004 0.2780 747  -1.473  0.8469
 LAR1b - LAR2  -0.85945 0.2780 747  -3.088  0.0126
 LAR1c - LAR2  -0.44941 0.2780 747  -1.615  0.6410

tissue = Pulp:
 contrast      estimate     SE  df t.ratio p.value
 LAR1a - LAR1b  1.51633 0.3410 747   4.448  0.0001
 LAR1a - LAR1c  1.11451 0.3410 747   3.269  0.0068
 LAR1a - LAR2   1.05723 0.3410 747   3.101  0.0120
 LAR1b - LAR1c -0.40181 0.3410 747  -1.179  1.0000
 LAR1b - LAR2  -0.45910 0.3410 747  -1.347  1.0000
 LAR1c - LAR2  -0.05728 0.3410 747  -0.168  1.0000

tissue = Ripe fruit:
 contrast      estimate     SE  df t.ratio p.value
 LAR1a - LAR1b  1.53599 0.4820 747   3.186  0.0090
 LAR1a - LAR1c  1.51944 0.4820 747   3.152  0.0101
 LAR1a - LAR2   0.24238 0.4820 747   0.503  1.0000
 LAR1b - LAR1c -0.01655 0.4820 747  -0.034  1.0000
 LAR1b - LAR2  -1.29362 0.4820 747  -2.683  0.0447
 LAR1c - LAR2  -1.27706 0.4820 747  -2.649  0.0495

tissue = Root:
 contrast      estimate     SE  df t.ratio p.value
 LAR1a - LAR1b  1.68750 0.2410 747   7.000  <.0001
 LAR1a - LAR1c  1.11567 0.2410 747   4.628  <.0001
 LAR1a - LAR2   0.33124 0.2410 747   1.374  1.0000
 LAR1b - LAR1c -0.57183 0.2410 747  -2.372  0.1076
 LAR1b - LAR2  -1.35626 0.2410 747  -5.626  <.0001
 LAR1c - LAR2  -0.78443 0.2410 747  -3.254  0.0071

tissue = Shoot:
 contrast      estimate     SE  df t.ratio p.value
 LAR1a - LAR1b  3.81890 0.8350 747   4.573  <.0001
 LAR1a - LAR1c  2.25855 0.8350 747   2.705  0.0420
 LAR1a - LAR2   1.72255 0.8350 747   2.063  0.2369
 LAR1b - LAR1c -1.56035 0.8350 747  -1.869  0.3725
 LAR1b - LAR2  -2.09634 0.8350 747  -2.510  0.0736
 LAR1c - LAR2  -0.53599 0.8350 747  -0.642  1.0000

tissue = Stem:
 contrast      estimate     SE  df t.ratio p.value
 LAR1a - LAR1b  2.43966 0.5900 747   4.132  0.0002
 LAR1a - LAR1c  2.07520 0.5900 747   3.514  0.0028
 LAR1a - LAR2   0.22466 0.5900 747   0.380  1.0000
 LAR1b - LAR1c -0.36446 0.5900 747  -0.617  1.0000
 LAR1b - LAR2  -2.21500 0.5900 747  -3.751  0.0011
 LAR1c - LAR2  -1.85054 0.5900 747  -3.134  0.0108
```

| | |
|---|---|
| | tissue = Terminal bud:<br>contrast estimate SE df t.ratio p.value<br>LAR1a - LAR1b 4.61602 0.4820 747 9.574 <.0001<br>LAR1a - LAR1c 1.10831 0.4820 747 2.299 0.1307<br>LAR1a - LAR2 1.07885 0.4820 747 2.238 0.1532<br>LAR1b - LAR1c -3.50771 0.4820 747 -7.276 <.0001<br>LAR1b - LAR2 -3.53717 0.4820 747 -7.337 <.0001<br>LAR1c - LAR2 -0.02945 0.4820 747 -0.061 1.0000<br><br>tissue = Young leaves:<br>contrast estimate SE df t.ratio p.value<br>LAR1a - LAR1b 3.64546 0.8350 747 4.366 0.0001<br>LAR1a - LAR1c 0.84392 0.8350 747 1.011 1.0000<br>LAR1a - LAR2 2.08353 0.8350 747 2.495 0.0768<br>LAR1b - LAR1c -2.80153 0.8350 747 -3.355 0.0050<br>LAR1b - LAR2 -1.56193 0.8350 747 -1.870 0.3709<br>LAR1c - LAR2 1.23961 0.8350 747 1.484 0.8287 |

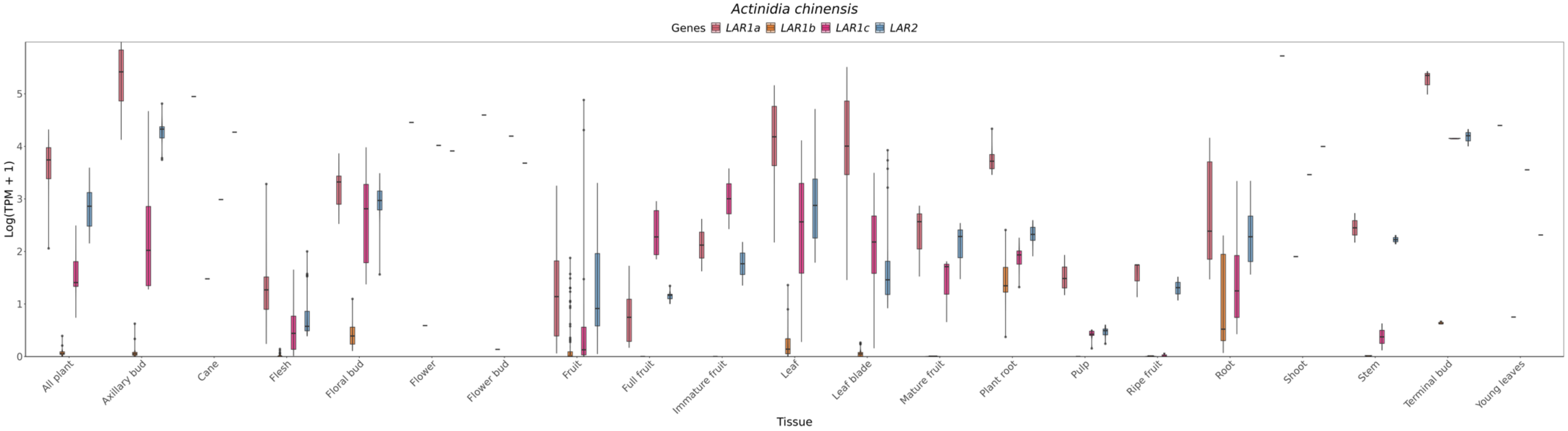

**Figure 4S.** Comparison of the expression of *LAR1a* (red), *LAR1b* (orange), *LAR1c* (pink) and *LAR2a* (blue) genes in the species *Actinidia chinensis*.

#### *Eucalyptus grandis*

Linear mixed model fit by REML. t-tests use Satterthwaite's method ['lmerModLmerTest']
Formula: TPM_log ~ Genes * tissue + (1 | Sample)
Data: data_long

REML criterion at convergence: 961.9

Scaled residuals:

| Min | 1Q | Median | 3Q | Max |
|---|---|---|---|---|
| -4.069 | -0.3202 | 0 | 0.3451 | 2.2926 |

Random effects:

| Groups | Name | Variance | Std.Dev. |
|---|---|---|---|
| Sample | (Intercept) | 0.1334 | 0.3652 |
| Residual | 0.4166 | 0.6454 | |
| Number of obs: | 447 | groups: | Sample, 149 |

Fixed effects:

| | *Estimate* | *Std. Error* | *df* | *T-value* | *Pr(>\|t\|)* | |
|---|---|---|---|---|---|---|
| *(Intercept)* | *3.171845* | *0.370792* | *367.738584* | *8.554* | *3.25E-16* | *** |
| *GenesLAR2a* | *-3.137178* | *0.456381* | *274.000003* | *-6.874* | *4.21E-11* | *** |
| *GenesLAR2b* | *-2.928155* | *0.456381* | *274.000003* | *-6.416* | *6.12E-10* | *** |
| *tissueControl root* | *0.945845* | *0.403093* | *367.738583* | *2.346* | *0.019482* | * |
| *tissueEctomycorrhiza* | *0.622535* | *0.424017* | *367.738583* | *1.468* | *0.142909* | |
| *tissueFruit bud* | *1.434466* | *0.829116* | *367.738581* | *1.73* | *0.084449* | . |
| *tissueLeaf* | *1.115323* | *0.829116* | *367.738581* | *1.345* | *0.179391* | |
| *tissueLeaf adult* | *1.233245* | *0.566394* | *367.738582* | *2.177* | *0.030089* | * |
| *tissueLeaf and twigs* | *-0.834142* | *0.428153* | *367.738583* | *-1.948* | *0.052148* | . |
| *tissueLeaf juvenile* | *0.863925* | *0.566394* | *367.738582* | *1.525* | *0.128042* | |
| *tissueLeaf midrib* | *-0.219222* | *0.47869* | *367.738582* | *-0.458* | *0.647251* | |
| *tissueMixed* | *0.643171* | *0.64223* | *367.738581* | *1.001* | *0.31726* | |
| *tissueStem* | *1.495982* | *0.381542* | *367.738583* | *3.921* | *0.000105* | *** |
| *tissueXylem* | *0.252605* | *0.420438* | *367.738583* | *0.601* | *0.548335* | |
| *GenesLAR2a:tissueControl root* | *1.091958* | *0.496139* | *274.000003* | *2.201* | *0.028576* | * |
| *GenesLAR2b:tissueControl root* | *1.324942* | *0.496139* | *274.000003* | *2.671* | *0.008026* | ** |
| *GenesLAR2a:tissueEctomycorrhiza* | *0.761166* | *0.521892* | *274.000003* | *1.458* | *0.145855* | |
| *GenesLAR2b:tissueEctomycorrhiza* | *0.806413* | *0.521892* | *274.000003* | *1.545* | *0.123458* | |
| *GenesLAR2a:tissueFruit bud* | *0.918013* | *1.020499* | *274.000003* | *0.9* | *0.369138* | |
| *GenesLAR2b:tissueFruit bud* | *1.039574* | *1.020499* | *274.000003* | *1.019* | *0.309248* | |
| *GenesLAR2a:tissueLeaf* | *-1.11748* | *1.020499* | *274.000003* | *-1.095* | *0.274464* | |
| *GenesLAR2b:tissueLeaf* | *0.831917* | *1.020499* | *274.000003* | *0.815* | *0.415663* | |

| | | | | | | |
|---|---|---|---|---|---|---|
| *GenesLAR2a:tissueLeaf adult* | *-1.267912* | *0.697134* | *274.000003* | *-1.819* | *0.070041* | . |
| *GenesLAR2b:tissueLeaf adult* | *-0.917419* | *0.697134* | *274.000003* | *-1.316* | *0.189278* | |
| *GenesLAR2a:tissueLeaf and twigs* | *0.918518* | *0.526984* | *274.000003* | *1.743* | *0.08246* | . |
| *GenesLAR2b:tissueLeaf and twigs* | *0.608164* | *0.526984* | *274.000003* | *1.154* | *0.249486* | |
| *GenesLAR2a:tissueLeaf juvenile* | *-0.599881* | *0.697134* | *274.000003* | *-0.86* | *0.390268* | |
| *GenesLAR2b:tissueLeaf juvenile* | *-0.869701* | *0.697134* | *274.000003* | *-1.248* | *0.213265* | |
| *GenesLAR2a:tissueLeaf midrib* | *1.724201* | *0.589186* | *274.000003* | *2.926* | *0.003716* | ** |
| *GenesLAR2b:tissueLeaf midrib* | *1.663536* | *0.589186* | *274.000003* | *2.823* | *0.005099* | ** |
| *GenesLAR2a:tissueMixed* | *-0.429915* | *0.790475* | *274.000003* | *-0.544* | *0.586974* | |
| *GenesLAR2b:tissueMixed* | *-0.838613* | *0.790475* | *274.000003* | *-1.061* | *0.289671* | |
| *GenesLAR2a:tissueStem* | *-0.190143* | *0.469612* | *274.000003* | *-0.405* | *0.685872* | |
| *GenesLAR2b:tissueStem* | *2.046957* | *0.469612* | *274.000003* | *4.359* | *1.85E-05* | *** |
| *GenesLAR2a:tissueXylem* | *0.001223* | *0.517488* | *274.000003* | *0.002* | *0.998115* | |
| *GenesLAR2b:tissueXylem* | *-0.226549* | *0.517488* | *274.000003* | *-0.438* | *0.661887* | |

Signif. codes: 0 '***' 0.001 '**' 0.01 '*' 0.05 '.' 0.1 ' ' 1

**Table 11S.** Estimated marginal means and pairwise comparison for the species *Eucalyptus grandis*. Degrees-of-freedom method: asymptotic
P value adjustment: bonferroni method for 3 tests.

| ***Estimated Marginal Means*** | ***Contrasts*** |
|---|---|
| tissue = Cambium:<br>Genes emmean SE df lower.CL upper.CL<br>LAR1 3.1718 0.3710 368 2.443 3.901<br>LAR2a 0.0347 0.3710 368 -0.694 0.764<br>LAR2b 0.2437 0.3710 368 -0.485 0.973 | tissue = Cambium:<br>contrast estimate SE df t.ratio p.value<br>LAR1 - LAR2a 3.1372 0.456 274 6.874 <.0001<br>LAR1 - LAR2b 2.9282 0.456 274 6.416 <.0001<br>LAR2a - LAR2b -0.2090 0.456 274 -0.458 1.0000 |
| tissue = Control root:<br>Genes emmean SE df lower.CL upper.CL<br>LAR1 4.1177 0.1580 368 3.807 4.429<br>LAR2a 2.0725 0.1580 368 1.762 2.383<br>LAR2b 2.5145 0.1580 368 2.204 2.825 | tissue = Control root:<br>contrast estimate SE df t.ratio p.value<br>LAR1 - LAR2a 2.0452 0.195 274 10.510 <.0001<br>LAR1 - LAR2b 1.6032 0.195 274 8.238 <.0001<br>LAR2a - LAR2b -0.4420 0.195 274 -2.271 0.0717 |
| tissue = Ectomycorrhiza:<br>Genes emmean SE df lower.CL upper.CL<br>LAR1 3.7944 0.2060 368 3.390 4.199<br>LAR2a 1.4184 0.2060 368 1.014 1.823<br>LAR2b 1.6726 0.2060 368 1.268 2.077 | tissue = Ectomycorrhiza:<br>contrast estimate SE df t.ratio p.value<br>LAR1 - LAR2a 2.3760 0.253 274 9.386 <.0001<br>LAR1 - LAR2b 2.1217 0.253 274 8.381 <.0001<br>LAR2a - LAR2b -0.2543 0.253 274 -1.004 0.9482 |
| tissue = Fruit bud:<br>Genes emmean SE df lower.CL upper.CL<br>LAR1 4.6063 0.7420 368 3.148 6.065<br>LAR2a 2.3871 0.7420 368 0.929 3.845<br>LAR2b 2.7177 0.7420 368 1.259 4.176 | tissue = Fruit bud:<br>contrast estimate SE df t.ratio p.value<br>LAR1 - LAR2a 2.2192 0.913 274 2.431 0.0471<br>LAR1 - LAR2b 1.8886 0.913 274 2.069 0.1184<br>LAR2a - LAR2b -0.3306 0.913 274 -0.362 1.0000 |
| tissue = Leaf:<br>Genes emmean SE df lower.CL upper.CL<br>LAR1 4.2872 0.7420 368 2.829 5.745<br>LAR2a 0.0325 0.7420 368 -1.426 1.491 | tissue = Leaf:<br>contrast estimate SE df t.ratio p.value<br>LAR1 - LAR2a 4.2547 0.913 274 4.661 <.0001<br>LAR1 - LAR2b 2.0962 0.913 274 2.297 0.0672 |

```
 LAR2b 2.1909 0.7420 368    0.733    3.649

tissue = Leaf adult:
 Genes emmean     SE  df lower.CL upper.CL
 LAR1  4.4051 0.4280 368    3.563    5.247
 LAR2a 0.0000 0.4280 368   -0.842    0.842
 LAR2b 0.5595 0.4280 368   -0.282    1.401

tissue = Leaf and twigs:
 Genes emmean     SE  df lower.CL upper.CL
 LAR1  2.3377 0.2140 368    1.917    2.759
 LAR2a 0.1190 0.2140 368   -0.302    0.540
 LAR2b 0.0177 0.2140 368   -0.403    0.439

tissue = Leaf juvenile:
 Genes emmean     SE  df lower.CL upper.CL
 LAR1  4.0358 0.4280 368    3.194    4.878
 LAR2a 0.2987 0.4280 368   -0.543    1.141
 LAR2b 0.2379 0.4280 368   -0.604    1.080

tissue = Leaf midrib:
 Genes emmean     SE  df lower.CL upper.CL
 LAR1  2.9526 0.3030 368    2.357    3.548
 LAR2a 1.5396 0.3030 368    0.944    2.135
 LAR2b 1.6880 0.3030 368    1.093    2.283

tissue = Mixed:
 Genes emmean     SE  df lower.CL upper.CL
 LAR1  3.8150 0.5240 368    2.784    4.846
 LAR2a 0.2479 0.5240 368   -0.783    1.279
 LAR2b 0.0482 0.5240 368   -0.983    1.079

tissue = Stem:
 Genes emmean     SE  df lower.CL upper.CL
 LAR1  4.6678 0.0899 368    4.491    4.845
 LAR2a 1.3405 0.0899 368    1.164    1.517
 LAR2b 3.7866 0.0899 368    3.610    3.963

tissue = Xylem:
 Genes emmean     SE  df lower.CL upper.CL
 LAR1  3.4244 0.1980 368    3.035    3.814
 LAR2a 0.2885 0.1980 368   -0.101    0.678
 LAR2b 0.2697 0.1980 368   -0.120    0.659

Degrees-of-freedom method: kenward-roger
Confidence level used: 0.95
```

```
 LAR2a - LAR2b  -2.1584 0.913 274  -2.365  0.0562

tissue = Leaf adult:
 contrast      estimate    SE  df t.ratio p.value
 LAR1 - LAR2a    4.4051 0.527 274   8.359  <.0001
 LAR1 - LAR2b    3.8456 0.527 274   7.297  <.0001
 LAR2a - LAR2b  -0.5595 0.527 274  -1.062  0.8679

tissue = Leaf and twigs:
 contrast      estimate    SE  df t.ratio p.value
 LAR1 - LAR2a    2.2187 0.263 274   8.420  <.0001
 LAR1 - LAR2b    2.3200 0.263 274   8.805  <.0001
 LAR2a - LAR2b   0.1013 0.263 274   0.385  1.0000

tissue = Leaf juvenile:
 contrast      estimate    SE  df t.ratio p.value
 LAR1 - LAR2a    3.7371 0.527 274   7.091  <.0001
 LAR1 - LAR2b    3.7979 0.527 274   7.207  <.0001
 LAR2a - LAR2b   0.0608 0.527 274   0.115  1.0000

tissue = Leaf midrib:
 contrast      estimate    SE  df t.ratio p.value
 LAR1 - LAR2a    1.4130 0.373 274   3.792  0.0006
 LAR1 - LAR2b    1.2646 0.373 274   3.394  0.0024
 LAR2a - LAR2b  -0.1484 0.373 274  -0.398  1.0000

tissue = Mixed:
 contrast      estimate    SE  df t.ratio p.value
 LAR1 - LAR2a    3.5671 0.645 274   5.527  <.0001
 LAR1 - LAR2b    3.7668 0.645 274   5.836  <.0001
 LAR2a - LAR2b   0.1997 0.645 274   0.309  1.0000

tissue = Stem:
 contrast      estimate    SE  df t.ratio p.value
 LAR1 - LAR2a    3.3273 0.111 274  30.060  <.0001
 LAR1 - LAR2b    0.8812 0.111 274   7.961  <.0001
 LAR2a - LAR2b  -2.4461 0.111 274 -22.099  <.0001

tissue = Xylem:
 contrast      estimate    SE  df t.ratio p.value
 LAR1 - LAR2a    3.1360 0.244 274  12.855  <.0001
 LAR1 - LAR2b    3.1547 0.244 274  12.932  <.0001
 LAR2a - LAR2b   0.0187 0.244 274   0.077  1.0000

Degrees-of-freedom method: kenward-roger
P value adjustment: bonferroni method for 3 tests
```

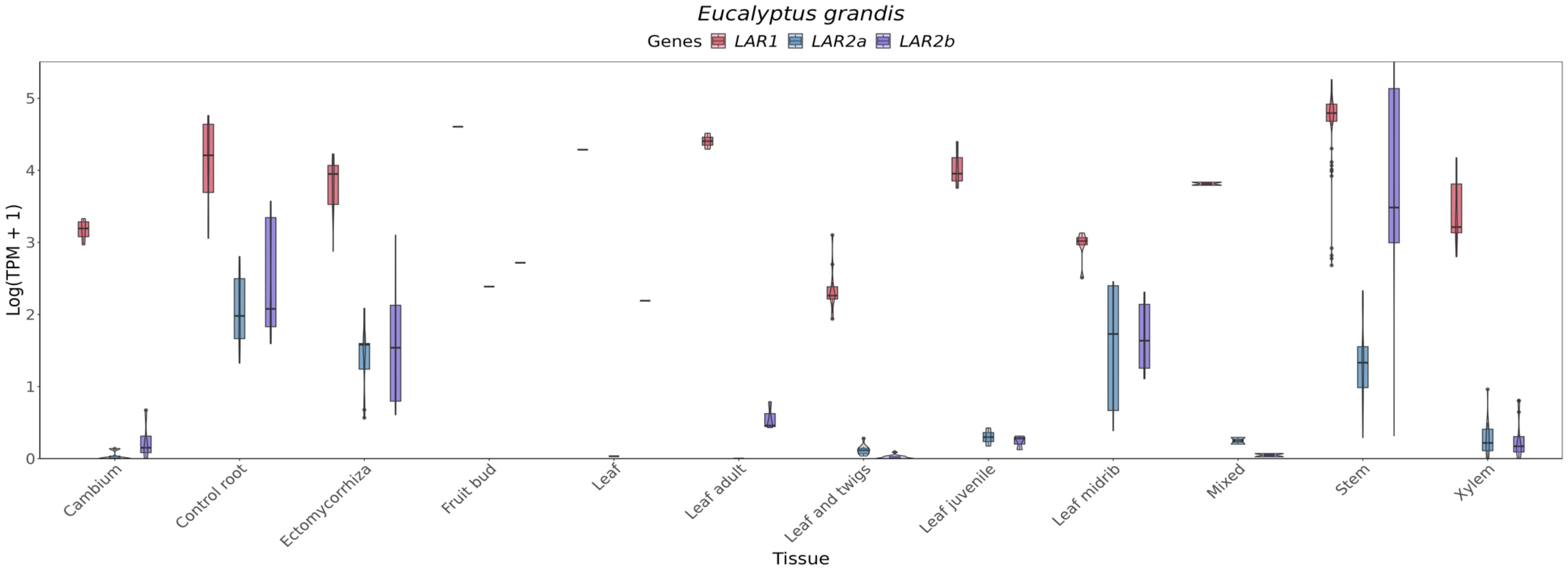

**Figure 5S.** Comparison of the expression of *LAR1a* (red), *LAR2a* (blue), and *LAR2b* (purple) genes in the species *Eucalyptus grandis*.

#### *Populus trichocarpa*

Linear mixed model fit by REML. t-tests use Satterthwaite's method ['lmerModLmerTest']
Formula: TPM_log ~ Genes * tissue + (1 | Sample)
Data: data_long

REML criterion at convergence: 7655.2

Scaled residuals:

| Min | 1Q | Median | 3Q | Max |
|---|---|---|---|---|
| -5.2516 | -0.3877 | 0.0057 | 0.3673 | 5.5306 |

Random effects:

| Groups | Name | Variance | Std.Dev. |
|---|---|---|---|
| Sample | (Intercept) | 0.5646 | 0.7514 |
| Residual | 0.1486 | 0.3855 | |
| Number of obs: | 3804 | groups: | Sample, 1902 |

Fixed effects:

| | *Estimate* | *Std. Error* | *df* | *T-value* | *Pr(>\|t\|)* | |
|---|---|---|---|---|---|---|
| *(Intercept)* | *3.78672* | *0.59715* | *2316.32522* | *6.341* | *2.73E-10* | *** |
| *GenesLAR2* | *-0.34828* | *0.38548* | *1884.00007* | *-0.904* | *0.366366* | |
| *tissueFlower* | *0.90106* | *0.6313* | *2316.32672* | *1.427* | *0.153623* | |
| *tissueInternode* | *-2.706* | *0.63838* | *2316.327* | *-4.239* | *2.33E-05* | *** |
| *tissueLate-dormant-bud* | *-3.00948* | *0.77091* | *2316.33092* | *-3.904* | *9.74E-05* | *** |
| *tissueLeaf* | *-0.36786* | *0.59823* | *2316.32527* | *-0.615* | *0.538678* | |
| *tissueMature-leaf* | *-1.5384* | *0.66017* | *2316.32781* | *-2.33* | *0.019876* | * |
| *tissueOpen-bud* | *-1.26494* | *0.84449* | *2316.33234* | *-1.498* | *0.134305* | |
| *tissuePredormant-bud* | *1.19204* | *0.70655* | *2316.32929* | *1.687* | *0.091716* | . |
| *tissueRoot* | *0.1208* | *0.59817* | *2316.32526* | *0.202* | *0.839977* | |
| *tissueSeedling* | *-1.07962* | *0.77091* | *2316.33092* | *-1.4* | *0.161514* | |
| *tissueShoot-apex* | *-2.1454* | *0.77091* | *2316.33092* | *-2.783* | *0.005431* | ** |
| *tissueStem* | *-1.63536* | *0.64499* | *2316.32725* | *-2.535* | *0.011295* | * |
| *tissueSwelling-bud* | *-0.73183* | *0.77091* | *2316.33092* | *-0.949* | *0.342569* | |
| *tissueVascular leaf* | *-0.58716* | *0.6237* | *2316.3264* | *-0.941* | *0.346588* | |
| *tissueWhole root system* | *-1.55354* | *0.64499* | *2316.32725* | *-2.409* | *0.016091* | * |
| *tissueXylem* | *-3.40956* | *0.59807* | *2316.32526* | *-5.701* | *1.34E-08* | *** |
| *tissueYoung-leaf* | *1.04921* | *0.64499* | *2316.32725* | *1.627* | *0.103938* | |
| *tissueYoung-stem* | *-0.37613* | *0.73135* | *2316.32997* | *-0.514* | *0.607095* | |
| *GenesLAR2:tissueFlower* | *-1.93239* | *0.40752* | *1884.00007* | *-4.742* | *2.28E-06* | *** |
| *GenesLAR2:tissueInternode* | *-0.60867* | *0.41209* | *1884.00007* | *-1.477* | *0.139837* | |
| *GenesLAR2:tissueLate-dormant-bud* | *0.09812* | *0.49765* | *1884.00008* | *0.197* | *0.843723* | |
| *GenesLAR2:tissueLeaf* | *-0.25688* | *0.38618* | *1884.00007* | *-0.665* | *0.506014* | |

| | | | | | | |
|---|---|---|---|---|---|---|
| *GenesLAR2:tissueMature-leaf* | *-0.52891* | *0.42616* | *1884.00007* | *-1.241* | *0.214721* | |
| *GenesLAR2:tissueOpen-bud* | *-0.63687* | *0.54514* | *1884.00009* | *-1.168* | *0.242849* | |
| *GenesLAR2:tissuePredormant-bud* | *0.1435* | *0.4561* | *1884.00008* | *0.315* | *0.75308* | |
| *GenesLAR2:tissueRoot* | *-0.55091* | *0.38614* | *1884.00007* | *-1.427* | *0.153822* | |
| *GenesLAR2:tissueSeedling* | *-1.07131* | *0.49765* | *1884.00008* | *-2.153* | *0.031464* | * |
| *GenesLAR2:tissueShoot-apex* | *-0.63473* | *0.49765* | *1884.00008* | *-1.275* | *0.202303* | |
| *GenesLAR2:tissueStem* | *-0.95* | *0.41636* | *1884.00007* | *-2.282* | *0.02262* | * |
| *GenesLAR2:tissueSwelling-bud* | *-0.77742* | *0.49765* | *1884.00008* | *-1.562* | *0.11841* | |
| *GenesLAR2:tissueVascular leaf* | *-1.44616* | *0.40262* | *1884.00007* | *-3.592* | *0.000337* | *** |
| *GenesLAR2:tissueWhole root system* | *0.80384* | *0.41636* | *1884.00007* | *1.931* | *0.053678* | . |
| *GenesLAR2:tissueXylem* | *0.20509* | *0.38607* | *1884.00007* | *0.531* | *0.595319* | |
| *GenesLAR2:tissueYoung-leaf* | *-0.2396* | *0.41636* | *1884.00007* | *-0.575* | *0.565056* | |
| *GenesLAR2:tissueYoung-stem* | *-0.81694* | *0.47211* | *1884.00008* | *-1.73* | *0.083722* | . |

Signif. codes:  0 '***' 0.001 '**' 0.01 '*' 0.05 '.' 0.1 ' ' 1

**Table 12S.** Estimated marginal means and pairwise comparison for the species *Populus trichocarpa*. Degrees-of-freedom method: asymptotic

| ***Estimated marginal means*** | ***Contrasts*** |
|---|---|
| tissue = Early-dormant-bud:<br>Genes emmean SE df asymp.LCL asymp.UCL<br>LAR1 3.787 0.5970 Inf 2.616 4.957<br>LAR2 3.438 0.5970 Inf 2.268 4.609<br><br>tissue = Flower:<br>Genes emmean SE df asymp.LCL asymp.UCL<br>LAR1 4.688 0.2050 Inf 4.286 5.089<br>LAR2 2.407 0.2050 Inf 2.006 2.809<br><br>tissue = Internode:<br>Genes emmean SE df asymp.LCL asymp.UCL<br>LAR1 1.081 0.2260 Inf 0.638 1.523<br>LAR2 0.124 0.2260 Inf -0.319 0.566<br><br>tissue = Late-dormant-bud:<br>Genes emmean SE df asymp.LCL asymp.UCL<br>LAR1 0.777 0.4880 Inf -0.178 1.733<br>LAR2 0.527 0.4880 Inf -0.429 1.483<br><br>tissue = Leaf:<br>Genes emmean SE df asymp.LCL asymp.UCL<br>LAR1 3.419 0.0360 Inf 3.348 3.490<br>LAR2 2.814 0.0360 Inf 2.743 2.884<br><br>tissue = Mature-leaf:<br>Genes emmean SE df asymp.LCL asymp.UCL<br>LAR1 2.248 0.2810 Inf 1.697 2.800<br>LAR2 1.371 0.2810 Inf 0.819 1.923 | tissue = Early-dormant-bud:<br>contrast estimate SE df z.ratio p.value<br>LAR1 - LAR2 0.348 0.3850 Inf 0.904 0.3663<br><br>tissue = Flower:<br>contrast estimate SE df z.ratio p.value<br>LAR1 - LAR2 2.281 0.1320 Inf 17.249 <.0001<br><br>tissue = Internode:<br>contrast estimate SE df z.ratio p.value<br>LAR1 - LAR2 0.957 0.1460 Inf 6.568 <.0001<br><br>tissue = Late-dormant-bud:<br>contrast estimate SE df z.ratio p.value<br>LAR1 - LAR2 0.250 0.3150 Inf 0.795 0.4267<br><br>tissue = Leaf:<br>contrast estimate SE df z.ratio p.value<br>LAR1 - LAR2 0.605 0.0233 Inf 26.010 <.0001<br><br>tissue = Mature-leaf:<br>contrast estimate SE df z.ratio p.value<br>LAR1 - LAR2 0.877 0.1820 Inf 4.827 <.0001<br><br>tissue = Open-bud:<br>contrast estimate SE df z.ratio p.value<br>LAR1 - LAR2 0.985 0.3850 Inf 2.556 0.0106<br><br>tissue = Predormant-bud:<br>contrast estimate SE df z.ratio p.value |

```
tissue = Open-bud:
 Genes emmean     SE  df asymp.LCL asymp.UCL
 LAR1   2.522 0.5970 Inf     1.351     3.692
 LAR2   1.537 0.5970 Inf     0.366     2.707

tissue = Predormant-bud:
 Genes emmean     SE  df asymp.LCL asymp.UCL
 LAR1   4.979 0.3780 Inf     4.239     5.719
 LAR2   4.774 0.3780 Inf     4.034     5.514

tissue = Root:
 Genes emmean     SE  df asymp.LCL asymp.UCL
 LAR1   3.908 0.0349 Inf     3.839     3.976
 LAR2   3.008 0.0349 Inf     2.940     3.077

tissue = Seedling:
 Genes emmean     SE  df asymp.LCL asymp.UCL
 LAR1   2.707 0.4880 Inf     1.751     3.663
 LAR2   1.288 0.4880 Inf     0.332     2.243

tissue = Shoot-apex:
 Genes emmean     SE  df asymp.LCL asymp.UCL
 LAR1   1.641 0.4880 Inf     0.686     2.597
 LAR2   0.658 0.4880 Inf    -0.297     1.614

tissue = Stem:
 Genes emmean     SE  df asymp.LCL asymp.UCL
 LAR1   2.151 0.2440 Inf     1.674     2.629
 LAR2   0.853 0.2440 Inf     0.375     1.331

tissue = Swelling-bud:
 Genes emmean     SE  df asymp.LCL asymp.UCL
 LAR1   3.055 0.4880 Inf     2.099     4.011
 LAR2   1.929 0.4880 Inf     0.974     2.885

tissue = Vascular leaf:
 Genes emmean     SE  df asymp.LCL asymp.UCL
 LAR1   3.200 0.1800 Inf     2.847     3.552
 LAR2   1.405 0.1800 Inf     1.052     1.758

tissue = Whole root system:
 Genes emmean     SE  df asymp.LCL asymp.UCL
 LAR1   2.233 0.2440 Inf     1.755     2.711
 LAR2   2.689 0.2440 Inf     2.211     3.167
tissue = Xylem:
 Genes emmean     SE  df asymp.LCL asymp.UCL
 LAR1   0.377 0.0332 Inf     0.312     0.442
 LAR2   0.234 0.0332 Inf     0.169     0.299
```

```
 LAR1 - LAR2    0.205 0.2440 Inf   0.840  0.4009

tissue = Root:
 contrast    estimate     SE  df z.ratio p.value
 LAR1 - LAR2    0.899 0.0226 Inf  39.861  <.0001

tissue = Seedling:
 contrast    estimate     SE  df z.ratio p.value
 LAR1 - LAR2    1.420 0.3150 Inf   4.510  <.0001

tissue = Shoot-apex:
 contrast    estimate     SE  df z.ratio p.value
 LAR1 - LAR2    0.983 0.3150 Inf   3.123  0.0018

tissue = Stem:
 contrast    estimate     SE  df z.ratio p.value
 LAR1 - LAR2    1.298 0.1570 Inf   8.250  <.0001

tissue = Swelling-bud:
 contrast    estimate     SE  df z.ratio p.value
 LAR1 - LAR2    1.126 0.3150 Inf   3.577  0.0003

tissue = Vascular leaf:
 contrast    estimate     SE  df z.ratio p.value
 LAR1 - LAR2    1.794 0.1160 Inf  15.439  <.0001

tissue = Whole root system:
 contrast    estimate     SE  df z.ratio p.value
 LAR1 - LAR2   -0.456 0.1570 Inf  -2.895  0.0038

tissue = Xylem:
 contrast    estimate     SE  df z.ratio p.value
 LAR1 - LAR2    0.143 0.0214 Inf   6.676  <.0001

tissue = Young-leaf:
 contrast    estimate     SE  df z.ratio p.value
 LAR1 - LAR2    0.588 0.1570 Inf   3.736  0.0002

tissue = Young-stem:
 contrast    estimate     SE  df z.ratio p.value
 LAR1 - LAR2    1.165 0.2730 Inf   4.275  <.0001

Degrees-of-freedom method: asymptotic
```

| |
|---|
| tissue = Young-leaf:<br> Genes emmean SE df asymp.LCL asymp.UCL<br> LAR1 4.836 0.2440 Inf 4.358 5.314<br> LAR2 4.248 0.2440 Inf 3.770 4.726<br>tissue = Young-stem:<br> Genes emmean SE df asymp.LCL asymp.UCL<br> LAR1 3.411 0.4220 Inf 2.583 4.238<br> LAR2 2.245 0.4220 Inf 1.418 3.073<br>Degrees-of-freedom method: asymptotic<br>Confidence level used: 0.95 |

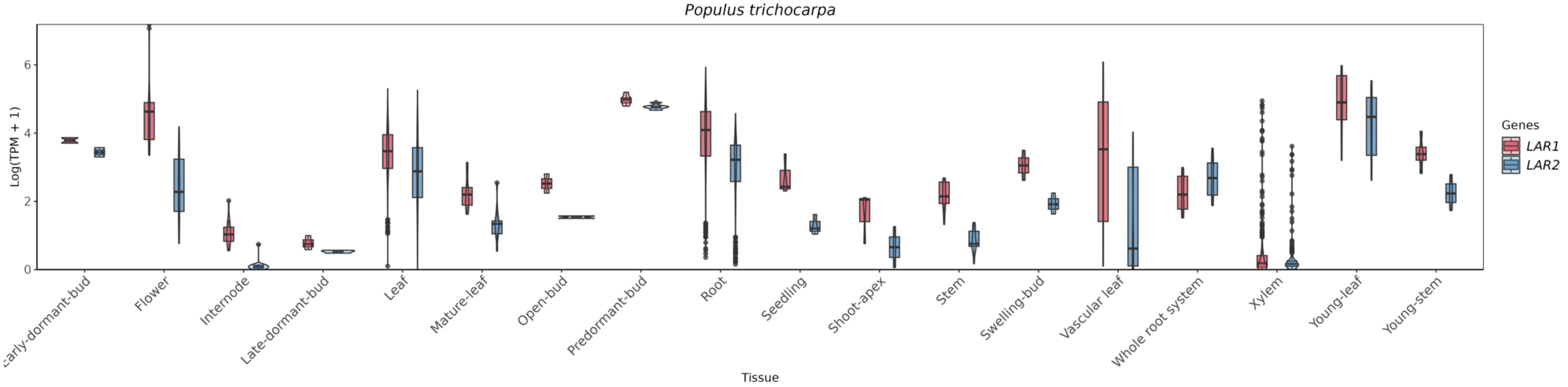

**Figure 6S.** Comparison of the expression of *LAR1* (red), and *LAR2* (blue) genes in the species *Populus trichocarpa.*

#### *Carya illinoinensis*

Linear mixed model fit by REML. t-tests use Satterthwaite's method ['lmerModLmerTest']
Formula: TPM_log ~ Genes * tissue + (1 | Sample)
Data: data_long

REML criterion at convergence: 7655.2

Scaled residuals:

| Min | 1Q | Median | 3Q | Max |
|---|---|---|---|---|
| -2.0486 | -0.7132 | -0.0787 | 0.5695 | 3.6612 |

Random effects:

| Groups | Name | Variance | Std.Dev. |
|---|---|---|---|
| Sample | (Intercept) | 0.002874 | 0.05361 |
| Residual | 0.438219 | 0.66198 | |
| Number of obs: | 178 | groups: | Sample, 89 |

Fixed effects:

| | Estimate | Std. Error | df | T-value | Pr(>\|t\|) | |
|---|---|---|---|---|---|---|
| (Intercept) | 3.4887 | 0.1917 | 163.993 | 18.197 | <2E-016 | *** |
| GenesLAR2 | -0.7197 | 0.2703 | 82 | -2.663 | 0.009317 | ** |
| tissuebud | -1.8182 | 0.3321 | 163.993 | -5.475 | 1.61E-07 | *** |
| tissueembryo | -0.8942 | 0.2572 | 163.993 | -3.476 | 0.000651 | *** |
| tissueflower | -1.9189 | 0.3834 | 163.993 | -5.004 | 1.43E-06 | *** |
| tissueKernel | -1.4147 | 0.3834 | 163.993 | -3.689 | 0.000305 | *** |
| tissueLeaf | -0.5105 | 0.2475 | 163.993 | -2.062 | 0.040745 | * |
| tissueseedling | -2.0248 | 0.2268 | 163.993 | -8.926 | 8.53E-16 | *** |
| GenesLAR2:tissuebud | 1.6143 | 0.4681 | 82 | 3.449 | 0.000892 | *** |
| GenesLAR2:tissueembryo | -1.374 | 0.3626 | 82 | -3.789 | 0.000287 | *** |
| GenesLAR2:tissueflower | 2.5682 | 0.5405 | 82 | 4.751 | 8.49E-06 | *** |
| GenesLAR2:tissueKernel | -0.7166 | 0.5405 | 82 | -1.326 | 0.188606 | |
| GenesLAR2:tissueLeaf | -0.9122 | 0.3489 | 82 | -2.615 | 0.01063 | * |
| enesLAR2:tissueseedling | 0.791 | 0.3198 | 82 | 2.474 | 0.015432 | * |

Signif. codes: 0 ‘***’ 0.001 ‘**’ 0.01 ‘*’ 0.05 ‘.’ 0.1 ‘ ’ 1

**Table 13S.** Estimated marginal means and pairwise comparison for the species *Carya illinoinensis*. Degrees-of-freedom method: kenward-roger. Confidence level used: 0.95

| ***Estimated Marginal Means*** | ***Contrasts*** |
|---|---|
| tissue = Branch:<br>Genes emmean SE df lower.CL upper.CL<br>LAR1 3.489 0.192 164 3.1102 3.867<br>LAR2 2.769 0.192 164 2.3905 3.148<br><br>tissue = bud:<br>Genes emmean SE df lower.CL upper.CL<br>LAR1 1.671 0.271 164 1.1351 2.206<br>LAR2 2.565 0.271 164 2.0297 3.100<br><br>tissue = embryo:<br>Genes emmean SE df lower.CL upper.CL<br>LAR1 2.595 0.171 164 2.2559 2.933<br>LAR2 0.501 0.171 164 0.1622 0.839<br><br>tissue = flower:<br>Genes emmean SE df lower.CL upper.CL<br>LAR1 1.570 0.332 164 0.9142 2.226<br>LAR2 3.418 0.332 164 2.7626 4.074<br><br>tissue = Kernel:<br>Genes emmean SE df lower.CL upper.CL<br>LAR1 2.074 0.332 164 1.4184 2.730<br>LAR2 0.638 0.332 164 -0.0179 1.293<br><br>tissue = Leaf:<br>Genes emmean SE df lower.CL upper.CL<br>LAR1 2.978 0.157 164 2.6692 3.287<br>LAR2 1.346 0.157 164 1.0373 1.655<br>tissue = seedling:<br>Genes emmean SE df lower.CL upper.CL<br>LAR1 1.464 0.121 164 1.2245 1.703<br>LAR2 1.535 0.121 164 1.2959 1.775 | tissue = Branch:<br>contrast estimate SE df t.ratio p.value<br>LAR1 - LAR2 0.7197 0.270 82 2.663 0.0093<br><br>tissue = bud:<br>contrast estimate SE df t.ratio p.value<br>LAR1 - LAR2 -0.8946 0.382 82 -2.341 0.0217<br><br>tissue = embryo:<br>contrast estimate SE df t.ratio p.value<br>LAR1 - LAR2 2.0937 0.242 82 8.662 <.0001<br><br>tissue = flower:<br>contrast estimate SE df t.ratio p.value<br>LAR1 - LAR2 -1.8485 0.468 82 -3.949 0.0002<br><br>tissue = Kernel:<br>contrast estimate SE df t.ratio p.value<br>LAR1 - LAR2 1.4363 0.468 82 3.068 0.0029<br><br>tissue = Leaf:<br>contrast estimate SE df t.ratio p.value<br>LAR1 - LAR2 1.6319 0.221 82 7.396 <.0001<br><br>tissue = seedling:<br>contrast estimate SE df t.ratio p.value<br>LAR1 - LAR2 -0.0713 0.171 82 -0.417 0.6775<br><br>Degrees-of-freedom method: kenward-roger<br>Confidence level used: 0.95 |

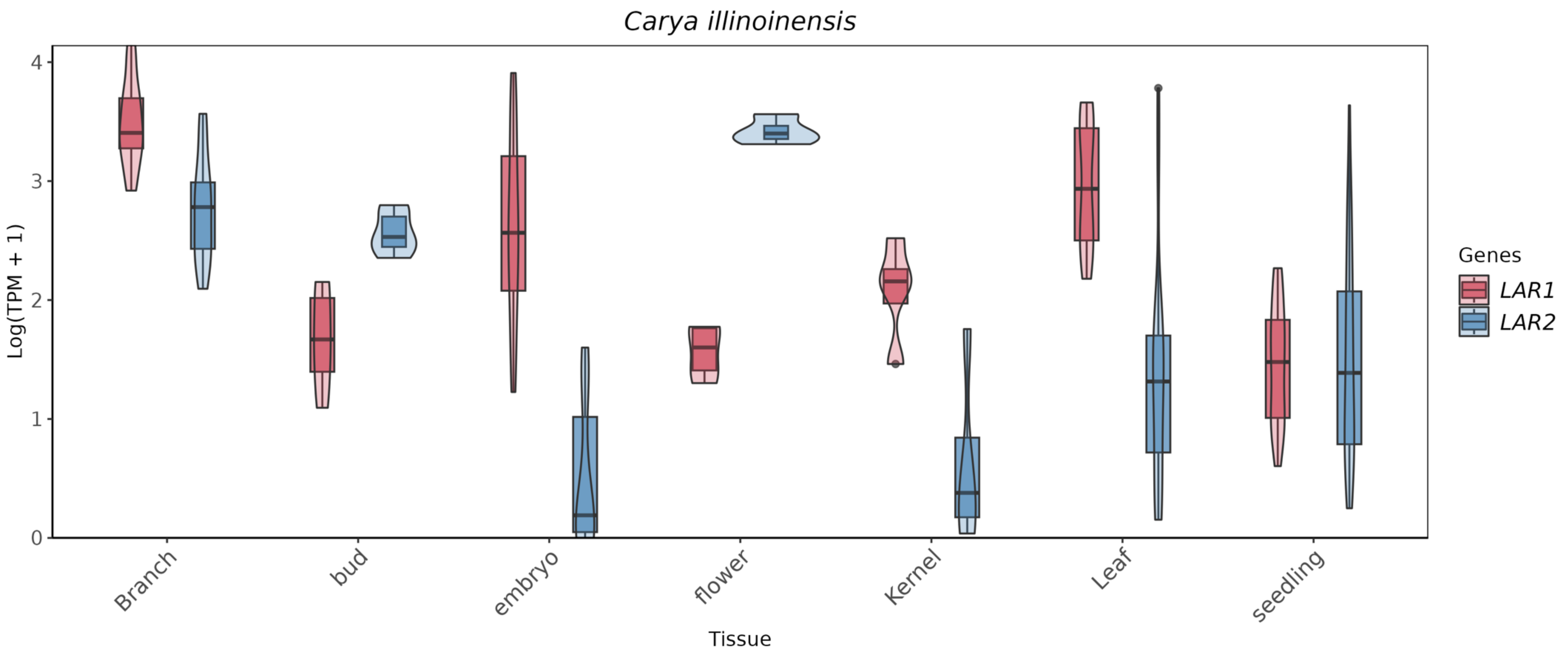

**Figure 7S.** Comparison of the expression of *LAR1* (red), and *LAR2* (blue) genes in the species *Carya illinoinensis*.

#### *Morella rubra*

Linear mixed model fit by REML. t-tests use Satterthwaite's method ['lmerModLmerTest']
Formula: TPM_log ~ Genes * tissue + (1 | Sample)
Data: data_long

REML criterion at convergence: 638.3

Scaled residuals:

| Min | 1Q | Median | 3Q | Max |
|---|---|---|---|---|
| -2.5316 | -0.5276 | -0.1306 | 0.4559 | 5.5753 |

Random effects:

| Groups | Name | Variance | Std.Dev. |
|---|---|---|---|
| Sample | (Intercept) | 0.1621 | 0.4026 |
| Residual | 0.4913 | 0.7009 | |
| Number of obs: | 272 | groups: | Sample, 136 |

Fixed effects:

| | *Estimate* | *Std. Error* | *df* | *T-value* | *Pr(>\|t\|)* | |
|---|---|---|---|---|---|---|
| *(Intercept)* | 4.86342 | 0.80833 | 244.92727 | 6.017 | 6.45E-09 | *** |
| *GenesLAR2* | -0.38874 | 0.99127 | 130 | -0.392 | 0.6956 | |
| *tissueflower* | 0.09484 | 0.88549 | 244.92727 | 0.107 | 0.9148 | |
| *tissuefruits* | -2.07341 | 0.81262 | 244.92727 | -2.552 | 0.0113 | * |
| *tissueleaves* | 0.29296 | 0.82014 | 244.92727 | 0.357 | 0.7212 | |
| *tissueroot* | 0.32799 | 1.14316 | 244.92727 | 0.287 | 0.7744 | |
| *tissuestem* | -0.52627 | 1.14316 | 244.92727 | -0.46 | 0.6457 | |
| *GenesLAR2:tissueflower* | -0.47636 | 1.08589 | 130 | -0.439 | 0.6616 | |
| *GenesLAR2:tissuefruits* | -1.62229 | 0.99653 | 130 | -1.628 | 0.106 | |
| *GenesLAR2:tissueleaves* | -1.03024 | 1.00575 | 130 | -1.024 | 0.3076 | |
| *GenesLAR2:tissueroot* | -0.61444 | 1.40187 | 130 | -0.438 | 0.6619 | |
| *GenesLAR2:tissuestem* | 1.19982 | 1.40187 | 130 | 0.856 | 0.3936 | |

Signif. codes: 0 ‘***’ 0.001 ‘**’ 0.01 ‘*’ 0.05 ‘.’ 0.1 ‘ ’ 1

**Table 14S.** Estimated marginal means and pairwise comparison for the species *Morella rubra*. Degrees-of-freedom method: kenward-roger. Confidence level used: 0.95

| ***Estimated Marginal Means*** | ***Contrasts*** |
|---|---|
| tissue = bud:<br>Genes emmean SE df lower.CL upper.CL<br>LAR1 4.863 0.8080 245 3.271 6.456<br>LAR2 4.475 0.8080 245 2.883 6.067<br><br>tissue = flower:<br>Genes emmean SE df lower.CL upper.CL<br>LAR1 4.958 0.3610 245 4.246 5.670<br>LAR2 4.093 0.3610 245 3.381 4.805<br><br>tissue = fruits:<br>Genes emmean SE df lower.CL upper.CL<br>LAR1 2.790 0.0834 245 2.626 2.954<br>LAR2 0.779 0.0834 245 0.615 0.943<br><br>tissue = leaves:<br>Genes emmean SE df lower.CL upper.CL<br>LAR1 5.156 0.1390 245 4.883 5.429<br>LAR2 3.737 0.1390 245 3.464 4.010<br><br>tissue = root:<br>Genes emmean SE df lower.CL upper.CL<br>LAR1 5.191 0.8080 245 3.599 6.784<br>LAR2 4.188 0.8080 245 2.596 5.780<br><br>tissue = stem:<br>Genes emmean SE df lower.CL upper.CL<br>LAR1 4.337 0.8080 245 2.745 5.929<br>LAR2 5.148 0.8080 245 3.556 6.740<br><br>Degrees-of-freedom method: kenward-roger<br>Confidence level used: 0.95 | tissue = bud:<br>contrast estimate SE df t.ratio p.value<br>LAR1 - LAR2 0.389 0.991 130 0.392 0.6956<br><br>tissue = flower:<br>contrast estimate SE df t.ratio p.value<br>LAR1 - LAR2 0.865 0.443 130 1.951 0.0532<br><br>tissue = fruits:<br>contrast estimate SE df t.ratio p.value<br>LAR1 - LAR2 2.011 0.102 130 19.669 <.0001<br><br>tissue = leaves:<br>contrast estimate SE df t.ratio p.value<br>LAR1 - LAR2 1.419 0.170 130 8.347 <.0001<br><br>tissue = root:<br>contrast estimate SE df t.ratio p.value<br>LAR1 - LAR2 1.003 0.991 130 1.012 0.3134<br><br>tissue = stem:<br>contrast estimate SE df t.ratio p.value<br>LAR1 - LAR2 -0.811 0.991 130 -0.818 0.4147<br><br>Degrees-of-freedom method: kenward-roger |

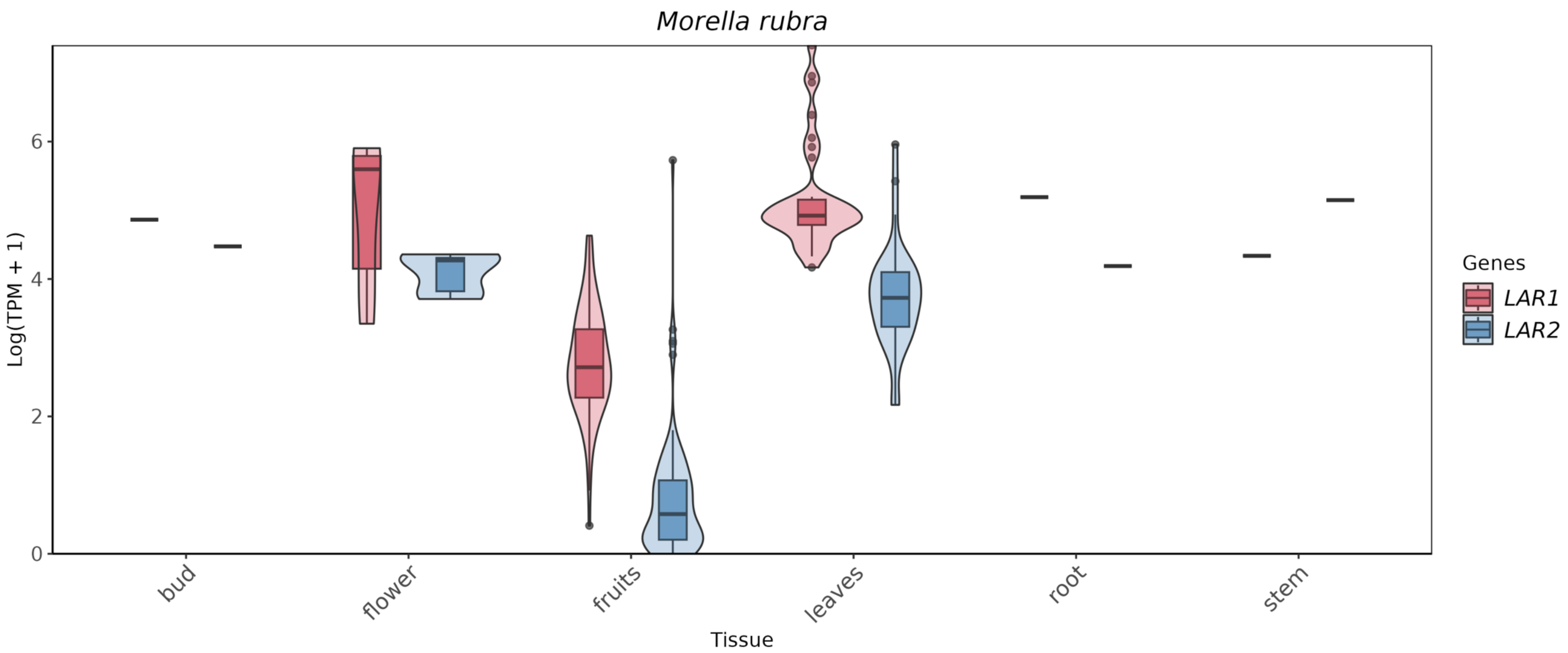

**Figure 8S.** Comparison of the expression of *LAR1* (red), and *LAR2* (blue) genes in the species *Morella rubra*.
